# Molecular phylogeny of historical micro-invertebrate specimens using *de novo* sequence assembly

**DOI:** 10.1101/2020.03.30.015669

**Authors:** R.J.S Orr, M. M. Sannum, S. Boessenkool, E. Di Martino, D.P. Gordon, H. Mello, M. Obst, M.H. Ramsfjell, A.M. Smith, L.H. Liow

## Abstract

Resolution of relationships at lower taxonomic levels is crucial for answering many evolutionary questions, and as such, sufficiently varied species representation is vital. This latter goal is not always achievable with relatively fresh samples. To alleviate the difficulties in procuring rarer taxa, we have seen increasing utilization of historical specimens in building molecular phylogenies using high throughput sequencing. This effort, however, has mainly focused on large-bodied or well-studied groups, with small-bodied and under-studied taxa under-prioritized. Here, we present a pipeline that utilizes both historical and contemporary specimens, to increase the resolution of phylogenetic relationships among understudied and small-bodied metazoans, namely, cheilostome bryozoans. In this study, we pioneer sequencing of air-dried bryozoans, utilizing a recent library preparation method for low DNA input. We use the *de novo* mitogenome assembly from the target specimen itself as reference for iterative mapping, and the comparison thereof. In doing so, we present mitochondrial and ribosomal RNA sequences of 43 cheilostomes representing 37 species, including 14 from historical samples ranging from 50 to 149 years old. The inferred phylogenetic relationships of these samples, analyzed together with publicly available sequence data, are shown in a statistically well-supported 65 taxa and 17 genes cheilostome tree. Finally, the methodological success is emphasized by circularizing a total of 27 mitogenomes, seven from historical cheilostome samples. Our study highlights the potential of utilizing DNA from micro-invertebrate specimens stored in natural history collections for resolving phylogenetic relationships between species.

## Introduction

Robust phylogenetic hypotheses are crucial to understanding many biological processes, ranging from those contributing to population history to those creating macroevolutionary patterns. The development of methods for phylogenetic estimation and high throughput sequencing (HTS) have, in combination, improved our understanding of the relationships among extant organisms. These advances are frequently applied to the estimation of relationships among higher taxa, e.g. distantly related animal phyla (Laumer et al., 2019; Simion et al., 2017), families of flowering plants (Léveillé-Bourret, Starr, Ford, Moriarty Lemmon, & Lemmon, 2017) and orders of birds (Jarvis et al., 2014). While these deeper branches have received significant attention, those at the leaves (e.g. genera and species) often remaining largely unresolved, especially for taxa that are understudied. For the resolution of relationships at lower taxonomic levels, crucial as a backbone for answering many evolutionary questions, a rich and broad species representation is vital.

Traditionally, researchers have sequenced relatively freshly collected specimens. However, there is an increasing utilization of historical specimens for molecular phylogenetic reconstruction in the absence of fresh tissue (e.g. Jarvis et al 2014). Historical specimens from museum or other institutional collections are valuable sources of information representing organisms that may be difficult or impossible to sample in contemporary populations (Holmes et al., 2016). Most studies that use historical specimens for phylogenetic reconstruction tend to focus on larger-bodied species that are recently extinct (Anmarkrud & Lifjeld, 2017; Bunce et al., 2009; Mitchell et al., 2014; Sharko et al., 2019), well-studied groups (Billerman & Walsh, 2019; Jarvis et al., 2014) and organisms of economic importance (Bonanomi, Therkildsen, Hedeholm, Hemmer-Hansen, & Nielsen, 2014; Larson et al., 2007; Larsson et al., 2019), although there are a few reports on invertebrate and other small-bodied taxa (Der Sarkissian et al., 2017; Kistenich et al., 2019; Sproul & Maddison, 2017). When employing HTS in such historical studies, nucleotide reads are typically mapped to a closely related reference genome, or one from the same species (Sharko et al., 2019). Alternatively, complete sequences of target regions from related species and/or genera (Anmarkrud & Lifjeld, 2017; Billerman & Walsh, 2019) are used to design baits or probes (Derkarabetian, Benavides, & Giribet, 2019; Ruane & Austin, 2017). While these approaches are excellent for groups whose sequences are relatively well-understood, they are unfeasible for clades with low inter-genus sequence conservation, or for those lacking sequence data from closely related reference organisms.

Here, we present a pipeline to use historical material from lesser-studied, small-bodied organisms for the purpose of reconstructing molecular phylogenetic relationships. Our target organisms are cheilostomes, the most species-rich order of the phylum Bryozoa, with ca. 6500 described extant species, representing about 80% of the living species diversity of the phylum (Bock & Gordon, 2013). Cheilostomes are lightly- to heavily-calcified, sessile, colonial metazoans common in benthic marine habitats. Most species are encrusting, while fewer are erect, and most are small (colony size c.1 cm^2^, module size c. 500 μm x 200 μm), and live on hard substrates that may be overgrown by other fouling organisms (including other bryozoans, hydroids, foraminiferans and tube worms).

Systematic relationships among cheilostome bryozoans remain largely based on morphological characters (Bock & Gordon, 2013) with molecular phylogenies being restricted to recently collected, ethanol-preserved samples where genetic data was obtained using PCR-based methods (Fuchs, Obst, & Sundberg, 2009; Knight, Gordon, & Lavery, 2011; Orr, Waeschenbach, et al., 2019; Waeschenbach, Taylor, & Littlewood, 2012), or more recently, by a HTS genome-skimming approach (Orr, Haugen, et al., 2019). While these studies have improved our understanding of the phylogenetic relationships among cheilostomes, many key taxa, potentially filling important phylogenetic positions, are hard to procure and remain unfeatured in sequencing projects. Contributing to the advancement of historical DNA methods (Billerman & Walsh, 2019; Knapp & Hofreiter, 2010), we pioneer the sequencing of air-dried bryozoan specimens (i.e. never preserved in ethanol) that have been stored up to 150 years since collection. To do so, we employ a recently developed DNA library preparation method (SMARTer ThruPLEX, Takara) to amplify and sequence historical samples with low DNA concentrations. We bypass the need for primers/probes, a clear advantage for cheilostomes which are known to have low inter-genus sequence conservation (Orr, Haugen, et al., 2019; Orr, Waeschenbach, et al., 2019; Waeschenbach et al., 2012). The *de novo* assembled sequences from the target colony itself were used as a reference for iterative mapping, avoiding assumptions about what could be justifiably used as a reference. In doing so, we generate mitochondrial and ribosomal RNA sequences of 43 cheilostome colonies representing 37 species and 30 genera, with 14 of these from historical samples ranging from 50 to 149 years old. As a derivative of the demonstration of our pipeline, we also present a well-supported cheilostome tree using 65 taxa and 17 genes. Finally, the success of the methodology we employed is highlighted by the circularization of seven historical cheilostome mitochondrial genomes. This approach emphasizes the potential for analyzing DNA from micro-invertebrate samples stored in natural history collections, especially for phylogenetic reconstruction of many hitherto inaccessible cheilostome genomes.

## Materials and Methods

Twenty-one dried historical cheilostome samples and 29 recently collected samples, of which 26 were ethanol-preserved and 3 dried, were targeted (Table S1). We selected the historical samples to represent a spread of collection dates while including those that are phylogenetically previously resolved and those that are currently enigmatic. Recently collected samples were selected for their potential for phylogenetic verification of the historical specimens. Each colony was subsampled for DNA isolation and scanning electron microscopy (SEM), using a Hitachi TM4040PLus. For microscopy, we bleached subsamples in diluted household bleach for a few hours to overnight, removing soft tissues in order to document skeletal morphology. SEMs of dried samples were taken both pre- and post-bleaching. All physical vouchers are stored at the Natural History Museum of Oslo, University of Oslo, and SEMs are available in the Online Supplementary Information (SI). Other metadata of the samples are reported in Table S1.

### DNA isolation, sequencing, assembly and annotation

Genomic DNA for all 50 specimens was isolated using the DNeasy Blood and Tissue kit (QIAGEN, Germantown, MD, USA). Colonies were homogenized in lysis buffer, using a pestle, in the presence of proteinase-K prior to a 24-hour 37°C incubation period. DNA was eluted in pre-heated 65°C Tris-Cl buffer (10 mM) and incubated at 37°C for 10 minutes. Recovered DNA was quantified prior to library preparation using a Qubit 2.0 fluorometer (ThermoFisher, US).

DNA from the historical specimens were isolated in a laboratory designed for handling samples with low DNA concentrations (sensi-lab, NHM, Oslo). DNA extractions were performed inside a hood equipped with UV lights and all equipment was bleached and UV-sterilized prior to use. Samples were vortexed twice in nuclease-free H_2_O, air-dried, then subject to UV for 10 minutes to minimize surface contaminants. These treated samples were subsequently crushed with a stainless-steel mortar and pestle (Gondek, Boessenkool, & Star, 2018) prior to DNA isolation.

Isolated DNA samples with >15ng were prepared with the standard KAPA HyperPrep kit (Roche, USA) by the Norwegian Sequencing Centre (NCS, Oslo, Norway), while those with <15ng were prepared using the SMARTer® ThruPLEX® DNA-Seq Kit (Takara Bio Inc, Japan) in the sensi-lab at NHM Oslo (Table S1). Genomic DNA up to 150 bp in read length was pair-end (PE) sequenced on an Illumina HiSeq4000 at the NCS. The non-historical DNA samples were fragmented to a 350 bp insert size, whilst no such step was performed on the historical samples with already short DNA length, giving a variable insert size (Table S1). A blank control taken during the DNA extraction of the historical samples was also sequenced (library preparation: SMARTer® ThruPLEX® DNA-Seq Kit).

Illumina HiSeq reads were quality checked using FastQC v.0.11.8 (Andrews, 2010), then quality- and adapter-trimmed using TrimGalore v0.4.4 (Krueger, 2015). All samples were *de novo* assembled with SPAdes 3.11.1 (Bankevich et al., 2012) using k-mers of 21,33, 55, 77, 99, and 127. The mitogenome and rRNA operon of each sample were identified with blastn (Altschul, Gish, Miller, Myers, & Lipman, 1990) against our own database of unpublished cheilostome sequences. For each historical sample, the SPAdes *de novo* assembled mitogenome sequence was used as the reference (seed) input for its own iterative mapped assembly using GetOrganelle (Jin et al., 2019) and NOVOplasty 3.7 (Dierckxsens, Mardulyn, & Smits, 2016), both under default settings. In addition, a mitogenome sequence from a taxon phylogenetically related to the historical sample in question was also provided for a second iterative assembly (Fig S1).

### Annotation and alignments

Mitogenomes from the separate assembly methods, for each of the samples were annotated with Mitos2 using a metazoan reference and invertebrate genetic code (Bernt et al., 2013) to identify two rRNA genes (rrnL and rrnS) and 13 protein coding genes (*atp6, atp8, cox1, cox2, cox3, cob, nad1, nad2, nad3, nad4, nad4l, nad5*, and *nad6*). In addition, two rRNA operon genes (ssu (18s) and lsu (28s)) were identified and annotated using RNAmmer (Lagesen et al., 2007). Further, mitogenes and rRNA operons from 27 bryozoan taxa (Table S2), obtained from NCBI, were aligned with our samples to compile a broader taxon sample representing a cheilostome ingroup and ctenostome outgroup. A minimum gene number of four per taxon, to avoid possible effects of missing-data on phylogenetic inference (Philippe et al., 2004; Wiens & Morrill, 2011), was set for this study (see Table 1 and S1 for the number of genes included for each taxon). MAFFT (Katoh & Standley, 2013) was used for alignment with default parameters: for the four rRNA genes (nucleotide) the Q-INS-i model, considering secondary RNA structure, was utilized; for the 13 protein-coding genes, in amino acid format, the G-INS-I model was used. The 17 separate alignments were edited manually using Mesquite v3.1 (Maddison & Maddison, 2017). Ambiguously aligned characters were removed from each alignment using Gblocks (Talavera & Castresana, 2007) with least stringent parameters. The single-gene alignments were concatenated using the catfasta2phyml perl script (Nylander, 2010). An initial mitochondrial supermatrix, consisting of up to five assemblies per sample (Fig. S1) was used to investigate possible differences among assembly methods using a phylogenetic analysis (see next section). Using the best combination of assembled genes from this initial supermatrix of historical samples, we then created a downstream final supermatrix, where each sample (now including all samples in Table 1) was represented only once in the final phylogenetic analysis. Here, the two rRNA operon genes were also included (Fig. 1). The alignments (both masked and unmasked) are available through Dryad (https://doi.org/???/dryad.???).

**Table 1.** Samples generated and analyzed in this study: BLEED stands for Bryozoan Lab for Ecology, Evolution and Development and BLEED numbers are numerical tags for the specimens. Abbreviations for countries: NZ = New Zealand, SWE = Sweden, AUS = Australia, NOR = Norway, IT = Italy and UK = United Kingdom. The size of the mitogenome, in base pairs (bp), is only shown if closed/circularized, otherwise the cell is labelled as “NO”. The columns “rRNA” and “Mt” represent the number of rRNA and mitochondrial genes annotated and used in phylogenetic inference, with a maximum of 2 and 15, respectively. An * succeeding the year of collection indicates an air-dried historical sample, defined here as 50 years or older. Multiple historical samples have “<1970” indicated because the collection year was unstated. However, taxonomic identifications were made in 1970 for these specimens, implying that they must have been collected then, or earlier. For an expanded overview of metadata see Table S1 and for additional taxa used, but not generated during this study, Table S2.

| Genus | Species | BLEED | Year | Country | rRNA | Mt | Mt size (bp) | Accession |
| --- | --- | --- | --- | --- | --- | --- | --- | --- |
| *Antarctothoa* | *delta* | 703 | 2018 | NZ | 2 | 12 | 14114 | ?? |
| *Arachnopusia* | *unicornis* | 183 | 2009 | NZ | 2 | 15 | NO | ?? |
| *Arachnopusia* | *unicornis* | 221 | 2011 | NZ | 2 | 15 | NO | ?? |
| *Bicellariella* | *ciliata* | 560 | 2018 | SWE | 2 | 15 | 19704 | ?? |
| *Bugula* | *neritina* | 1200 | 1960* | AUS | 2 | 15 | 15414 | ?? |
| *Caberea* | *angusta* | 160 | 2009 | NZ | 2 | 13 | NO | ?? |
| *Caberea* | *ellisii* | 1190 | <1970* | NOR | 0 | 14 | 17589 | ?? |
| *Calpensia* | *nobilis* | 1184 | <1970* | IT | 2 | 14 | 14049 | ?? |
| *Celleporina* | *sinuata* | 328 | 2015 | NZ | 1 | 14 | 15651 | ?? |
| *Cornuticella* | *trapezoidea* | 1687 | 2018 | NZ | 2 | 14 | 14590 | ?? |
| *Cradoscrupocellaria* | *reptans* | 1192 | <1970* | NO | 1 | 15 | NO | ?? |
| *Electra* | *pilosa* | 819 | 2018 | NO | 2 | 14 | 13970 | ?? |
| *Electra* | *pilosa* | 1172 | 2018 | NO | 2 | 13 | 13965 | ?? |
| *Electra* | *pilosa* | 1178 | <1970* | UNKNOWN | 0 | 4 | NO | ?? |
| *Electra* | *scuticifera* | 98 | 2015 | NZ | 2 | 12 | 13804 | ?? |
| *Emma* | *tricellata* | 196 | 2015 | NZ | 2 | 14 | 16275 | ?? |
| *Flustra* | *foliacea* | 568 | 2018 | SWE | 2 | 15 | 16802 | ?? |
| *Hincksina* | sp. | 61 | 2009 | NZ | 2 | 14 | 15726 | ?? |
| *Hippothoa* | sp. | 1187 | 1875* | UK | 0 | 15 | NO | ?? |
| *Membranipora* | *membranacea* | 816 | 2018 | NO | 2 | 15 | 14739 | ?? |
| *Membranipora* | *membranacea* | 1163 | 2018 | SWE | 2 | 15 | 14736 | ?? |
| *Myriapora* | *truncata* | 1197 | 1871* | IT | 2 | 15 | NO | ?? |
| *Omalosecosa* | *ramulosa* | 1177 | <1970* | NO | 1 | 8 | NO | ?? |
| *Oshurkovia* | *littoralis* | 1194 | <1970* | UK | 2 | 13 | 13973 | ?? |
| *Parasmittina* | *aotea* | 86 | 2010 | NZ | 2 | 13 | 14232 | ?? |
| *Parasmittina* | *jeffreysi* | 1202 | 1901* | Arctic | 2 | 15 | 14260 | ?? |
| *Parasmittina* | *solenosmiloides* | 1267 | 2015 | NZ | 2 | 15 | 17725 | ?? |
| *Patsyella* | *acanthodes* | 131 | 2011 | NZ | 2 | 14 | NO | ?? |
| *Porella* | *compressa* | 1188 | <1970* | NO | 1 | 5 | NO | ?? |
| *Porella* | *concinna* | 579 | 2018 | SWE | 2 | 13 | 14277 | ?? |
| *Porella* | *concinna* | 1169 | 2018 | SWE | 2 | 14 | 14278 | ?? |
| *Pterocella* | *scutella* | 104 | 2015 | NZ | 2 | 13 | 14025 | ?? |
| *Rhabdozoum* | *wilsoni* | 695 | 2018 | NZ | 0 | 15 | NO | ?? |
| *Rhynchozoon* | *angulatum* | 694 | 2018 | NZ | 2 | 13 | 14372 | ?? |
| *Rhynchozoon* | *zealandicum* | 127 | 2015 | NZ | 2 | 12 | 14092 | ?? |
| *Securiflustra* | *securifrons* | 551 | 2018 | SWE | 2 | 8 | NO | ?? |
| *Securiflustra* | *securifrons* | 1179 | <1970* | NO | 2 | 15 | 17457 | ?? |
| *Steginoporella* | *perplexa* | 100 | 2011 | NZ | 2 | 12 | 13700 | ?? |
| *Stephanollona* | *scintillans* | 679 | 2018 | NZ | 1 | 12 | NO | ?? |
| *Stephanollona* | sp. | 170 | 2009 | NZ | 2 | 8 | NO | ?? |
| *Terminocella* | n.sp. | 194 | 2010 | NZ | 2 | 15 | NO | ?? |
| *Tessaradoma* | *boreale* | 1183 | <1970* | NO | 2 | 13 | 14211 | ?? |
| *Turbicellepora* | *smitti* | 1182 | <1970* | NO | 0 | 14 | NO | ?? |

**Figure 1.**
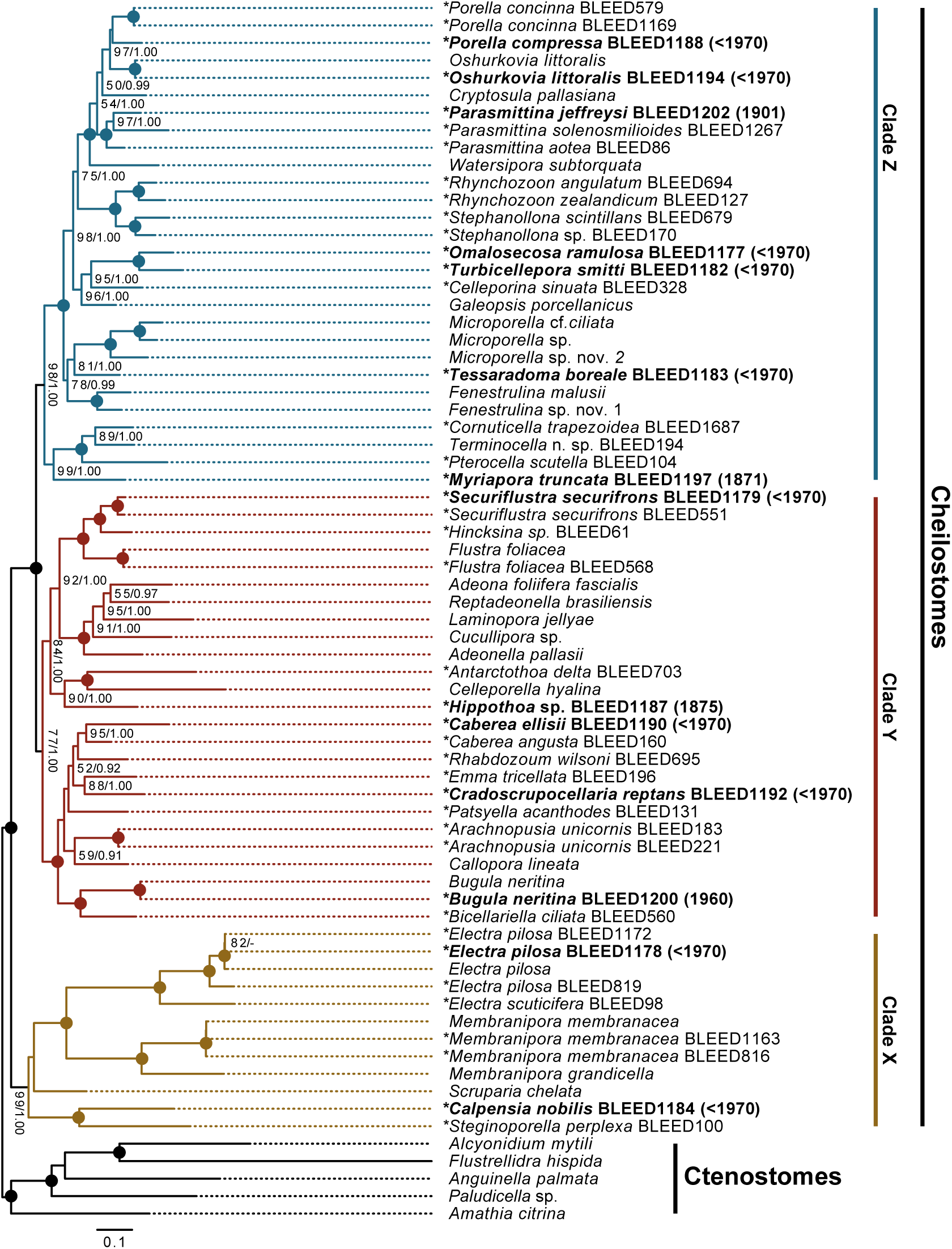
The inferred phylogeny of cheilostomes based on 17 genes from historical and recently sampled material. Maximum likelihood topology of 65 cheilostome ingroup taxa and 5 ctenostome outgroup taxa with 8324 nucleotide and amino acid characters inferred using RAxML (100 heuristic searches and bootstrap of 500 pseudoreplicates). The numbers on the internal nodes are ML bootstrap values (BS from RAxML) followed by posterior probabilities (PP from MrBayes). Circles indicate 100 BP and 1.00 PP. Only BS >50 and PP >0.95 are shown, dash indicates values below this. * indicates taxa generated in this study. Bold font indicates historical samples with sampling year in brackets. Clade X (blue), Clade Y (red), and Clade X (brown) discussed in the text are highlighted. See Table S1 for gene list.

### Phylogenetic reconstruction

Maximum likelihood (ML) phylogenetic analyses were carried out for each single gene alignment using the “AUTO” parameter in RAxML v8.0.26 (Stamatakis, 2006) to establish the evolutionary model with the best fit. The general time reversible (GTR+G) was the preferred model for the four rRNA genes (18s, 28s, rrnS and rrnL), and MtZoa+G for all 13 protein coding genes. The concatenated datasets, divided into rRNA and protein gene partitions, each with its own separate gamma distribution were analyzed using RAxML. The topology with the highest likelihood score of 100 heuristic searches was chosen. Bootstrap values were calculated from 500 pseudo-replicates.

Bayesian inference (BI) was performed using a modified version of MrBayes incorporating the MtZoa evolutionary model (Huelsenbeck & Ronquist, 2001; Tanabe, 2016) for the samples represented in the final supermatrix (Fig. 1). The dataset was executed, as before, with rRNA and protein gene partitions under their separate gamma distributions. Two independent runs, each with three heated and one cold Markov Chain Monte Carlo (MCMC) chain, were initiated from a random starting tree. The MCMC chains were run for 20,000,000 generations with trees sampled every 1,000^th^ generation. The posterior probabilities and mean marginal likelihood values of the trees were calculated after the burnin phase. The average standard deviation of split frequencies between the two runs was <0.01, indicating convergence of the MCMC chains.

## Results

We successfully sequenced and assembled 43 cheilostome colonies representing 37 species from 30 genera (Table 1). Fourteen of these come from historical samples, each representing a separate genus. The oldest known historical sample we assembled was of *Myriapora truncata* (BLEED 1197) from 1871. The 14 successfully assembled historical cheilostome samples had a total DNA input range of 1.3 – 304 ng for Illumina library preparation (Table S1). Of the 21 sequenced historical samples, one failed to provide any sequence read data. Of the 20 assembled historical samples, one was removed from the dataset based on the minimum gene number used for phylogenetic inference (Table S1), a result of low sequencing depth. For an additional four of the 20 assembled samples, no mitogenome or rRNA operon genes for the intended cheilostome target were identified with blastn, despite adequate sequencing depth (Table S1). And for a remaining sample, the assembly provided a limited contig number with no cheilostome target identifiable. The negative control provided 3.6 million reads that upon assembly gave 50 contigs >500 bp, with a 1,561 bp maximum. All assembled contigs from the negative control were attributed to contamination from *Canis familiaris* and *Homo sapiens*.

Our mitochondrial comparative phylogeny of three assembly methods (GetOrganelle, NOVOplasty and SPAdes) with different seeds for the historical samples demonstrated congruence (Fig. S1). The nodes subtending each sample are fully supported monophylies. The only observed discrepancy between the three assembly methods is seen for *Porella compressa* (BLEED 1188), where the sequence assembled with NOVOplasty using a modern *P. concinna* (BLEED 579) sample as seed, groups with the latter species, rather than itself. The NOVOplasty assembly from *Porella compressa* (BLEED 1188) recovered only two genes (*nad3* and *nad6*) representing 124 (3%) of 4144 inferred characters. The SPAdes assembly, in contrast, recovered five of 15 mitochondrial genes representing 1295 or 33% of the inferred characters. We also observe variation in the genes that are recovered between assembly methods for other specimens. The *de novo* method using SPAdes recovered the most mitochondrial genes per sample on average (Fig. S1). This was followed by the iterative method of NOVOplasty, with its results independent of seed input. Lastly, GetOrganelle consistently recovered the fewest mitochondrial genes per sample. Additionally, GetOrganelle failed to assemble in approximately 30% of cases. Despite these differences, the phylogeny demonstrates congruence among the assembly methods, with all samples, but the sequence from *P. compressa* using NOVOplasty, being monophyletic across assemblies.

Given the phylogenetic congruence of the assembly methods, we concatenated genes from these separate assemblies to form a final matrix with each sample represented only once (see methods and Table 1) to infer the phylogeny presented in Fig. 1. Specifically, the input for each specimen to this supermatrix utilized the assembly method that gave the highest number of annotated genes. When there is an equal number of recovered genes among assemblies for a given sample, prioritization was as follows: the *de novo* assembly of SPAdes, before that of NOVOplasty utilizing a SPAdes seed, and lastly NOVOplasty using a seed from a close relative (Fig. S1).

In presenting our main phylogeny we only highlight relationships represented by families constituting two or more genera receiving at least moderate support (>70 bootstrap support). The ingroup (cheilostomes) is monophyletic and separated from the ctenostome outgroup with full support 100 bootstrap (BS) / 1.00 Posterior Probability (PP) (Fig. 1). The first clade to diverge within the cheilostomes (Clade X, Fig. 1) is a highly supported (99 BS /1.00 PP) grouping including *Calpensia*, *Steginoporella*, *Scruparia*, *Membranipora* and *Electra*, with the latter two genera, forming a fully supported monophyly. The remaining cheilostome taxa form a fully supported group divided into two main sub-clades. The first sub-clade (Clade Y, Fig. 1) is moderately supported (77 BS / 1.00 PP), and further split into two groups. The basal of these two groupings is a fully supported lineage of two bugulids. The candids comprising *Cradoscrupocellaria* and *Emma* have moderate support (88 BS / 1.00 PP), but place paraphyletic to a highly supported (95 BS / 1.00 PP) monophyly of two *Caberea* species, from the same family. The terminal of these two groupings receives moderate support (84 BS / 1.00 PP) with a better resolved internal branching pattern than the previous clade. In ascending order, we recover a lineage of three hippothoids with high support (90 BS / 1.00 PP), the adeonids represented by five genera with full support, and a flustrid clade, again fully supported and harboring three genera. The latter two family groupings additionally form a highly supported (92 BS / 1.00 PP) monophyly. The second main sub-clade (Clade Z, Fig. 1) is highly supported (98 BS / 1.00 PP), and further splits into multiple lineages. Again, in ascending order, a fully supported lineage of three catenicellid genera that form a highly supported (99 BS / 1.00 PP) sister relationship with *Myriapora truncata*, and excluded from terminal relationships with full support. We find a moderately supported (78 BS / 0.99 PP) grouping of *Fenestrulina*, *Microporella* and *Tessaradoma* excluded from the terminal split with (98 BS / 1.00 PP). A highly supported (96 BS/ 1.00 PP) monophyly of four celleporids moderately (75 BS / 1.00 PP) excluded from the subsequent split. A fully supported philodoporid grouping comprises two genera and four species. And finally, a monophyly comprising a fully supported grouping of three *Parasmittina* species is observed and highly supported (97 BS / 1.00 PP) clade formed by the species *Oshurkovia littoralis* with members of *Porella*. The placements of *Watersipora subtorquata* and *Cryptosula pallasiana* are only weakly supported within this final clade.

Of the 14 historical samples, *Bugula neritina* (BLEED 1200), *Electra pilosa* (BLEED 1178), *Oshurkovia littoralis* (BLEED 1194) and *Securiflustra securifrons* (BLEED 1179) all form fully supported monophyletic intraspecies relationships. *Caberea ellisii* (BLEED 1190) and *Parasmittina jeffreysi* (BLEED 1202) form highly (95 BS/ 1.00 PP) and fully supported monophylies, respectively, within their respective genera. Further, *Porella compressa* (BLEED 1188) places within a genus monophyly, albeit without support. *Hippothoa* sp. (BLEED1187), *Omalosecosa ramulosa* (BLEED1177) and *Turbicellepora smitti* (BLEED1182), and *Cradoscrupocellaria reptans* (BLEED1192), are all highly supported within their corresponding families. The three remaining taxa, *Calpensia nobilis* (BLEED1184), *Myriapora truncata* (BLEED1197) and *Tessaradoma boreale* (BLEED1183) all lack the inclusion of a close taxonomic relative in our phylogenetic inference. However, for *M. truncata* (BLEED1197) we present a phylogeny of *cox1* (Fig. S2) demonstrating that our historical sample forms a fully supported monophyly with that of a *M. truncata cox1* sequence obtained from NCBI (ATX63952).

In total, mitochondrial genomes of 27 of 43 samples were circularized with a size range of 13700-19704 bp (Table 1). Of these, seven were from the 14 historical samples, with the oldest being that of *Parasmittina jeffreysi* (BLEED 1202, 14260 bp, collected in 1901; Fig. 2).

## Discussion

Continued explorative expeditions and the discovery of yet unknown organisms will provide optimal organic material for DNA sequencing, and subsequent analyses of extant species. However, natural history collections are unique sources that can provide historical material for organisms that may be difficult or impossible to sample from contemporary populations. Experimentation with laboratory techniques and bioinformatic tools of such historical samples can expand the information space for our general understanding of the biology, including the genetics and phylogenetics, of both well- and under-studied species. In this study, we expanded on our knowledge of a small-bodied and understudied phylum, by applying HTS and a combination of *de novo* and iterative assembly methods.

**Figure 2.**
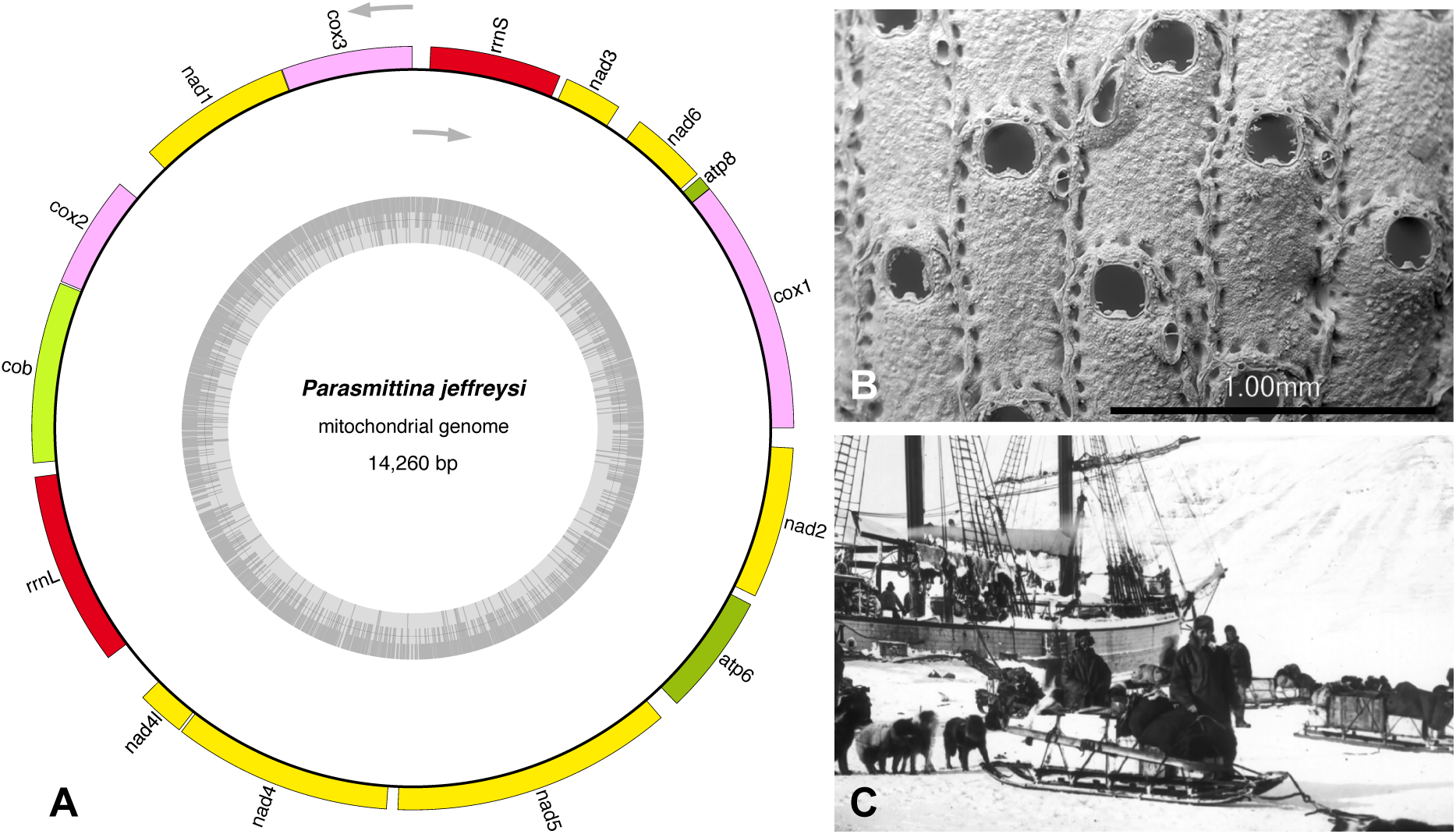
The circularized mitochondrial genome of *Parasmittina jeffreysi* (BLEED 1202). The figure shows (A) the circularised 14260 bp mitogenome of *P. jeffreysi* (BLEED1202, collected in 1901) with gene order constructed and annotated using OGDRAW v1.3.1 (Greiner, Lehwark, & Bock, 2019); the SEM (B) of the same sample, and a photograph (C) from the 1901 Norwegian FRAM II expedition where this sample was collected. The FRAM II picture is used with permission of the Norwegian Polar Institute.

### Overcoming challenges of sequencing understudied groups

While the cheilostome molecular tree is very far from complete (Fuchs et al., 2009; Knight et al., 2011; Orr, Haugen, et al., 2019; Orr, Waeschenbach, et al., 2019; Waeschenbach et al., 2012), we have contributed to this community endeavor by sequencing a number of species that have never been sequenced before and by demonstrating that it is possible, without much extra effort, to extract sequence data of many genes from old, air-dried samples. We utilized *de novo* and iterative methods to assemble our cheilostome sequence data to overcome the challenges of reference-based assemblies, due to the high degree of observed sequence variability among cheilostomes. This is especially important because systematic relationships remain largely based on morphological characters for cheilostomes (Bock & Gordon, 2013) which we know can sometimes be misleading (Orr, Haugen, et al., 2019; Orr, Waeschenbach, et al., 2019), hence seed choice is fraught with difficulties. By using *de novo* assembled sequences from the target colony itself as a reference for iterative mapping, we circumvent the difficulties of reference-choice. To explore the validity of our approach, we also explored the robustness of the phylogenetic placement by using both nominal conspecifics and congenerics as references where available (Fig. S1). Encouragingly, we find that in the majority of the cases, even historical samples can be used as their own reference for iterative mapping assemblies, at least for the samples we studied. Phylogenetic comparison of these assemblies consistently demonstrated fully supported monophylies, independent of method and the supplied reference. While SPAdes, in general, recovered the most complete assemblies and mitochondrial genes, the output of NOVOplasty was almost comparable, and even better in some cases. We propose that the iterative method employed in NOVOplasty could supplement that of the *de novo*, beckoning integration into mitogenome assembly pipelines, and removing the necessity for a reference sequence from a closely related species. Conversely, the use of GetOrganelle was superfluous for our dataset, recovering the fewest genes per sample, and having the highest failure rate. The iterative methods of GetOrganelle and NOVOplasty have previously been compared for chloroplast assemblies (de Oliveira, Duijm, & Steege, 2020; Freudenthal et al., 2019) and combined for plastid genome construction (Ma & Lu, 2019; Tan et al., 2019; Yan, Zhang, Zhao, & Yuan, 2019). Results from these previous comparative analyses, in contrast to our own, indicate GetOrganelle may be more suited to chloroplast assemblies. Additional iterative methods, such as MitoZ (Meng, Li, Yang, & Liu, 2019), have not been evaluated here and could be included in future studies. Finally, the pipeline we describe in the present study proved suitable for circularizing mitochondrial genomes (27 samples, ranging from 13700-19704 bp; Table 1), including from historical samples >100 years old (Fig. 2). While detailed mitogenome comparisons are beyond the scope of this current study, they are now available for further genome evolution research.

### An expanded cheilostome phylogeny

The phylogeny presented here is the most broadly taxonomically sampled cheilostome tree to date. In our discussion of the topology we focus on the relationships that have received at least moderate support (see results). Our tree (Fig. 1) lends statistical support to previous work on the molecular phylogeny of cheilostomes (Knight et al., 2011; Orr, Haugen, et al., 2019; Orr, Waeschenbach, et al., 2019; Waeschenbach et al., 2012), but also demonstrates well-supported branching patterns for new relationships. In summary, the most basal cheilostome clade (Clade X, Fig. 1) consists of *Electra*, *Membranipora* and *Scruparia*, reminiscent of the topology depicted by Waeschenbach et al. (2012), but we add two newly sequenced genera, namely, *Calpensia* and *Steginoporella* to this grouping. *Calpensia* (currently placed in Microporidae) is sister to *Steginoporella*, based on 14 and 16 genes respectively (Fig 1). They share a frontal, depressed, pseudoporous cryptocyst, which in *Calpensia* and *Thalamoporella*, the sister of *Steginoporella* (Knight et al., 2011), bears two opesiules while in *Steginoporella* are paired opercular indentations, outlined by the median process, forming opesiule-like lateral openings in some species. This result calls for a revision of the systematic placement of *Calpensia*, likely in the superfamily Thalamoporelloidea, where additional molecular sequence data from *Thalamoporella* could prove useful. Presently, only partial *cox1* and rrnL sequence data are available each from two different *Thalamoporella* species, and as such, were not included in this study. In the first main cheilostome clade (Clade Y, Fig. 1) we add the genera *Arachnopusia, Bicellariella, Caberea, Emma, Hincksina, Hippothoa*, *Patsyella* and *Rhabdozoum* to that previously shown (Orr, Haugen, et al., 2019; Waeschenbach et al., 2012). *Hippothoa*, the morphologically highly distinct hippothoids, places with high confidence with the previously sequenced *Antarctothoa* and *Celleporella* (Knight et al., 2011). *Hincksina* places with commonly occurring members of the family Flustridae (namely *Flustra* and *Securiflustra*). Even though *Hincksina* is in need of revision with regards to its relationship with *Gregarinidra*, the placement of this sample within the Flustridae seems beyond doubt. Moving on to the second main clade (Clade Z, Fig. 1) of cheilostomes represented in our topology, we place *Cornuticella, Myriapora, Omalosecosa, Parasmittina, Porella, Pterocella, Stephanollona, Terminocella, Tessaradoma and Turbicellepora* for the first time. *Myriapora* places as sister to the catenicellids, a position that may be the result of a limited taxon sample. Catenicellidae have a fundamentally gymnocystal-shield, with the earliest, Maastrichtian, species having a costate frontal area (A. Cheetham, pers. comm. In Banta & Wass, 1979). In contrast, Myriaporidae have a pseudoporous-lepralioid-shield and, in addition, a vastly different astogeny and colony form is observed between these two families (Ferretti, Magnino, & Balduzzi, 2007; Wass, 1983). *Tessaradoma boreale* (family Tessaradomidae) places between *Microporella* and *Fenestrulina*, two genera that are superficially similar but are now found to belong to separate families (Orr, Waeschenbach, et al., 2019). *Tessaradoma*, prior to the introduction of the family Tessaradomidae (also including the genus *Smithsonius*), has been considered within Microporellidae (MacGillivary, 1895) despite these families having contrasting frontal shield forms. A key feature of *Tessaradoma*, which partly explains its attributions to Microporellidae (among other families), is a suboral opening in the frontal shield interpreted as a spiramen (Gordon, 1993). However, broader taxon sampling is needed to confirm the affinity of *Tessaradoma* to Microporellidae, prompting a future inclusion of *Taylorius*, as previously suggested (Orr, Waeschenbach, et al., 2019), *Smithsonius*, and lastly *Siphonicytara*, which may have descended from a *Beisselina*-like (Tessaradomidae) ancestor (Gordon & Taylor, 2015).

### Verifying the placement of historical samples

The recent introduction of HTS and genome-skimming to bryozoology has brought overdue support to phylogenetic relationships, but this approach is not without its limitations (Orr, Haugen, et al., 2019; Orr, Waeschenbach, et al., 2019). Cheilostome bryozoans live in close proximity with other fouling/encrusting organisms, including other bryozoan species. As such, the isolation and subsequent amplification of solely target DNA is challenging. In our study we verified that the targeted bryozoan in 11 of our historical samples was sequenced successfully via phylogenetic placement (Fig. 1). However, *Calpensia* (BLEED 1184), *Myriapora* (BLEED 1197) and *Tessaradoma* (BLEED 1184) cannot be verified the same way. The placement of these three specimens are challenging to validate as we lack close taxonomic relatives in our phylogenetic inference to support their respective positions (Fig. 1). However, other information, beyond relative phylogenetic proximity, can help to verify specific samples. Firstly, as discussed earlier, *Calpensia* shares important morphological traits with *Steginoporella* and its relatives even though it currently nominally belongs to the phylogenetically more derived, but one-size-fits all family Microporidae (Gray, 1848; Jullien, 1888). Secondly, the *Myriapora cox1* gene has previously been amplified utilizing barcoding primers (Cahill et al., 2017). We can therefore confirm the placement of *Myriapora* (BLEED 1197) in the main phylogeny (Fig. 1) based on 17 genes, by showing its *cox1* affinity to that of *Myriapora cox1* from NCBI (Fig. S2). Lastly, the position of *Tessaradoma* (BLEED 1184) as sister to *Microporella* remains a working hypothesis until increased taxon sampling can either confirm or reject this position. We emphasize that nothing can substitute for the continued accumulation of broadly taxonomically sampled sequence and morphological data for aiding phylogenetic understanding. Lastly, and considering the confirmed phylogenetic placement of all but one of our 14 historical samples, contaminant issues can be considered negligible. Our negative control provided only a limited read number, that upon assembly gave few and short contigs attributed to mammal DNA, which were filtered out during the assembly pipeline. Considering the encrusting and fouling nature of bryozoans in general, contamination especially from the same Order is a concern independent of sample age. Here, we have treated our cheilostome samples as a metagenomes, as already demonstrated in (Orr, Haugen, et al., 2019).

### Sample age and quality versus sequencing success

The aim of our study was to demonstrate the possibility of sequencing historical, air-dried cheilostome material. As it was not our aim to systematically evaluate the lower limit of total DNA that could be amplified and HTS sequenced for bryozoans, we merely confirm success down to 1.3ng total DNA (see Table S1), although total DNA as low as 50pg is theoretically possible to amplify using the SMARTer ThruPLEX Illumina library preparation. Of the 21 historical samples we sequenced, only three samples had few or no reads, despite apparently successful library preparation. A single sample had too few assembled contigs to identify the cheilostome target. For four samples, one or more contaminants were sequenced instead (analyses not shown) and subsequently identified and removed from analyses (Table S1). As mentioned, contamination is common also for recently collected samples, highlighting the necessity of robust filtering steps in the bioinformatic pipeline. It may be expected that non-encrusting but erect colonies could be more easily freed of contaminants, resulting in improved assemblies and subsequent phylogenetic placement. However, this idea is not supported in the present study given the distribution of failed and successful samples among erect and encrusting colony forms (Table S2, Fisher’s Exact Test p >>0.05). While we did not systematically study the effect of sample age, as multiple factors could not be controlled for (including species, quality and volume of starting material and treatment during and after collecting and/or first preservation), there appears to be no simple relationship between age or preservation method and the sequencing success for cheilostome bryozoans (Table S1). Given these observations, we encourage researchers to attempt the sequencing of historical cheilostome samples of any age or species if ample material is available for experimentation.

### The importance of natural history collections

Natural history collections are treasure-troves of past information that can potentially shed light on many questions that we may not even have yet thought of, with tools that are continuously being developed. We have capitalized on historical samples stored at the Museum of Natural History at the University of Oslo, accumulated during periods where future sequencing technologies or phylogenetic estimation techniques belonged to the realm of science fiction (Short, Dikow, & Moreau, 2018; Yeates, Zwick, & Mikheyev, 2016). Many more such collections exist throughout the world and our relatively simple pipeline widens the possibilities of extracting useful sequence information from many more under-studied phyla and small-bodied invertebrates, beyond cheilostome bryozoans. This work illustrates the importance of natural history collections, and the need to store and maintain these for future generations and technological advances.

## Supporting information

SEMs

## Acknowledgements

We thank Rønning A.H. for help in the NHM Oslo collections, Thorbek B.L.G. for assistance in the sensi-lab, Mills S. for handling and export of the New Zealand NIWA (National Institute of Water and Atmospheric Research) samples, and Rust S. for sorting samples from the University of Otago. Sequencing was performed at the Norwegian Sequencing Centre (University of Oslo) and analyses done on the project node nn9244k on the Abel server (University of Oslo). This project has received funding from the European Research Council (ERC) under the European Union’s Horizon 2020 research and innovation programme (grant agreement No 724324 to LHL) and MMS has received student funding from NHM Oslo. Field work in Sweden was supported by Assemble Plus grant to RJSO (grant number 730984). MO was supported by SYNTHESYS Plus grant (grant number 823827).

## Author contributions

RJSO, SB and LHL designed the study, HM, AMS, DPG, MO, RJSO, LHL, MHR collected the material, RJSO and MMS performed the lab work and bioinformatics, MHR and EDM took the SEMs, DPG, EDM, MO identified the taxa, RJSO and LHL wrote the first draft of the manuscript which all authors commented and agreed on.

---

**Figure S1.**
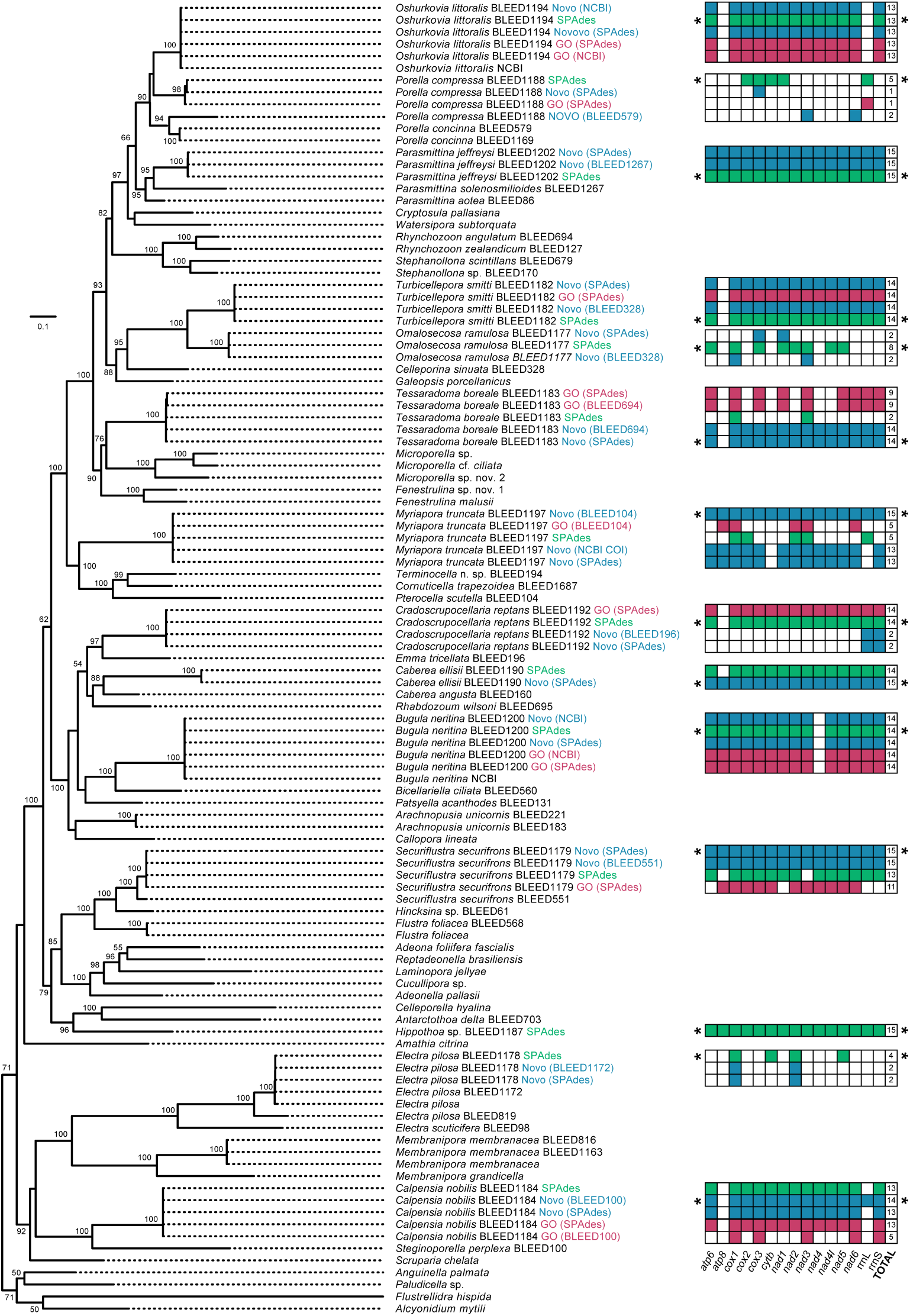
Phylogeny comparing assembly methods of historical samples based. Maximum likelihood topology of 109 taxa and 4144 nucleotide and amino acid characters inferred using RAxML (20 heuristic searches and bootstrap of 100 pseudoreplicates). Only BootStrap values BS >50 are shown. The three different assembly methods are highlighted as follows: NOVOplasty (“Novo” blue font), GetOrganelle (“GO” red font), and SPAdes (“SPAdes” green font). The reference/seed used is in brackets following the assembly method with “SPAdes” indicating the *de novo* assembly of the same sample was utilized. Note that not all methods were applicable to all samples. The table to the right of the tree shows the individual and total genes recovered (of a maximum of 15) for each sample and method, with a colored box indicating a positive result. The gene order per column is as follows: *atp6, atp8, cox1, cox2, cox3, cytb, nad1, nad2, nad3, nad4, nad4l, nad5, nad6*, rrnL and rrnS. The * to the left and right of a row indicate that the result from this assembly method was utilized in the main phylogeny shown in Fig. 1.

**Figure S2.**
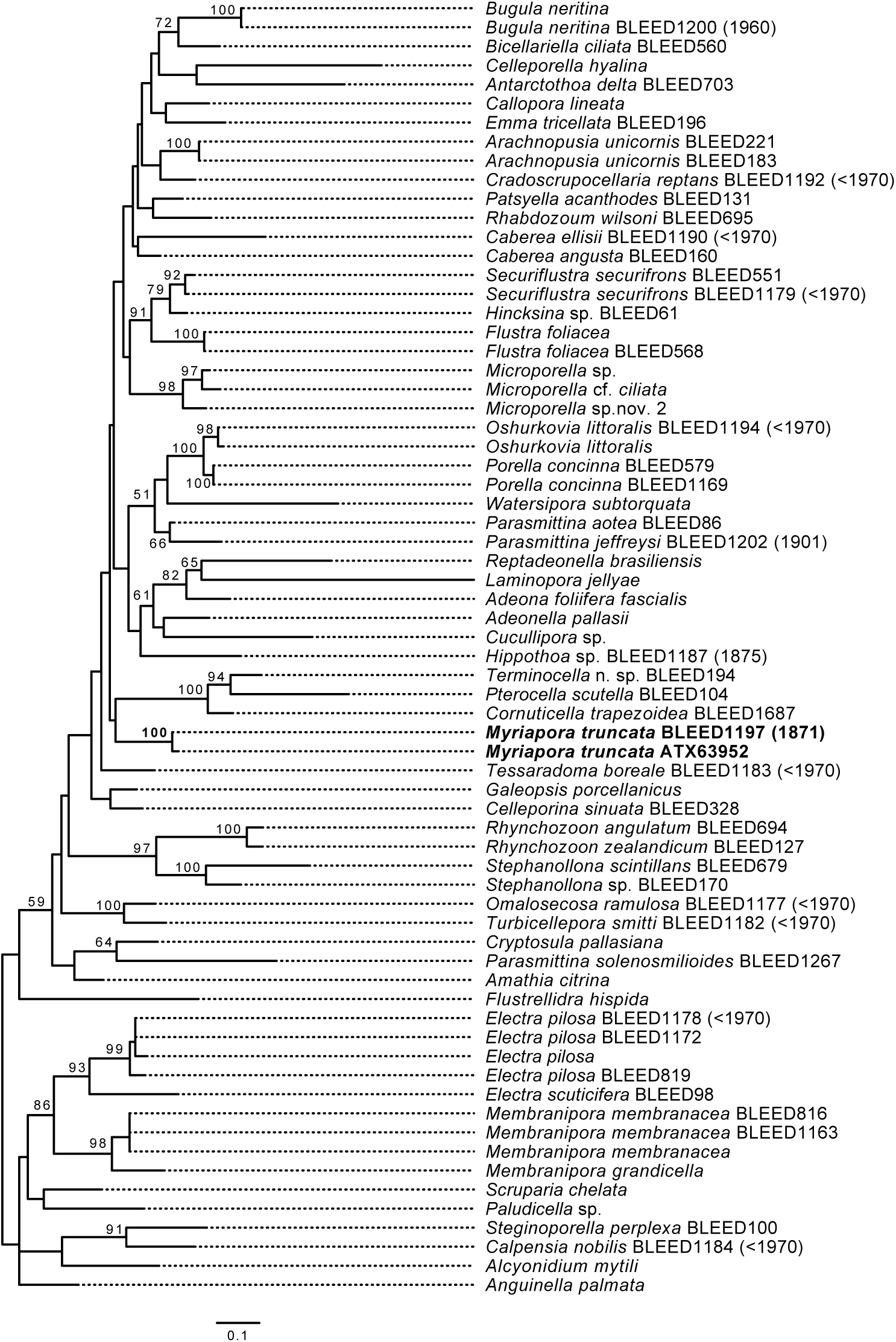
Phylogeny of *cox1* confirming the placement of *Myriapora truncata* (BLEED1197) from 1871. Maximum likelihood topology of 68 taxa and 490 amino acid characters inferred using RAxML (20 heuristic searches and bootstrap of 100 pseudoreplicates). Only Bootstrap values BS >50 are shown. The *M. truncata* (BLEED1197) from 1871 and the NCBI sequence (ATX63952) are shown in bold, as is the BS support value for this branch.

