## Supplementary material for "Molecular phylogeny of historical micro-invertebrate specimens using *de novo* sequence assembly": SEMs

### Supplementary information

Scanning Electron Micrographs (SEMs) of voucher specimens and taxonomic notes

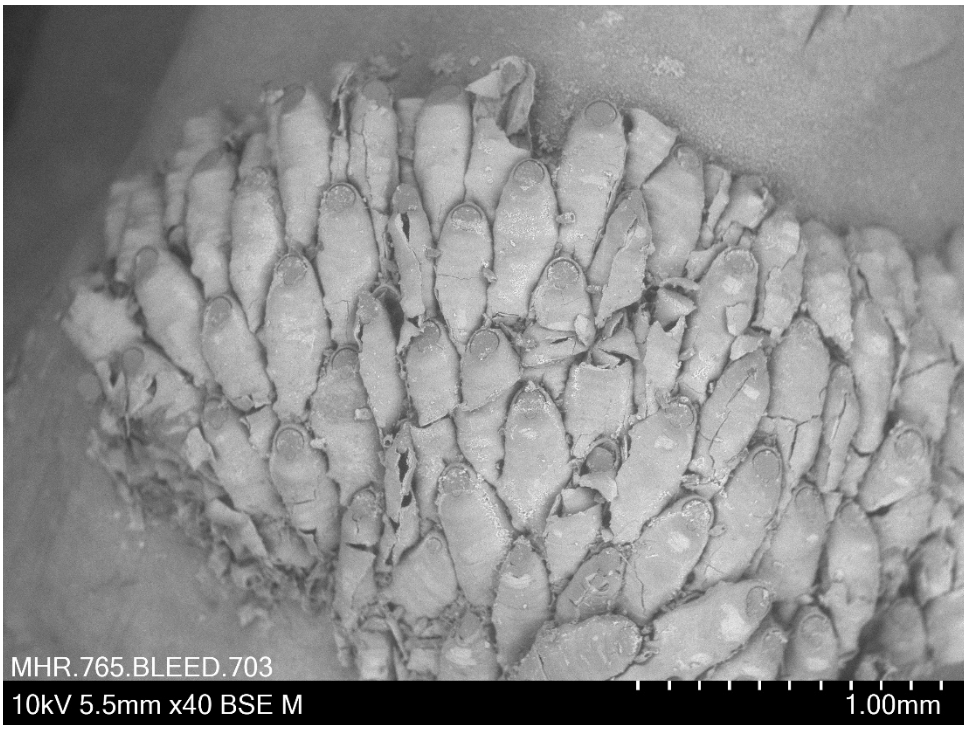


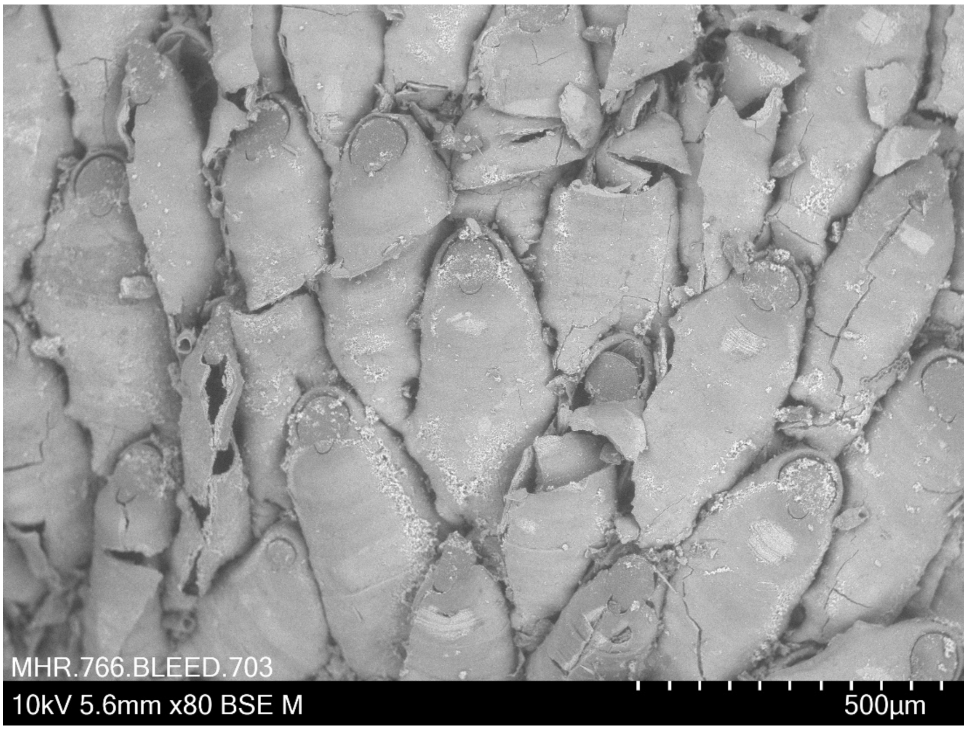


***Antarctothoa delta*** **Ryland & Gordon, 1977 BLEED 703**
Collection date 9.4.2018 (A. M. Smith, D.P.Gordon, H. Mello) Pres. ethanol
Lat: -47.221, Long: 167.608, Depth: intertidal, Shipbuilders Cove, Stewart Island, New Zealand.

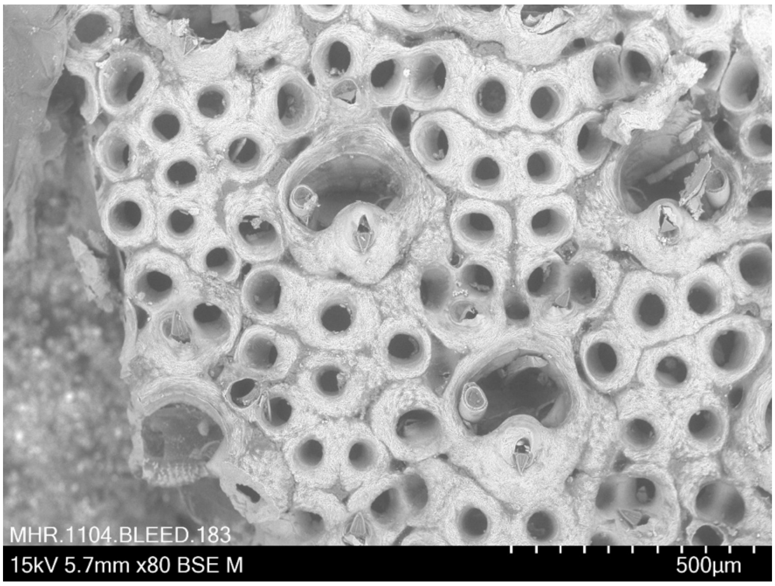


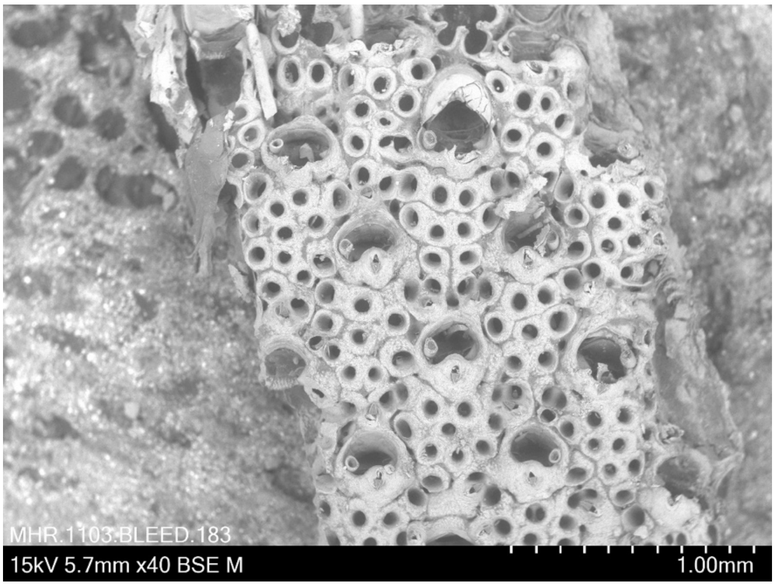


***Arachnopusia unicornis* (Hutton, 1873) BLEED 183**
Coll. 14 July 2009  Pres. ethanol
Lat: -34.373, Long: 172.923, Depth: 72 m, New Zealand
Oceans Survey 2020 Station ID: TAN0906/154 NIWA 56548

<u>Taxonomic notes</u>: The illustrated specimen has perforations in sectors, reminiscent of *A. perforata* but less so of *A. unicornis* [but see BLEED 221]. However, the distal rim of the orifice in *A. perforata* bears avicularia, which is not the case in this specimen. *Arachnopusia* species in New Zealand is highly morphologically variable morphologically but both the morphology and limited sequence variation suggests both BLEED 221 and BLEED 183 are *A. unicornis.*

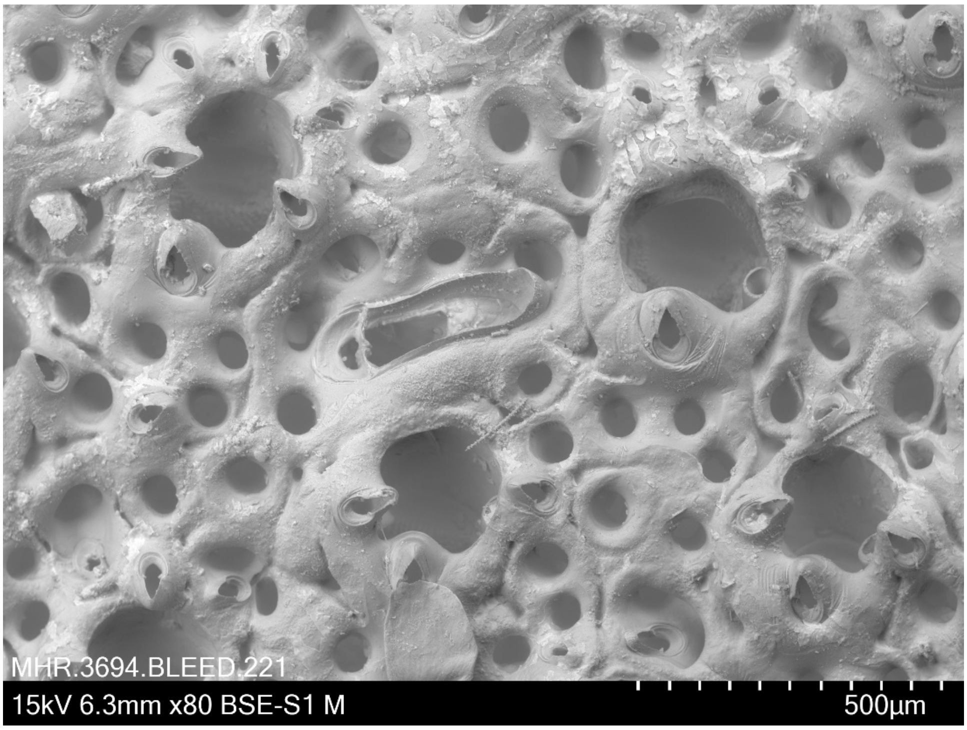


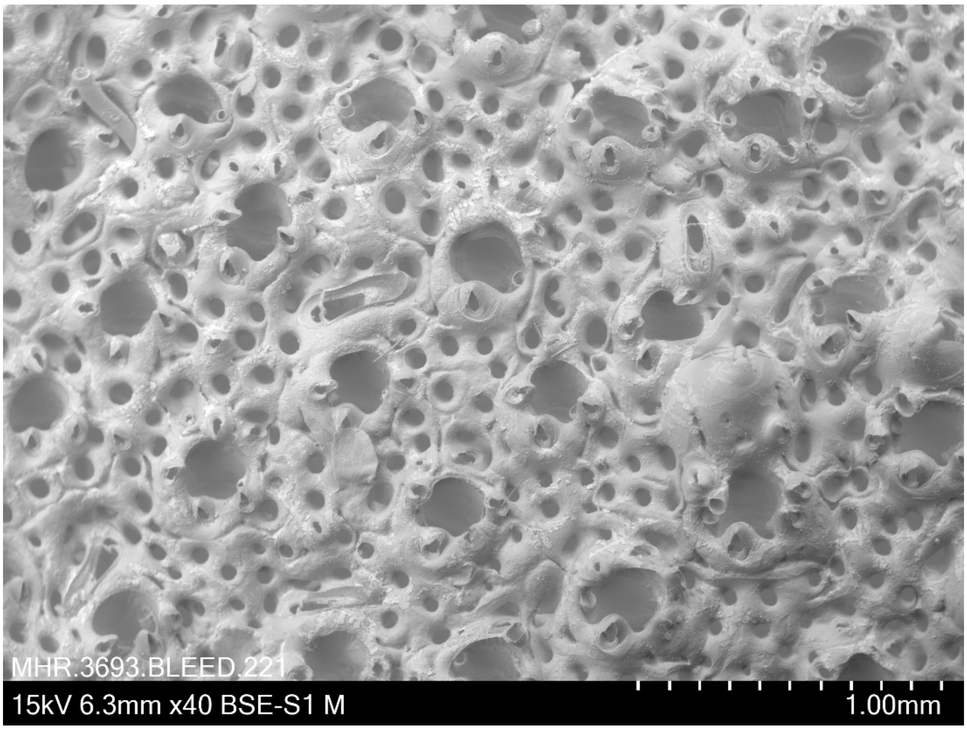


***Arachnopusia unicornis* (Hutton, 1873) BLEED 221**
Coll. 30 November 2011 (A.M.Smith) Pres. ethanol
Lat: -46.985, Long: 168.235, Depth: 47m, New Zealand

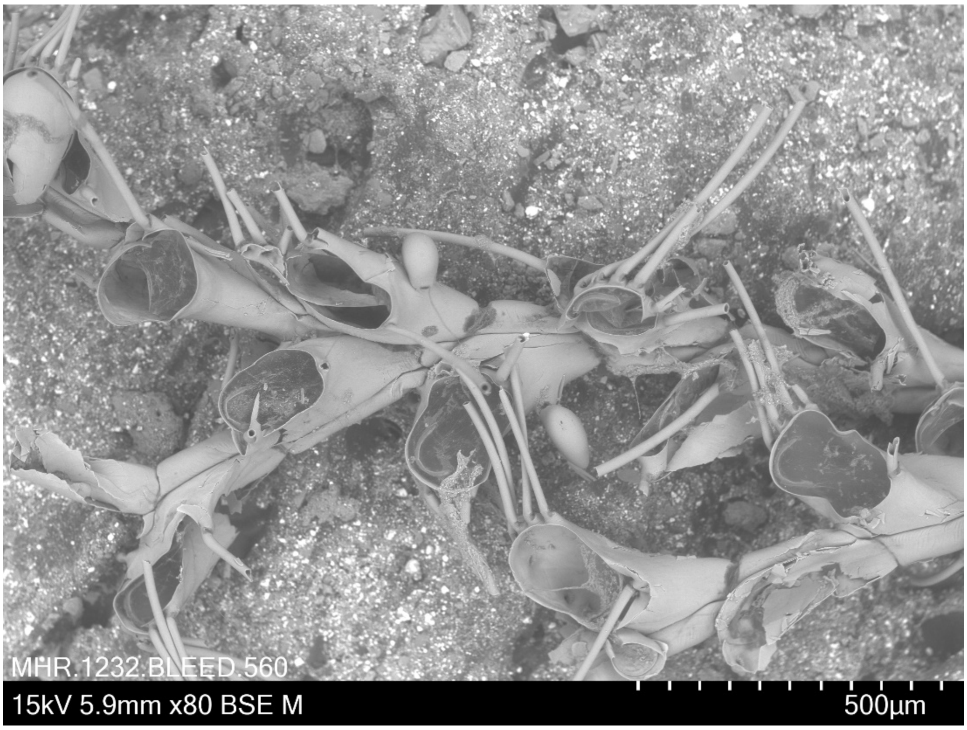


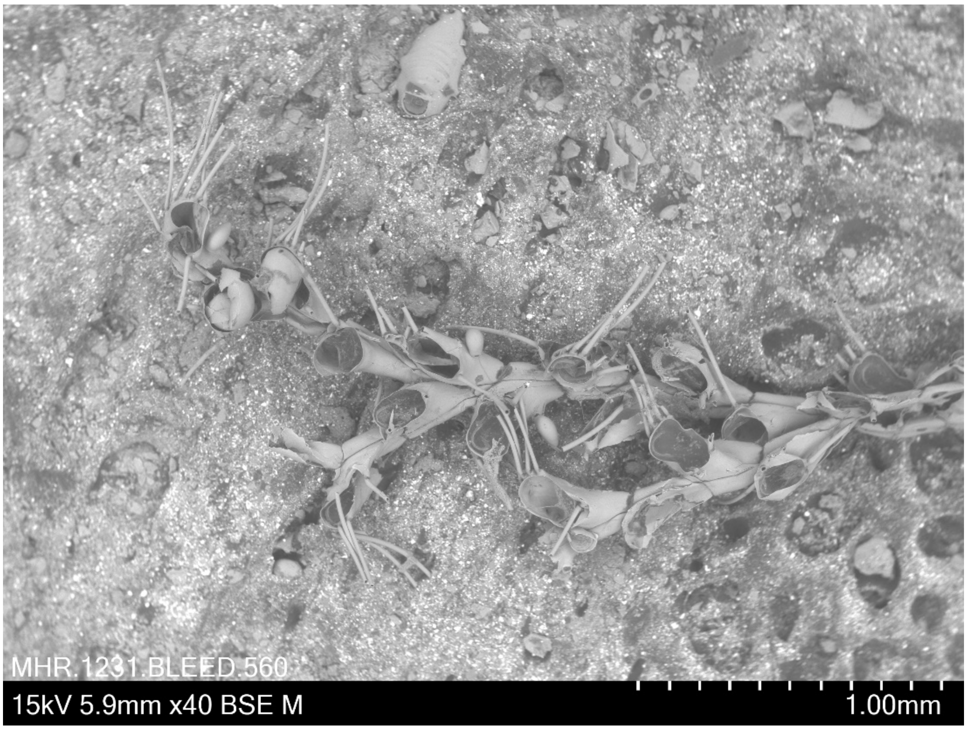


***Bicellariella ciliata* (Linnaeus, 1758) BLEED 560**
Coll. 28 May 2018 (M. Obst, R.J.S. Orr, L.H. Liow) Pres. ethanol
Lat: 58.247, Long: 11.404, 15-45 m, off Kristineberg, Sweden

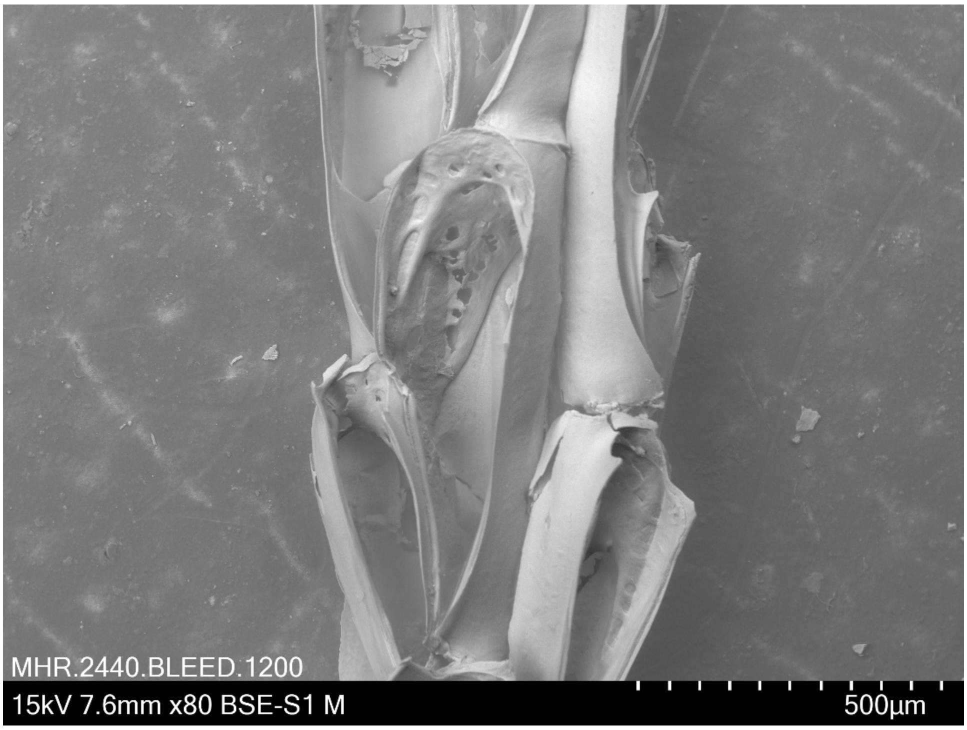


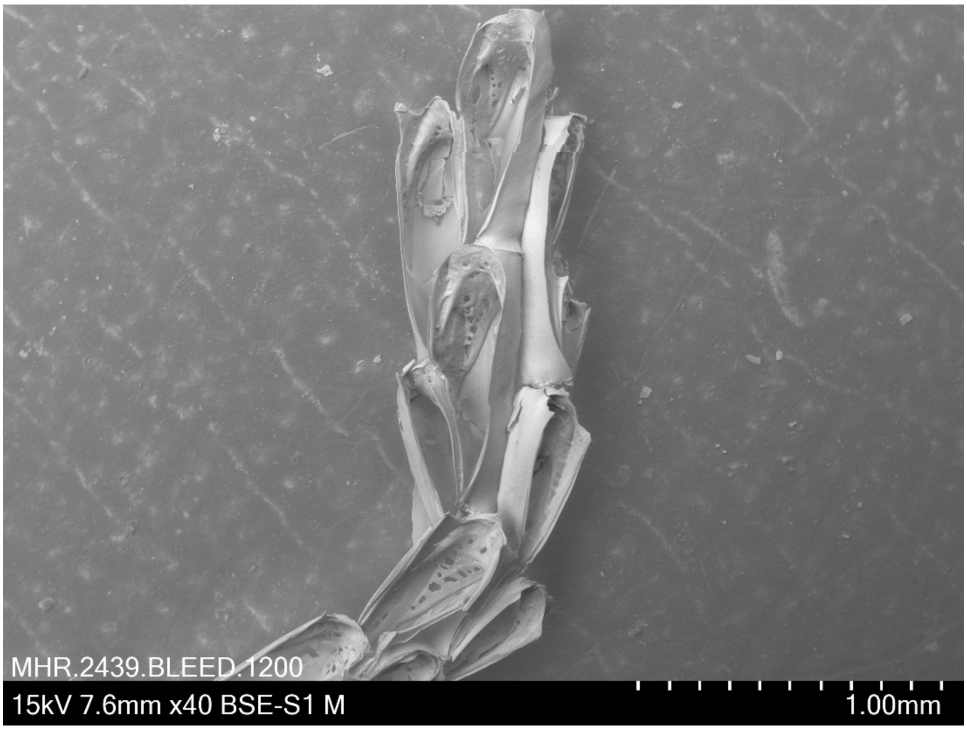


***Bugula neritina* (Linnaeus 1758) BLEED 1200**
Coll. 1871 Pres. dried
Unregistered specimen from NHM Oslo
Location: Australia. No other metadata available.

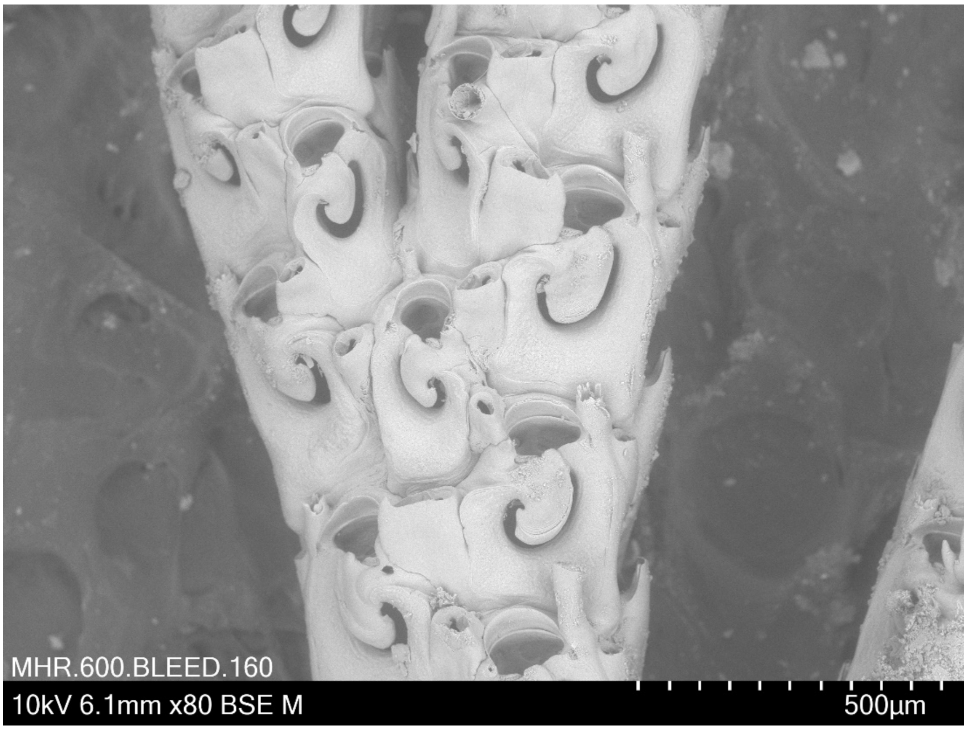


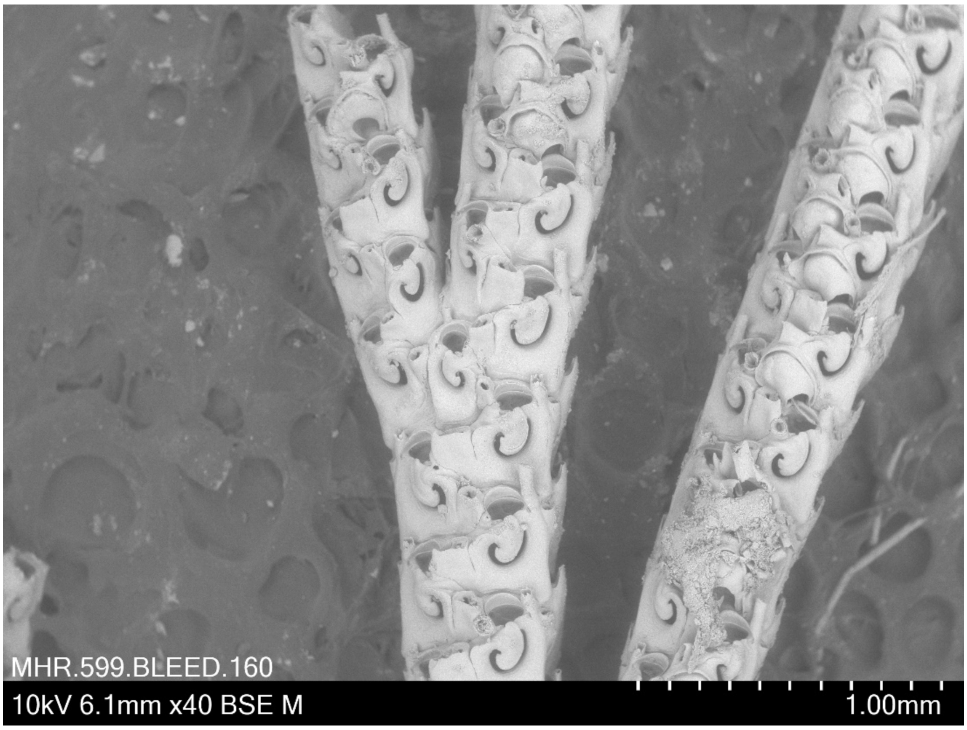


***Caberea angusta* Hastings, 1943 BLEED 160**
Coll. 5 July 2009 Pres. ethanol
Lat: -35.553, Long: 174.553, Depth: 57m, New Zealand
Station ID: TAN0906/25 NIWA 54782

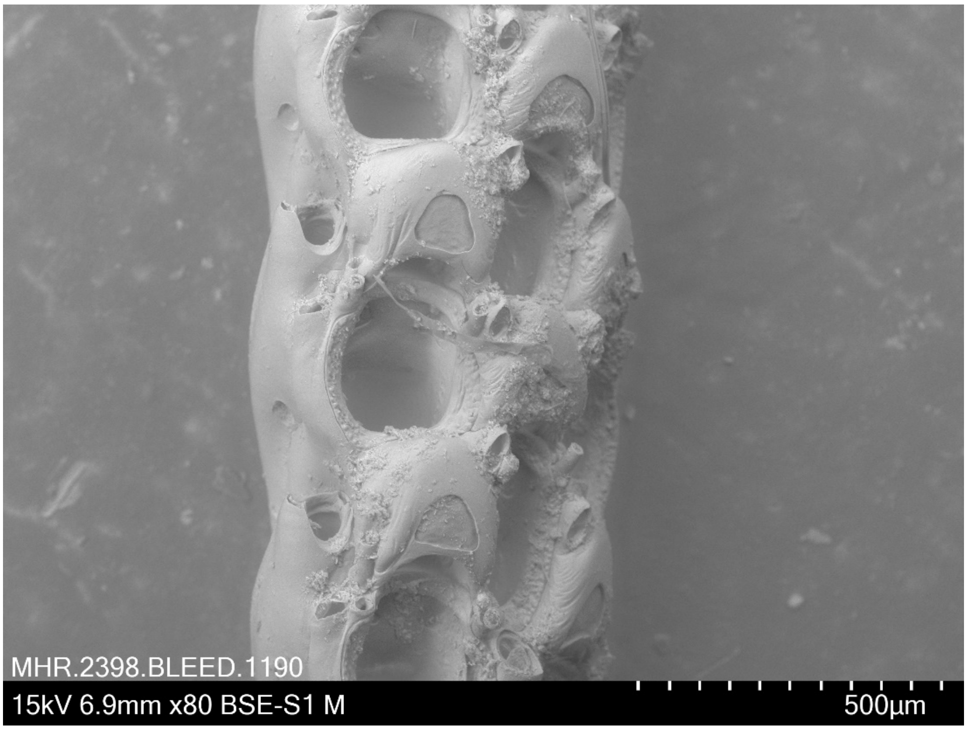


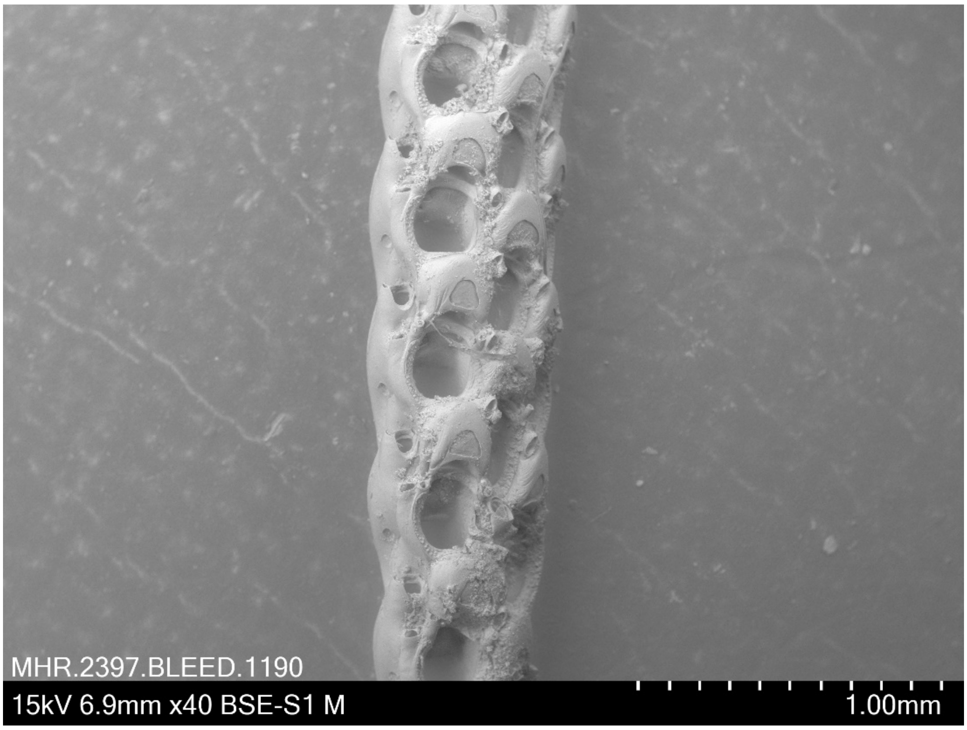


***Caberea ellisii* (Fleming, 1814) BLEED 1190**
Pres. dried
Identified by I. Vigeland in 1970 (unregistered specimen from NHM Oslo)
Location: Hammerfest, Norway. No other metadata available.

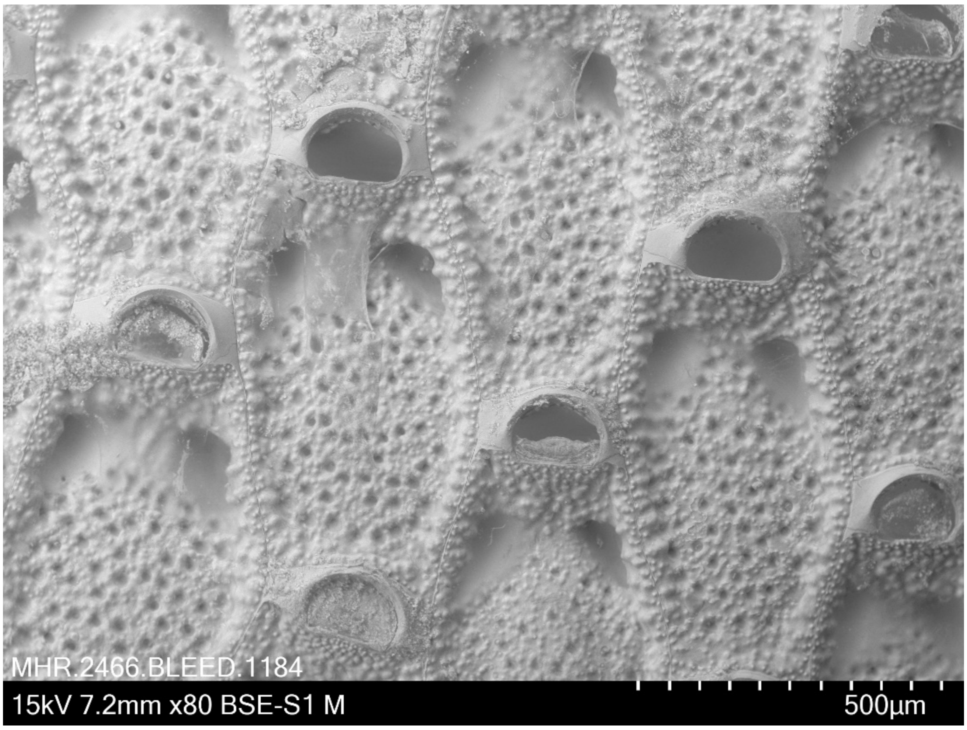


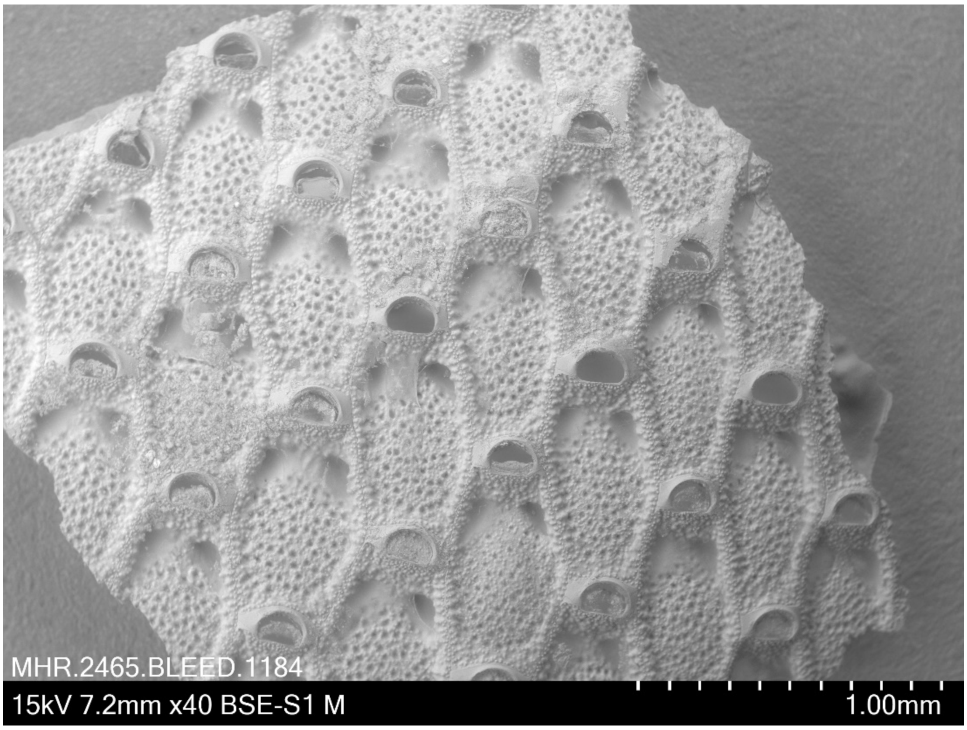


***Calpensia nobilis* (Esper, 1796) BLEED 1184**
Pres. dried
Identified by I. Vigeland in 1970 (unregistered specimen from NHM Oslo)
Location: Messina, Italy. No other metadata available.

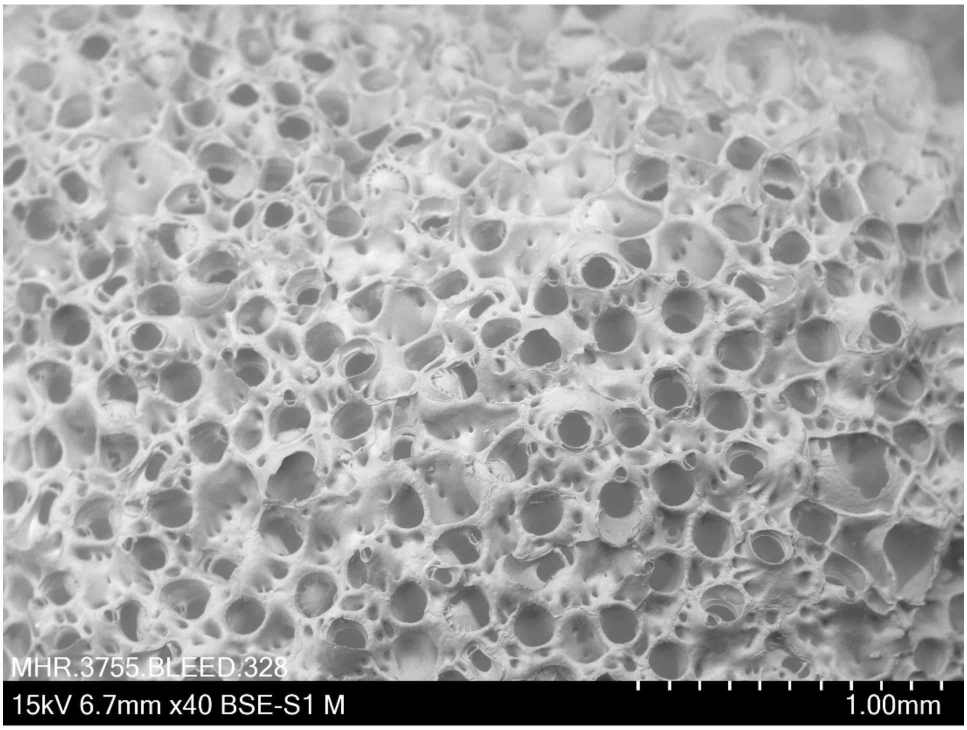


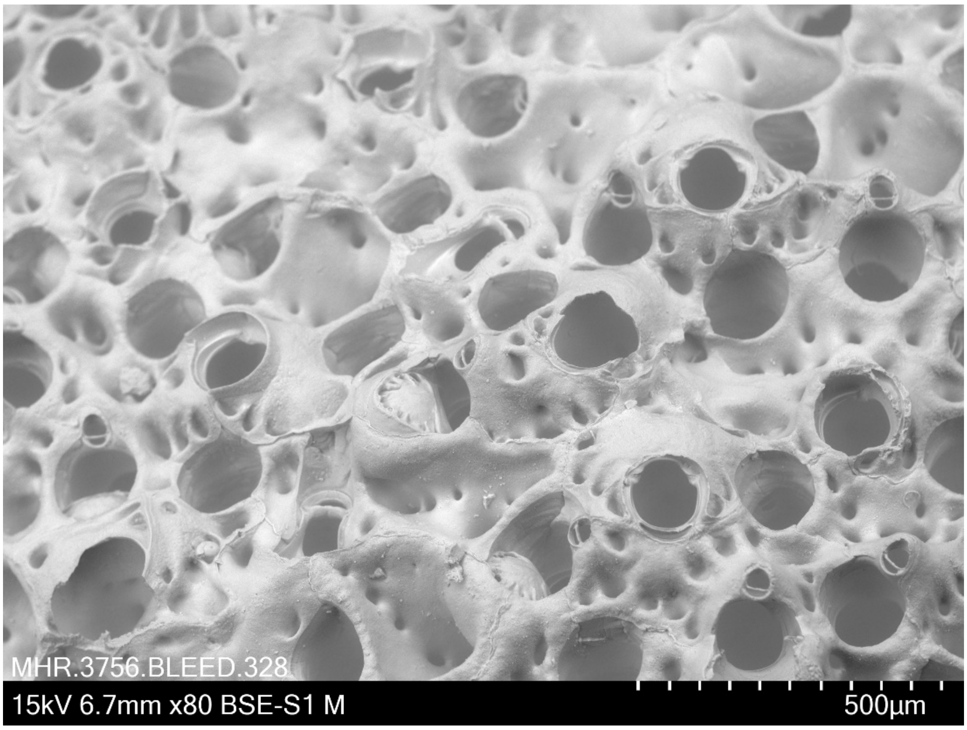


***Celleporina sinuata* Gordon, 1989 BLEED 328**
Coll. 28.5.2015 Pres. ethanol
Lat: - 35.7982, Long: 74.45948, Depth: ? m (Whangarei harbor depths), New Zealand
Station ID: WRE21050 NIWA 71326

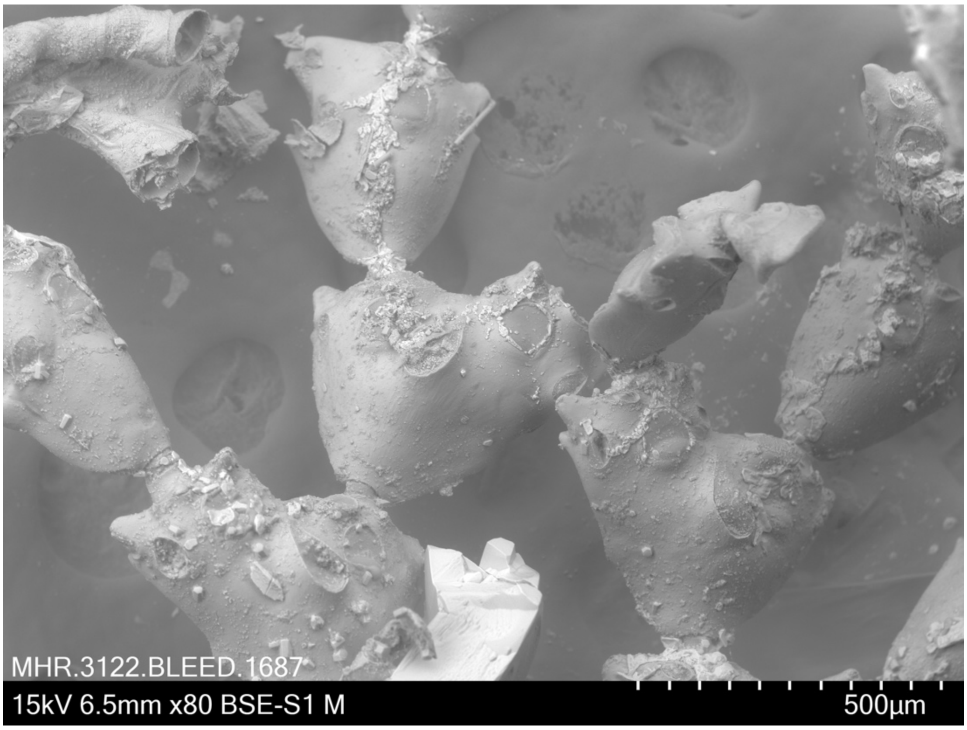


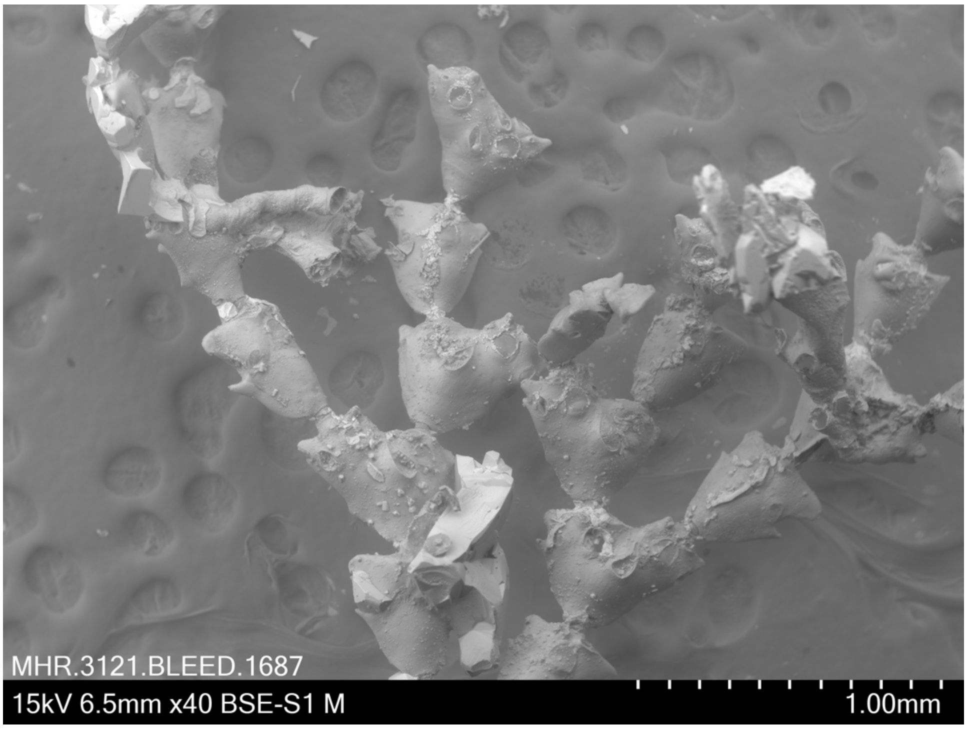


***Cornuticella trapezoidea* Powell, 1967 BLEED 1687**
Coll. 18.3.2019 (A.M. Smith, D.P. Gordon) Pres. ethanol
Lat: - 34.3800, Long: 172.86, Depth: 54.6, N of Hooper Point, E Spirits Bay, New Zealand

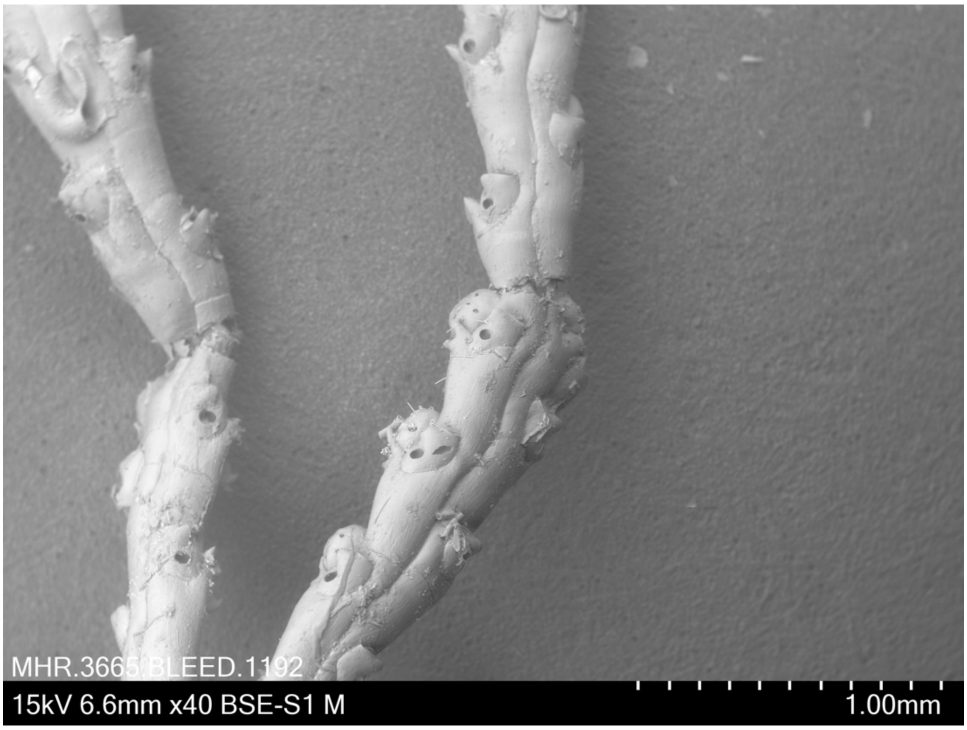


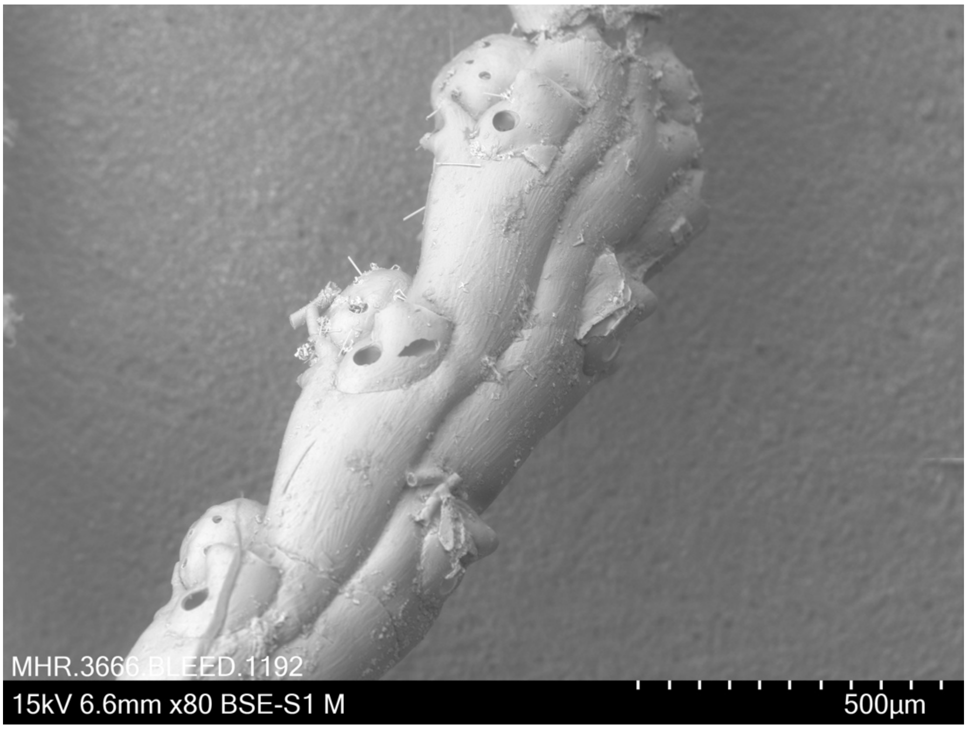


***Cradoscrupocellaria reptans*** **(Linneaus, 1758) BLEED 1192**
Pres. dried
Identified by I. Vigeland in 1970 (unregistered specimen from NHM Oslo)
Location: Bergen, Norway. No other metadata available.

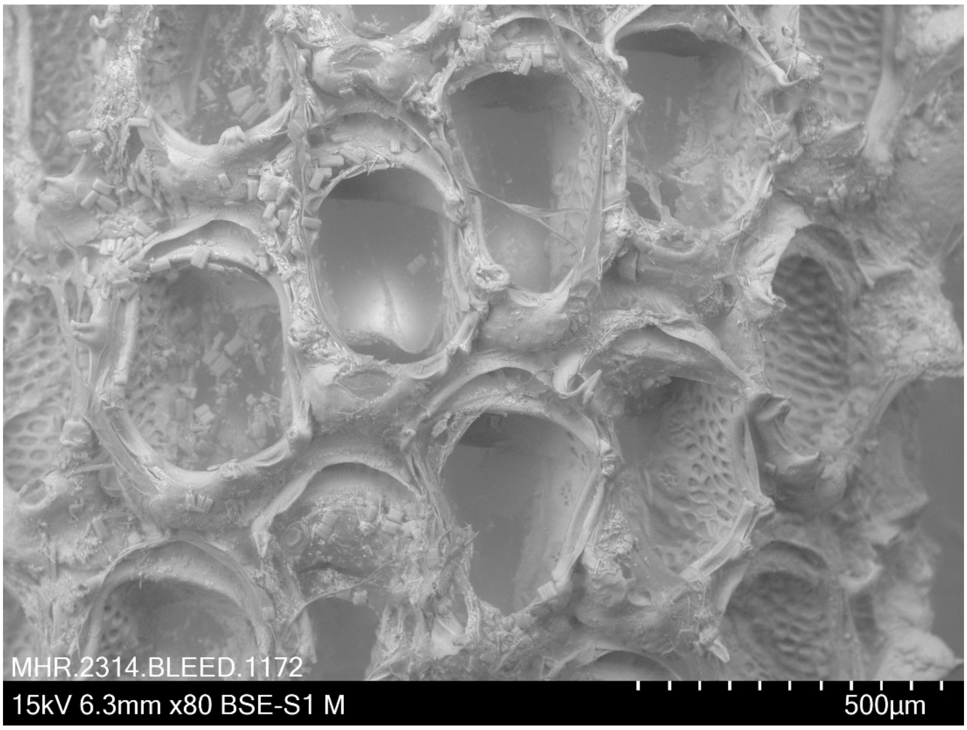


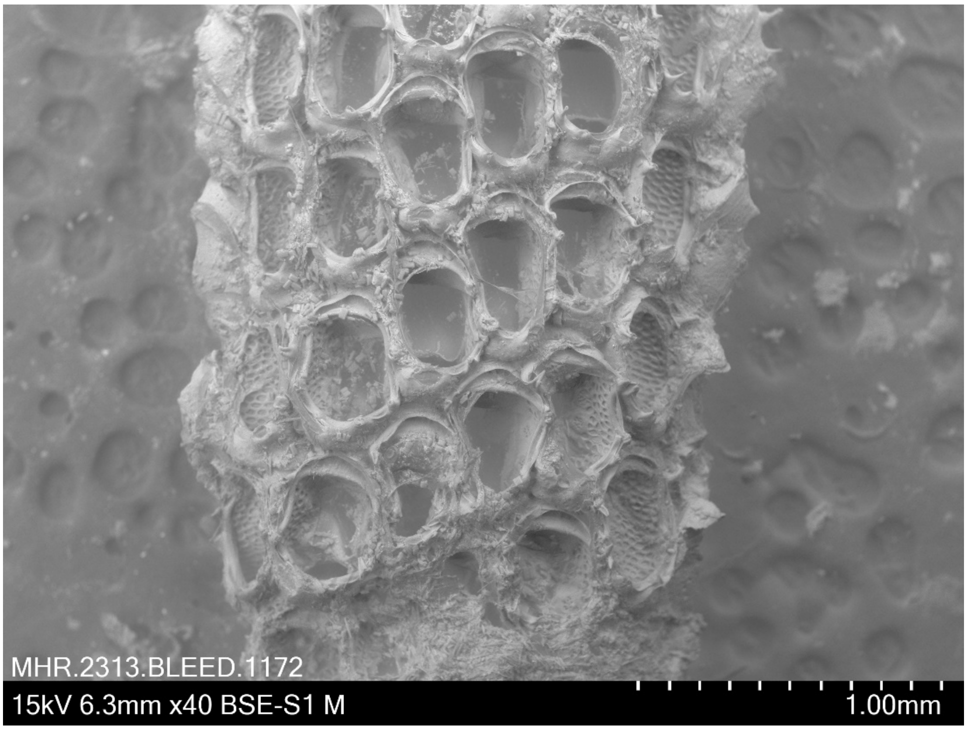


***Electra pilosa* (Linnaeus 1767) BLEED 1172**
Coll. 16.7.2018 (L.H.Liow) Pres. dried
Locality: Stråholmen off Kragerø, Norway, washed up on shore.

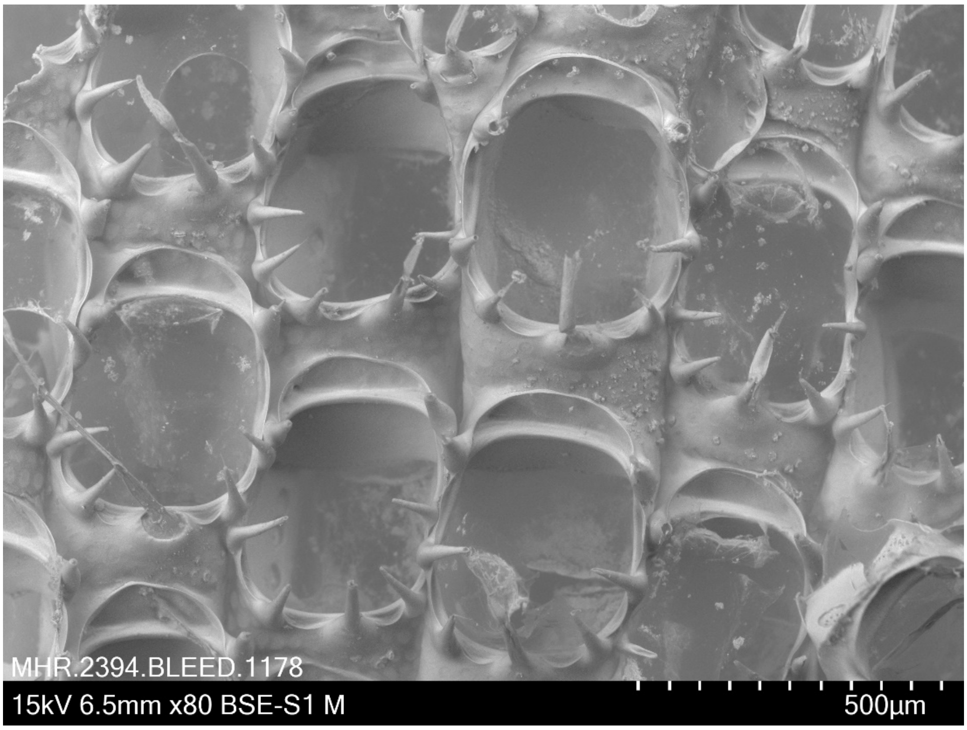


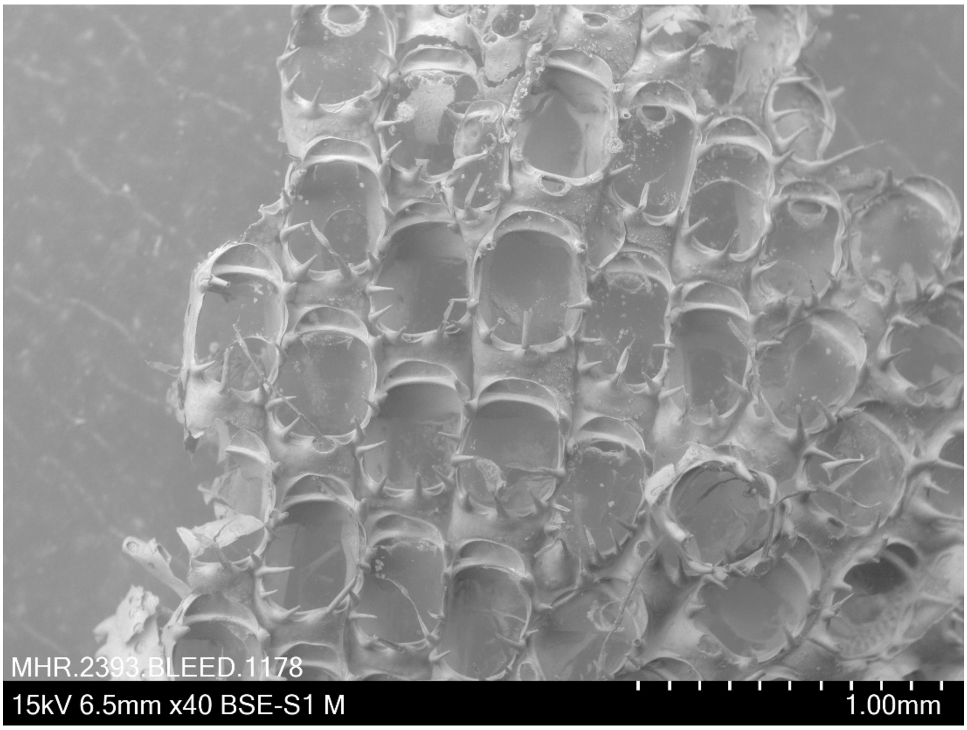


***Electra pilosa* (Linnaeus, 1767) BLEED 1178**
Pres. dried
Identified by I. Vigeland in 1970 (unregistered specimen from NHM Oslo)
no other meta data available

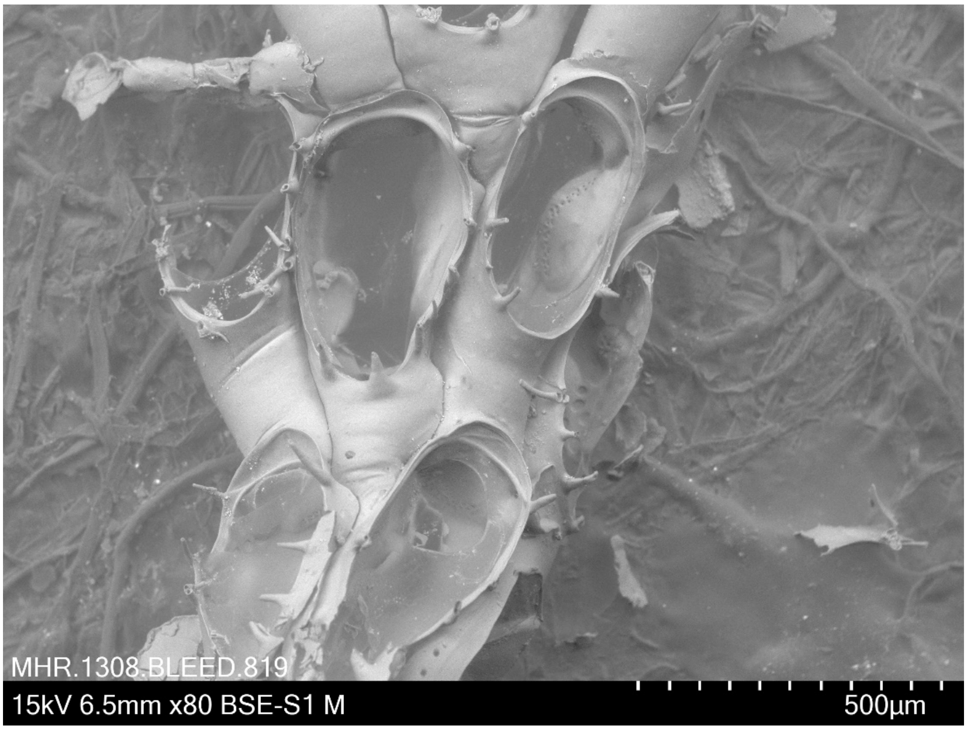


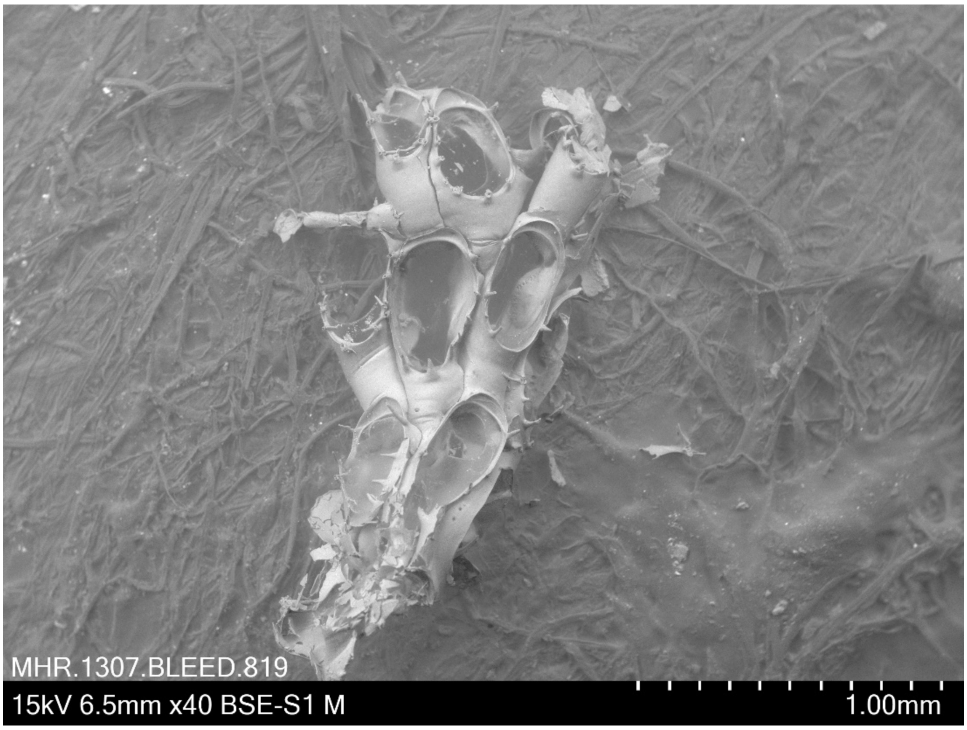


***Electra pilosa* (Linnaeus, 1767) BLEED 819**
Coll. 11 September 2018 (M.H. Ramsfjell) Pres. ethanol
Lat: 60.394, Long: 5.310, Depth: intertidal, Bergen
BER-B-S1

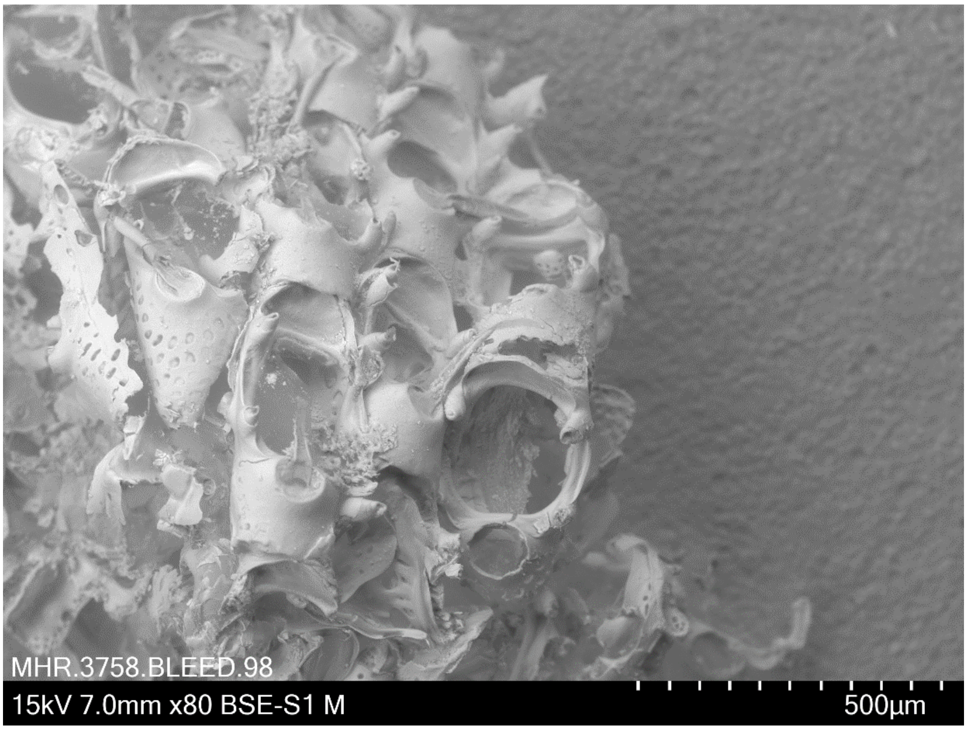


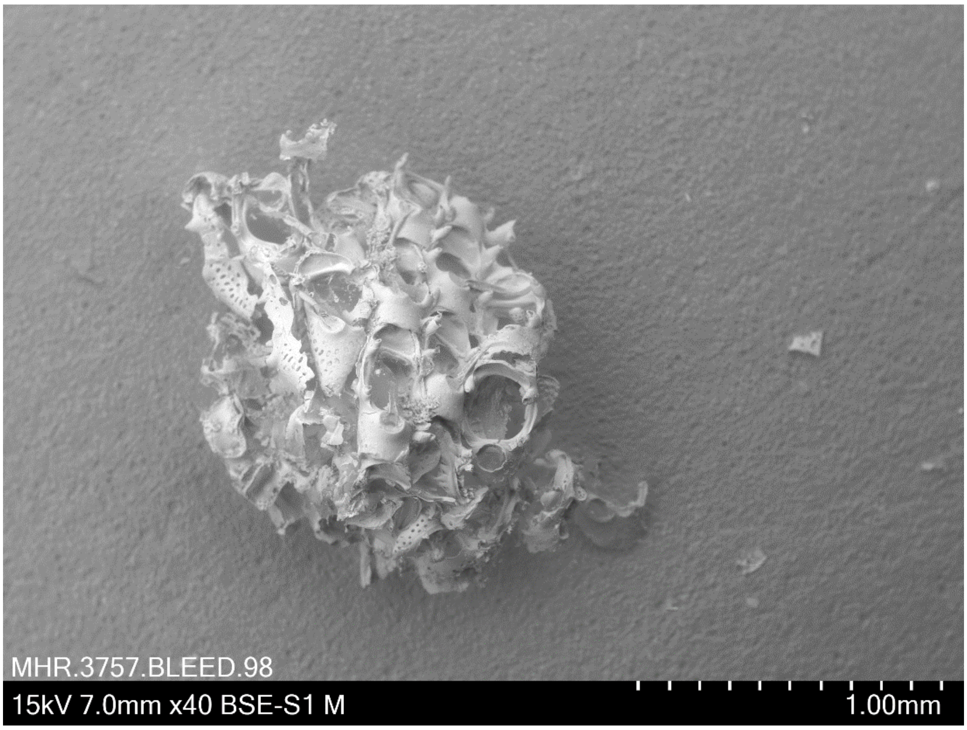


***Electra scuticifera*** **Niculina, 2008 BLEED 98**
Coll. 20.5.2015 (D.P. Gordon) Pres. ethanol
Lat -40.900, Long 176.217, Depth intertidal, Castlepoint, Wairarapa, New Zealand

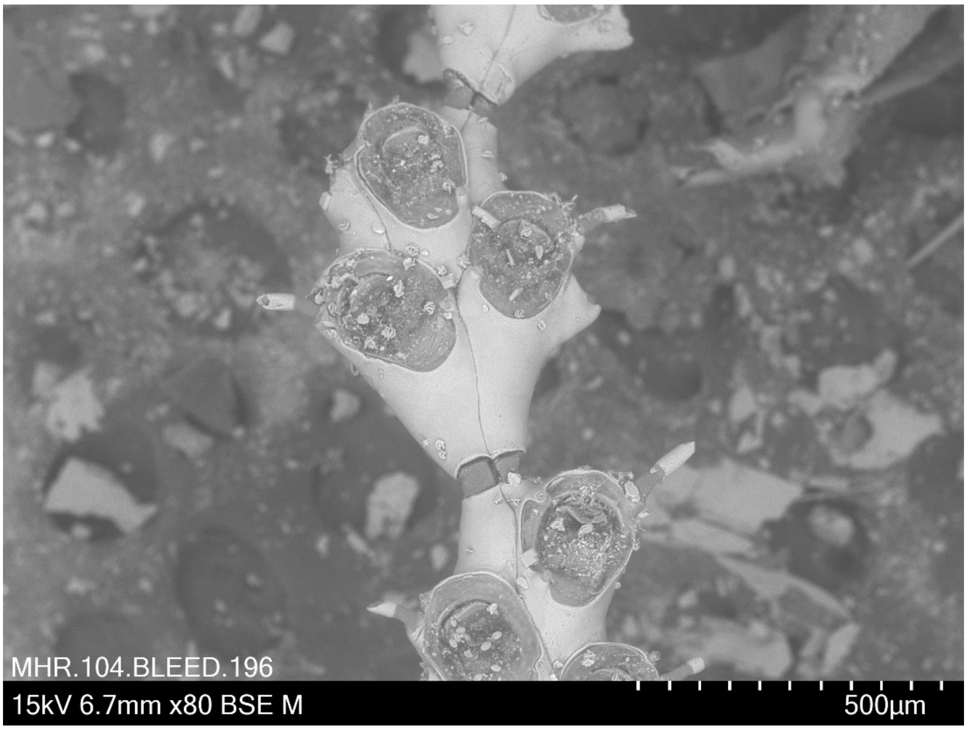


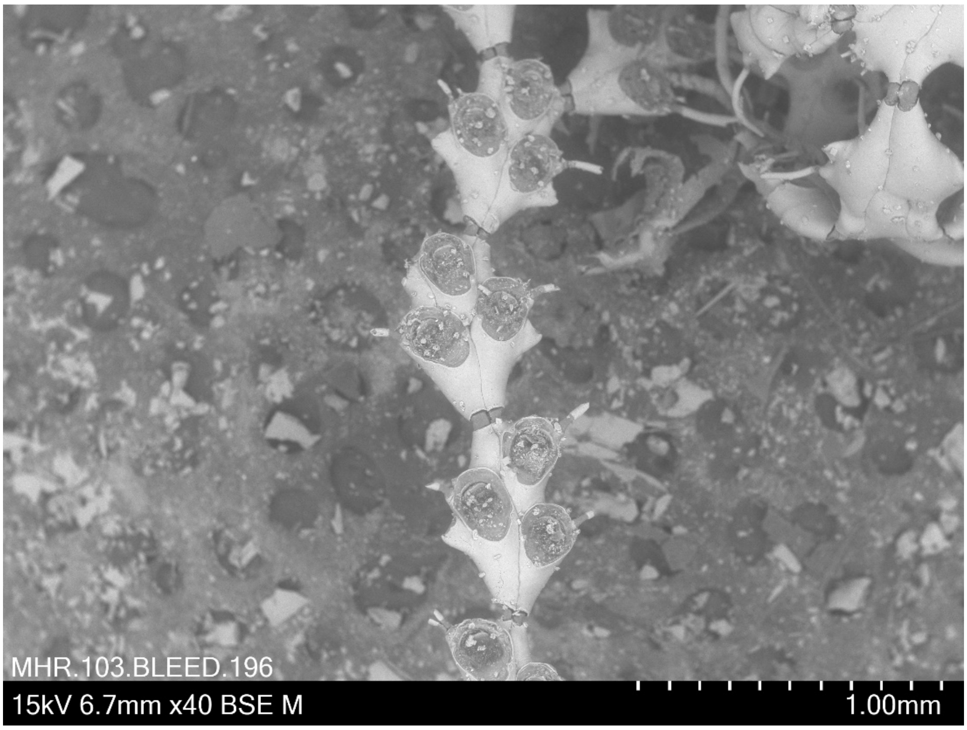


***Emma tricellata* Busk, 1852 BLEED 196**
Coll. 14 July 2009 (D.P. Gordon) Pres. ethanol
Lat: -41.08, Long: 176.09, Depth: intertidal, Orui, Wairarapa, New Zealand

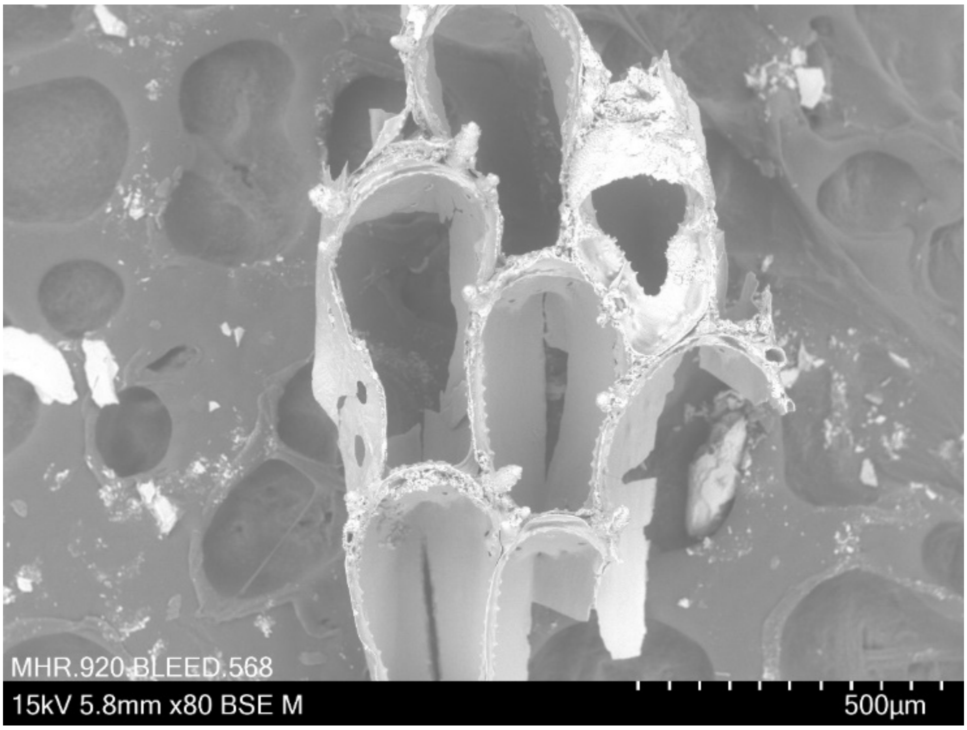


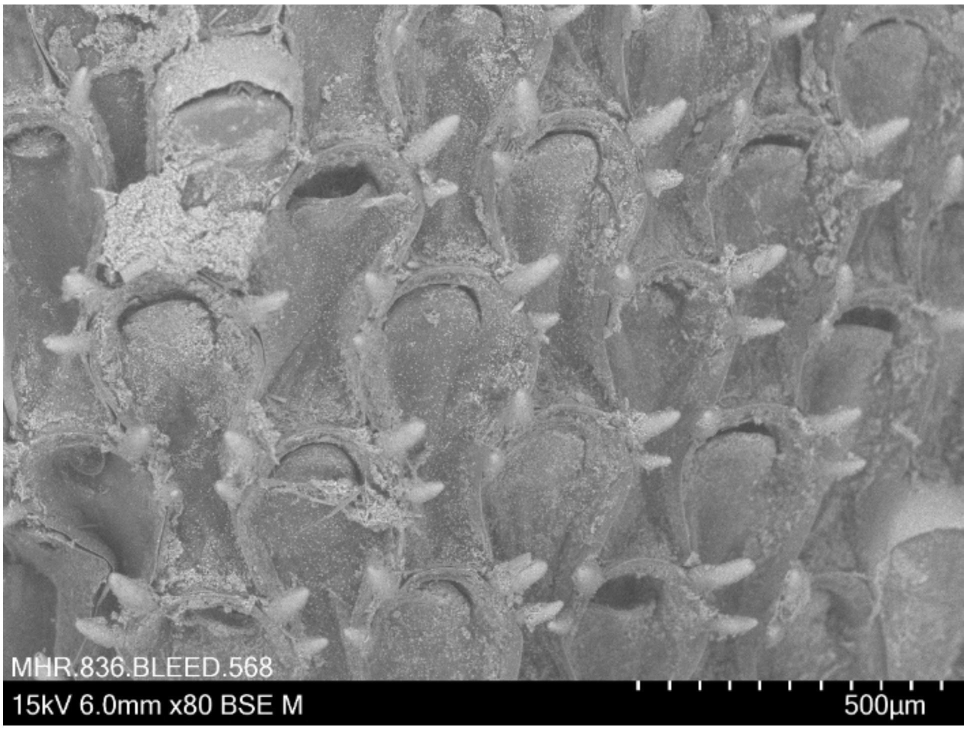


***Flustra foliacea*** **(Linnaeus, 1761) BLEED 568**
Coll. 28 May 2018 (M. Obst, R.J.S. Orr, L.H. Liow) Pres. ethanol
Lat: 58.247445, Long: 11.404071, 15-45 m, off Kristineberg, Sweden

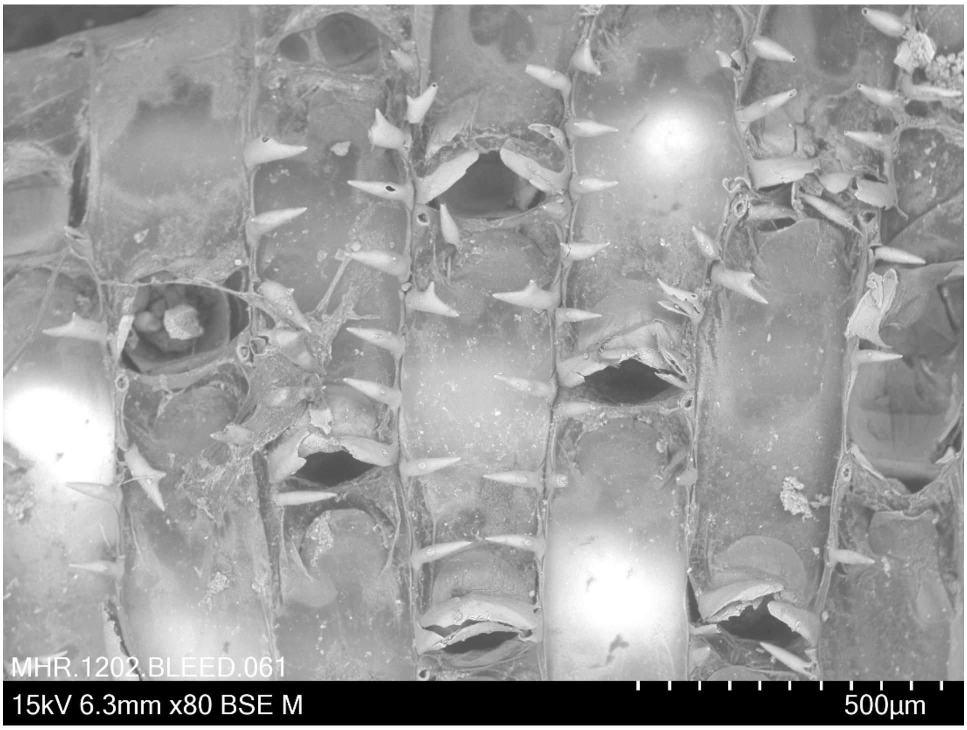


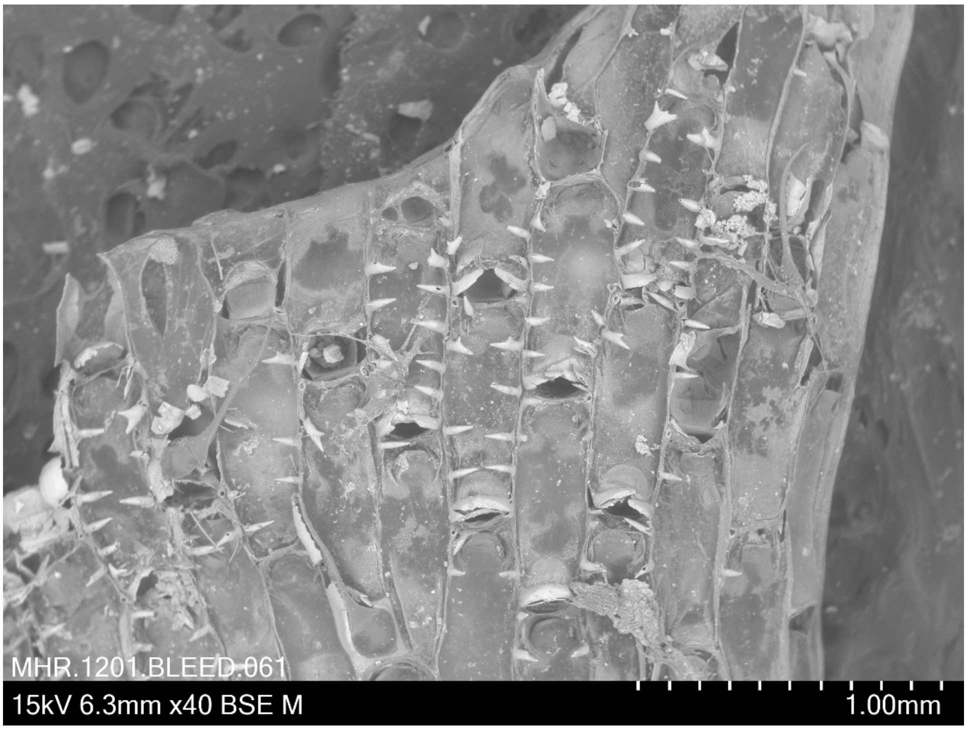


***Hincksina* sp. BLEED 61**
Collection date 4 July 2009 Pres. ethanol
Lat: 35.473, Long: 174.678, Depth: 111 m, New Zealand
Oceans Survey 2020 Station ID: TAN0906/8 NIWA 54560
Taxonomic notes: Compare *Hincksina* Norman, 1903 with *Gregarinidra* Barroso, 1949 with which it may be synonymized.

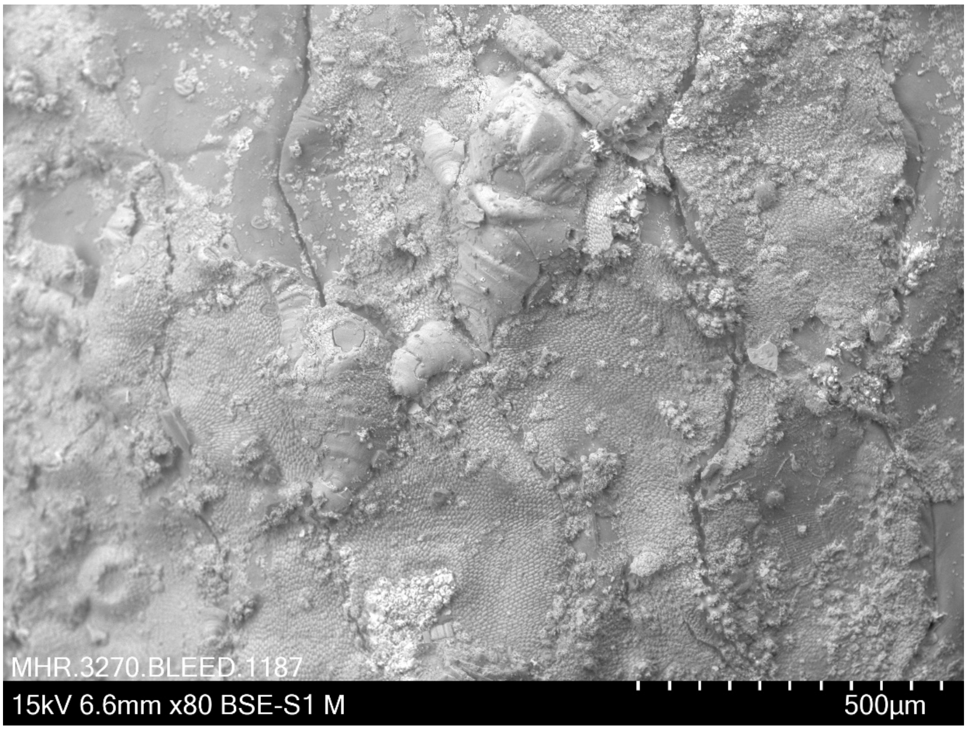


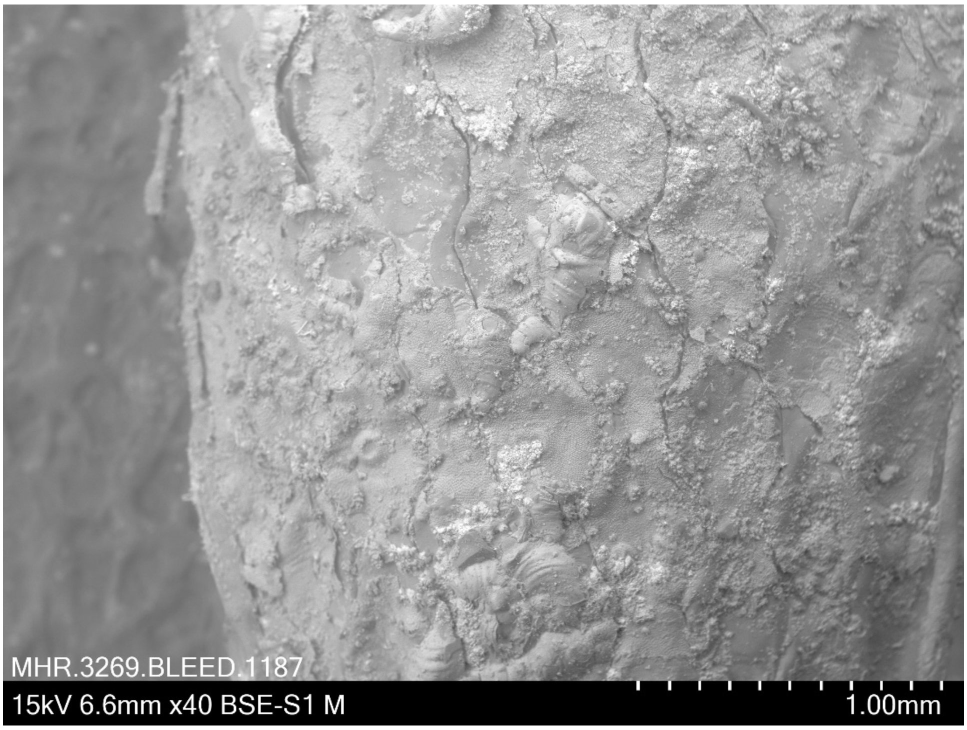


***Hippothoa* sp. BLEED 1187**
Coll. 1875? Pres. dried
Identified by I. Vigeland in 1970 (unregistered specimen from NHM Oslo)
Location: North Shields, England. No other metadata available.
<u>Taxonomic notes</u>: This specimen is reminiscent of *H. divaricata* Lamouroux, 1821 but the cauda is broader and a median carina on the frontal shield is not seen.

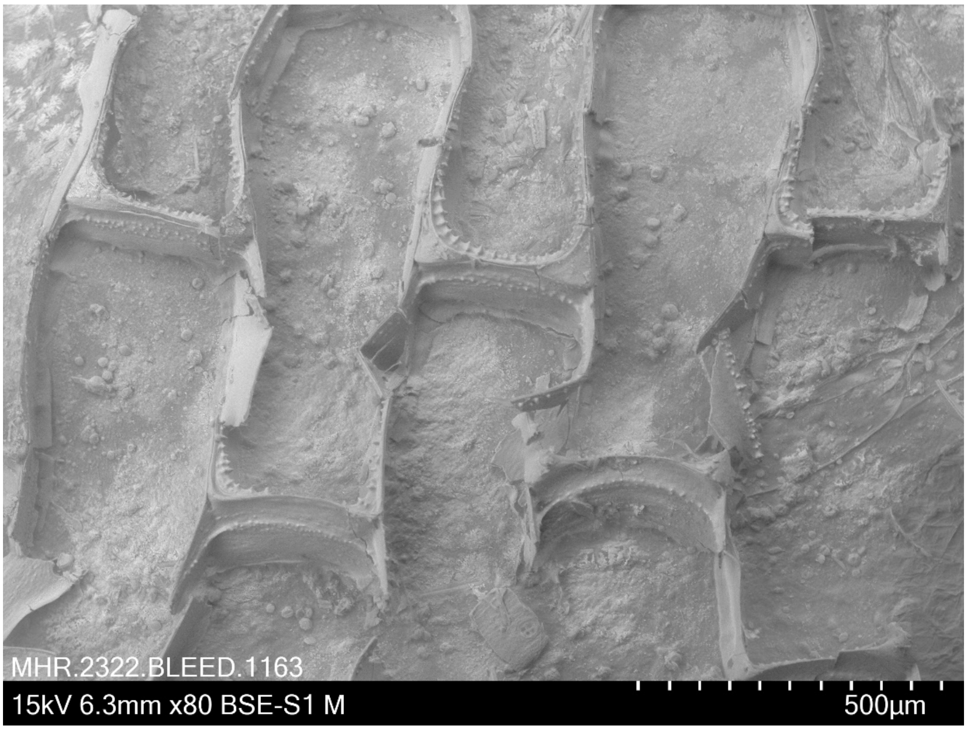


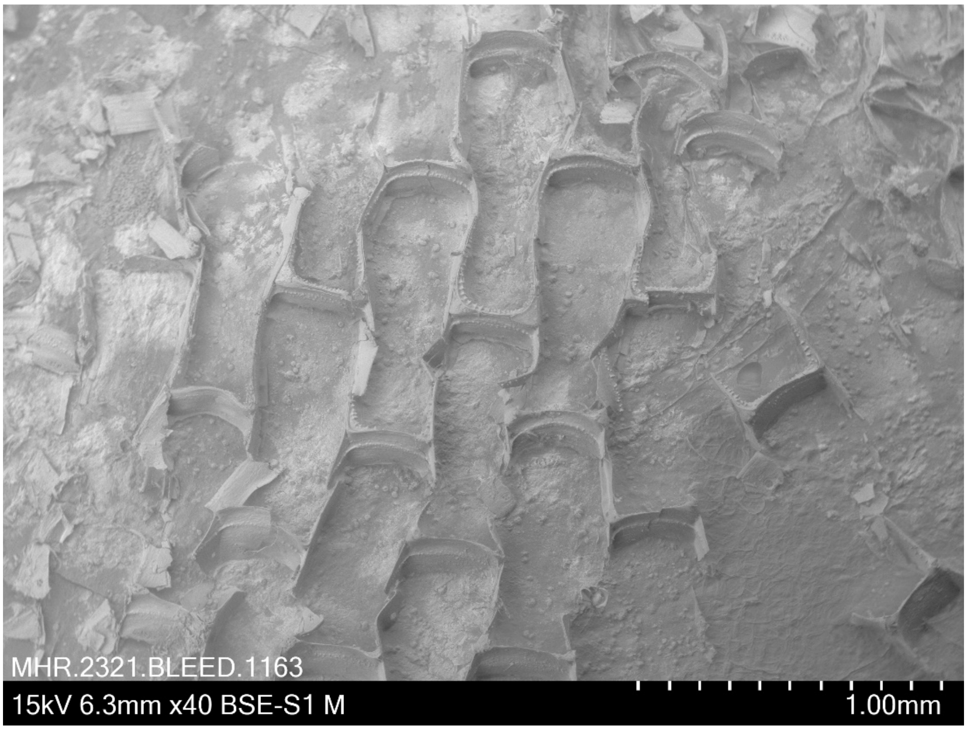


***Menbranipora membranacea* (Linnaeus, 1767) BLEED 1163**
Coll. 28 May 2018 (L.H. Liow) Pres. dried
Lat: 58.247445, Long: 11.404071, 15-45 m, off Kristineberg, Sweden

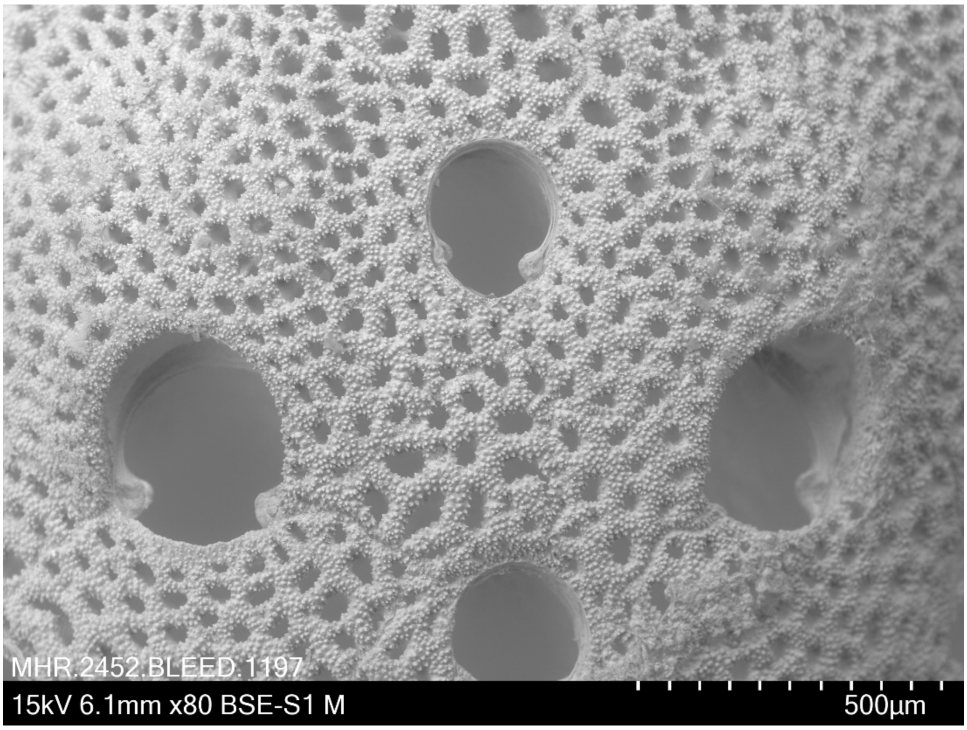


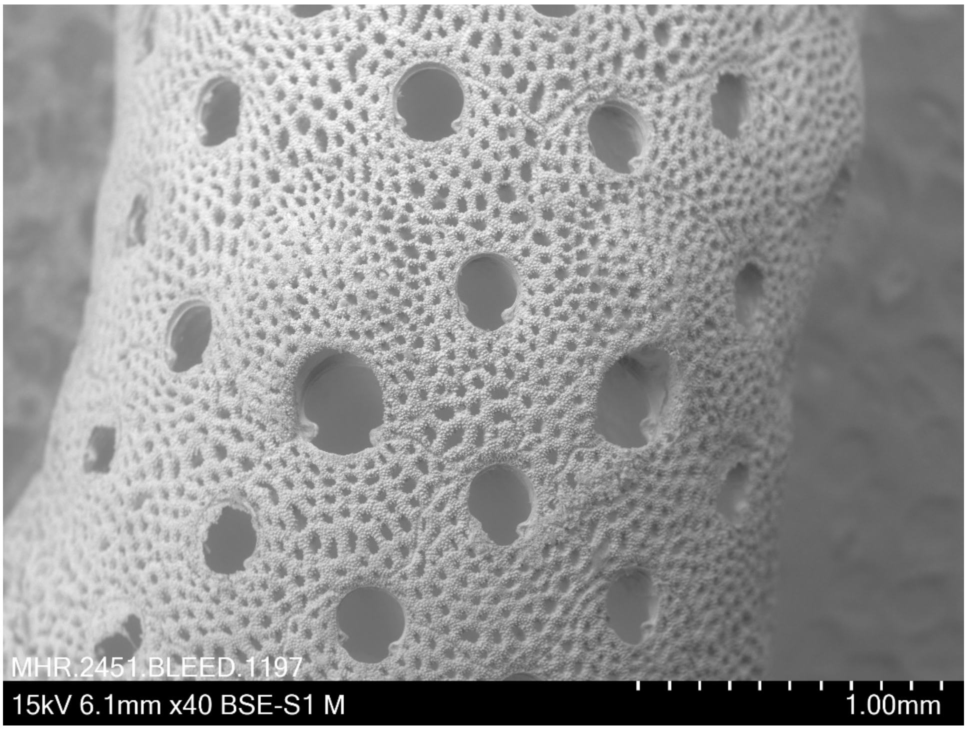


***Myriapora truncata*** **(Pallas, 1776) BLEED 1197**
Coll. 1871 Pres. dried
Identified by I. Vigeland in 1970 (unregistered specimen from NHM Oslo)
Location: Bay of Naples, Italy. No other metadata available.

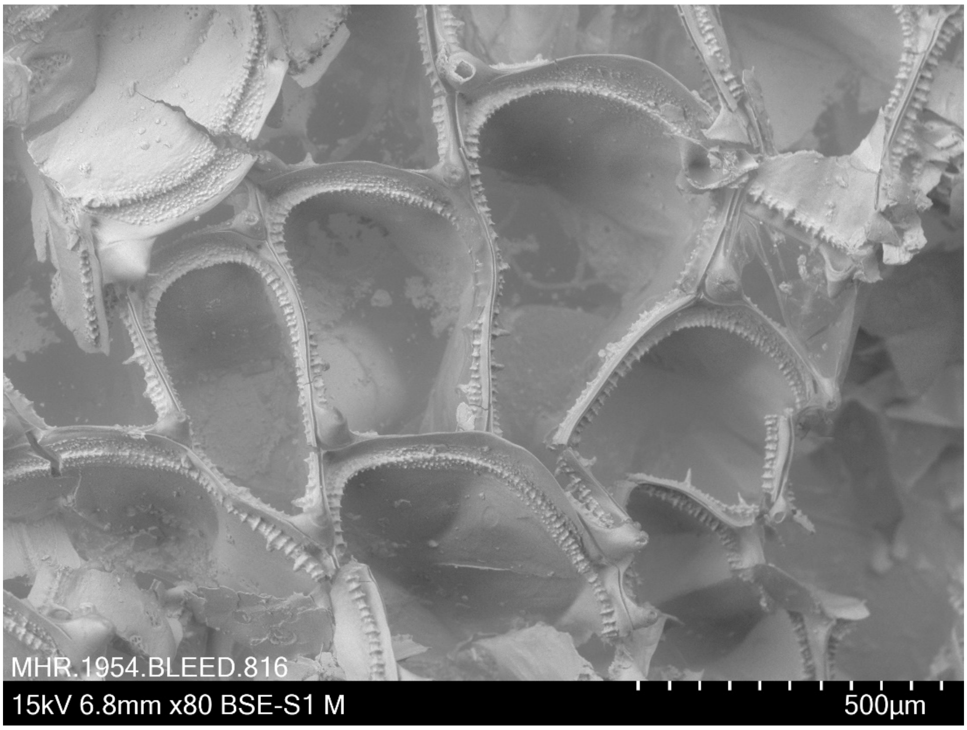


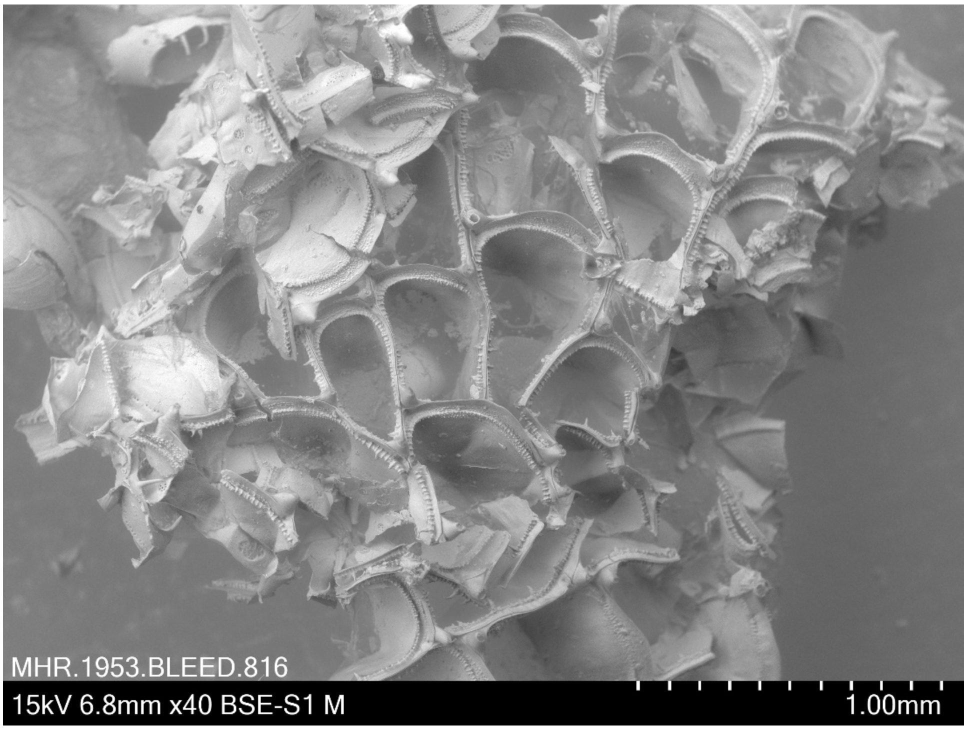


***Membranipora membranacea*** **(Linnaeus, 1767) BLEED 816**
Coll. 4 September 2018 (Maria Capa) Pres. ethanol
Lat: 63.439001, Long: 10.50548, Depth: intertidal, Trondheim, Norway
GRIFAS-S4

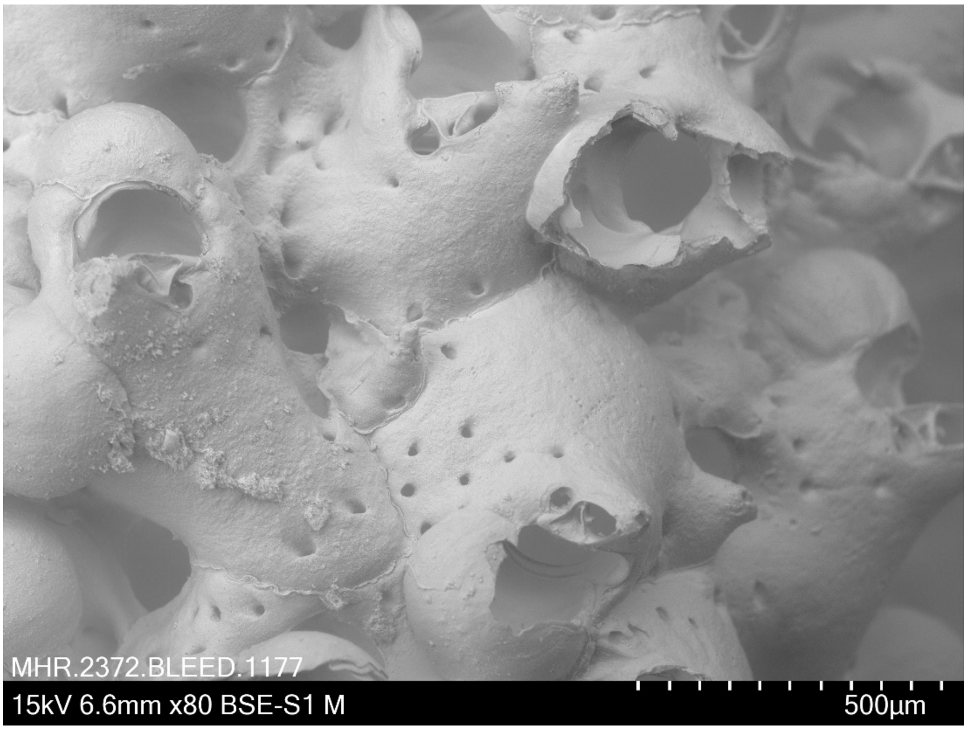


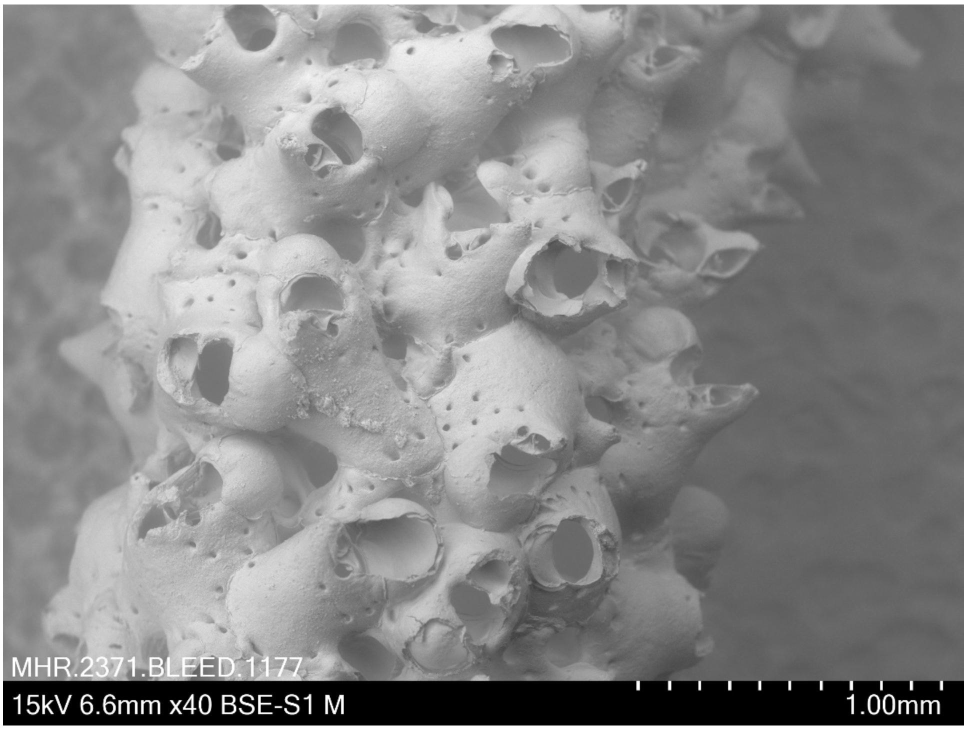


***Omalosecosa ramulosa* (Linnaeus, 1767) BLEED 1177**
Pres. dried
Identified by I. Vigeland in 1970 (unregistered specimen from NHM Oslo)
Location: Bergenfjord, Norway. No other metadata available.

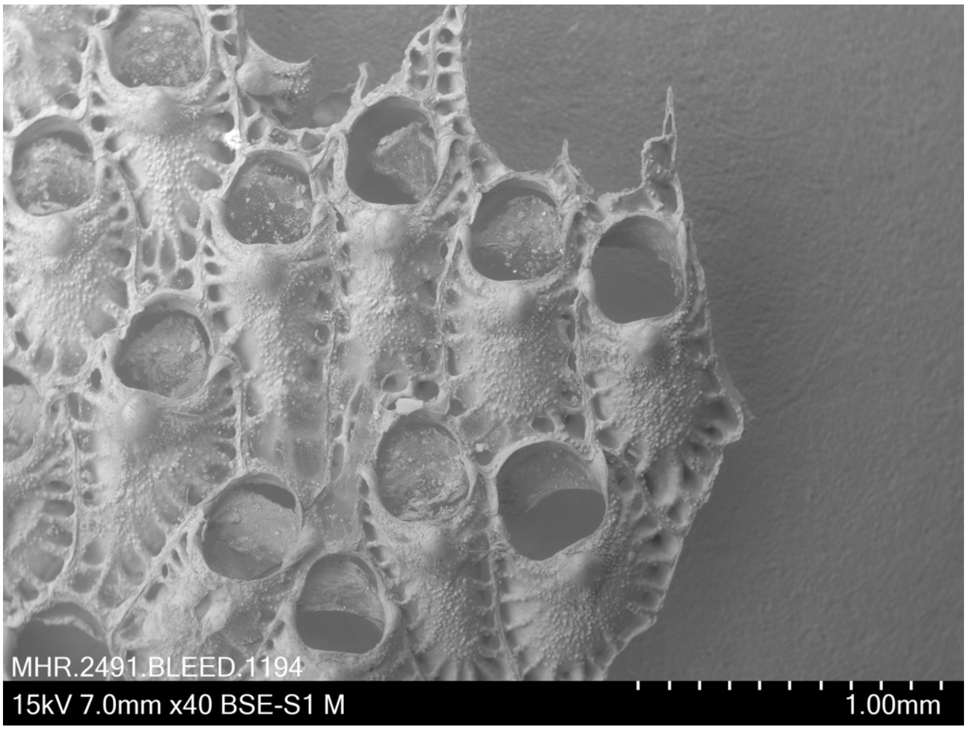


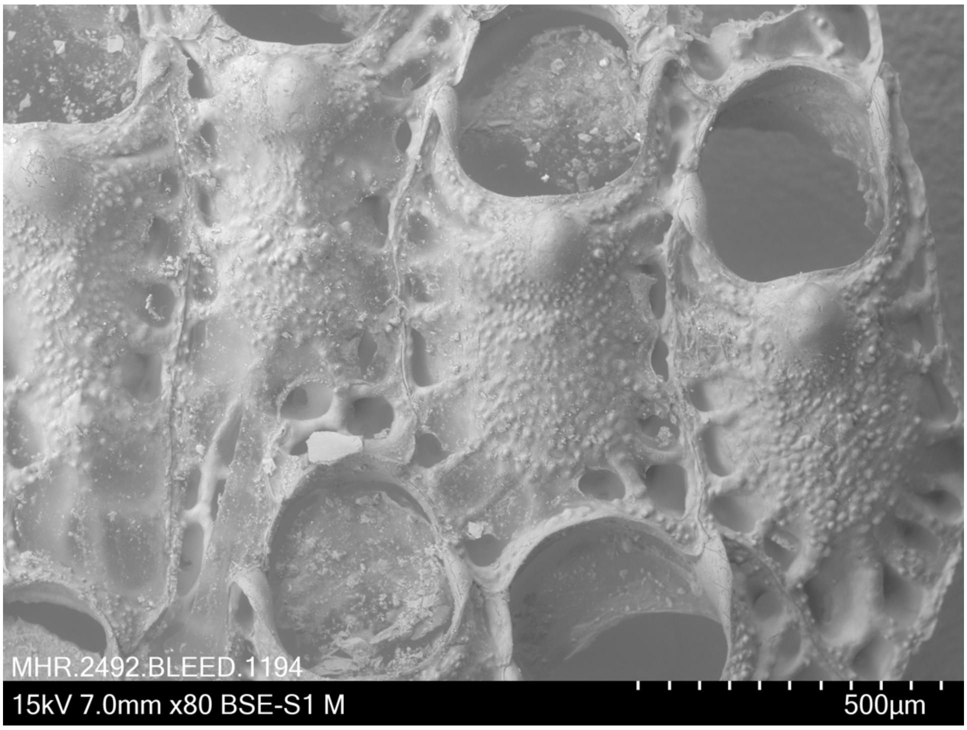


***Oshurkovia littoralis* (Hastings, 1944) BLEED 1194**
Pres. dried
Identified by I. Vigeland in 1970 (unregistered specimen from NHM Oslo)
Location: Britain. No other metadata available.

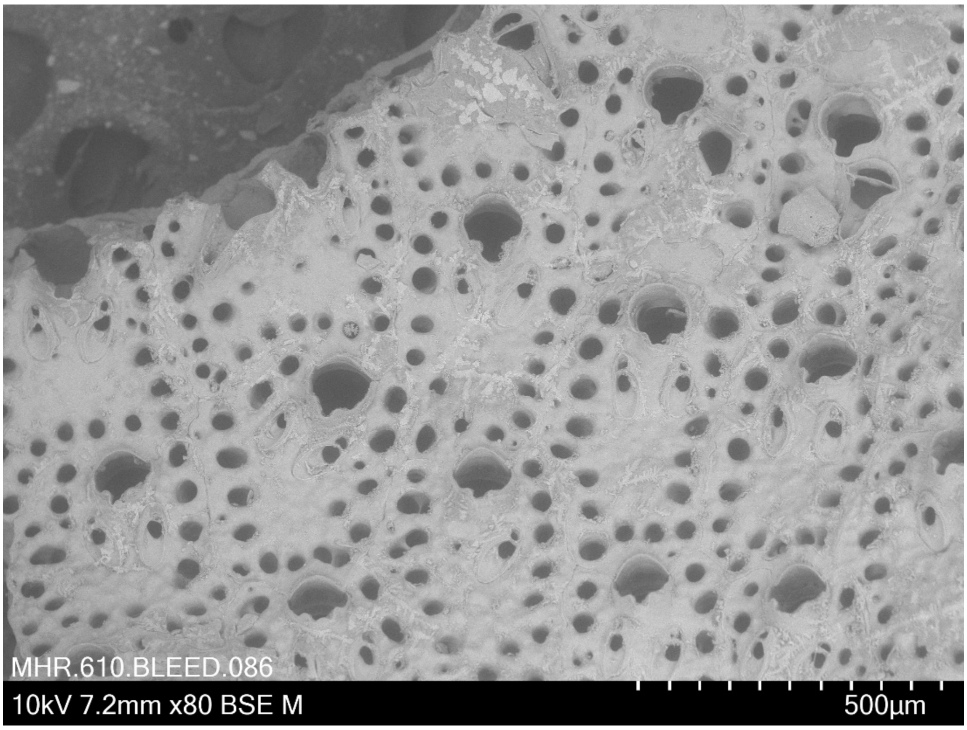


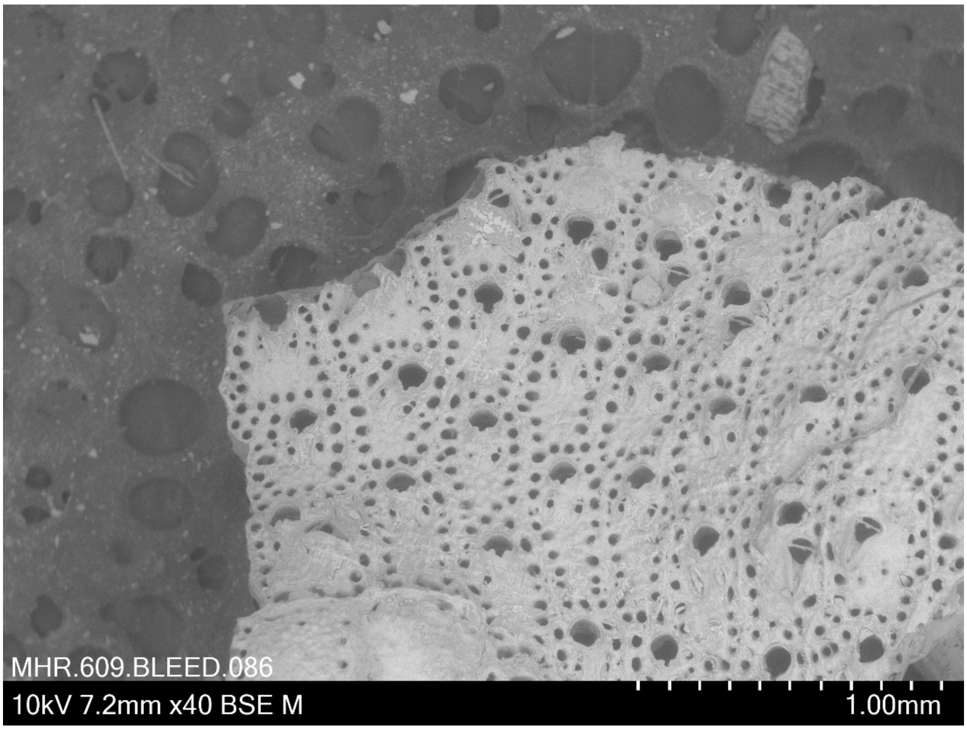


***Parasmittina aotea* (Brown, 1952) BLEED 86**
Coll. 18.2.2010 Pres. ethanol
Lat: -47.933, Long: 168.667, Depth: 146-149 m, New Zealand
SOP Station ID: TRIP3072/1 NIWA 61918

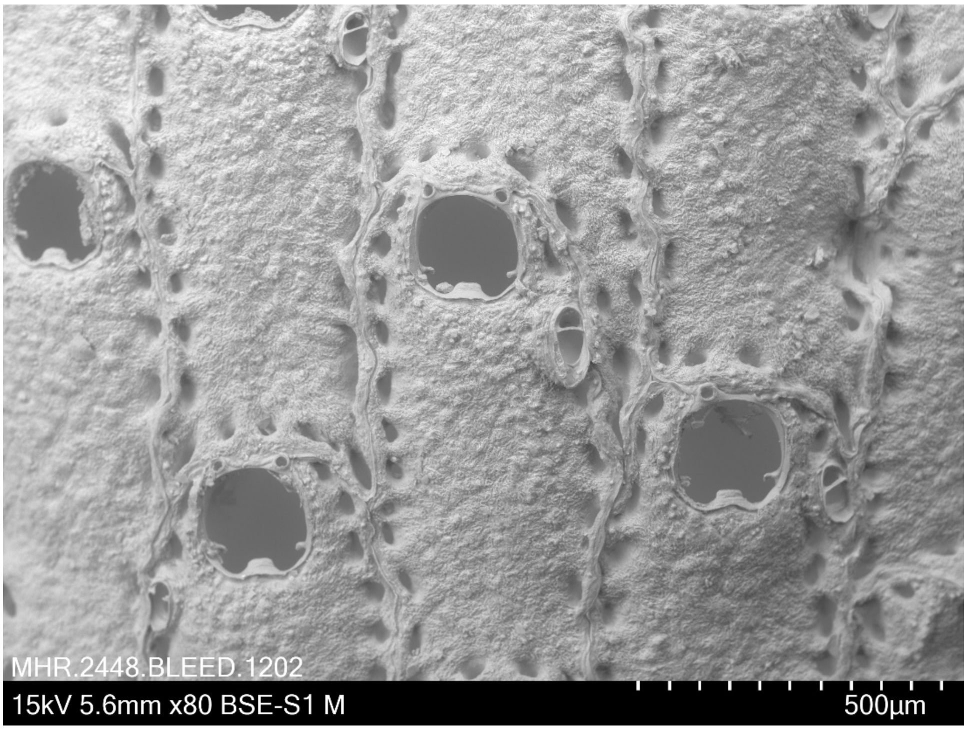


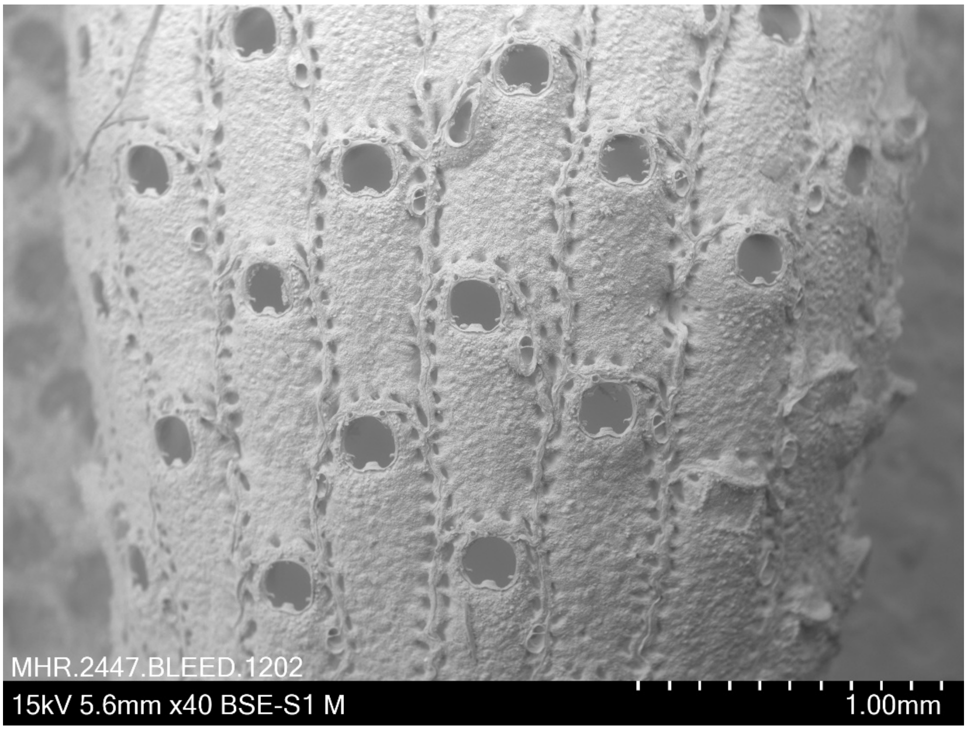


***Parasmittina jeffreysi* Norman, 1876 BLEED 1202**
Collection date 16.8.1901 Pres. dried
unregistered FRAM II specimen from NHM Oslo
Location: Artic (Greenland to Canada). No other metadata available.

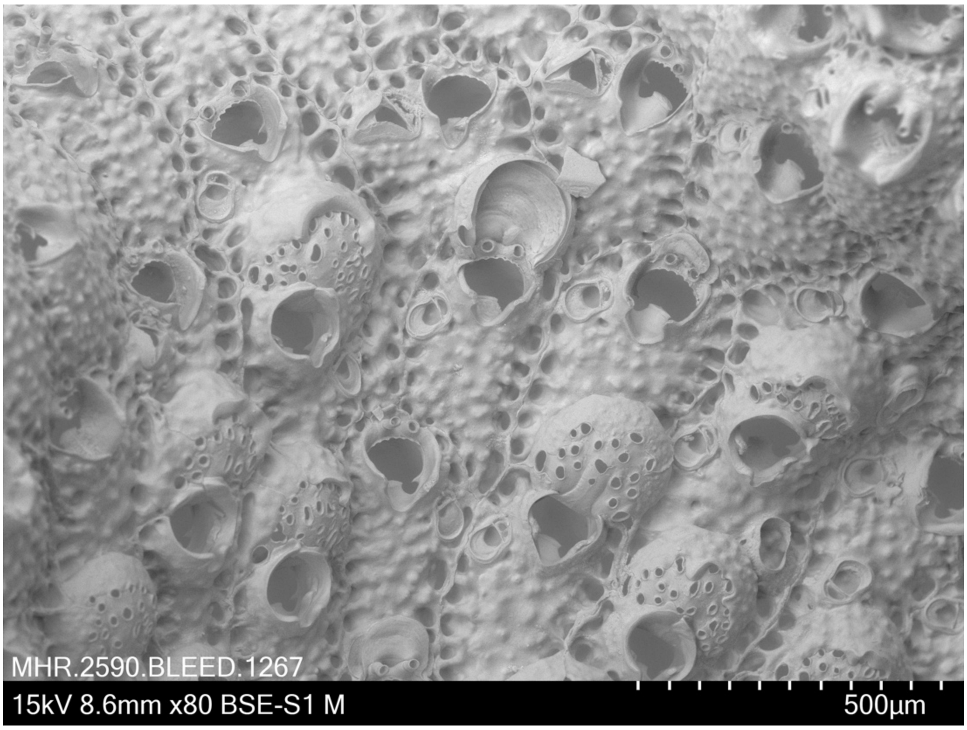


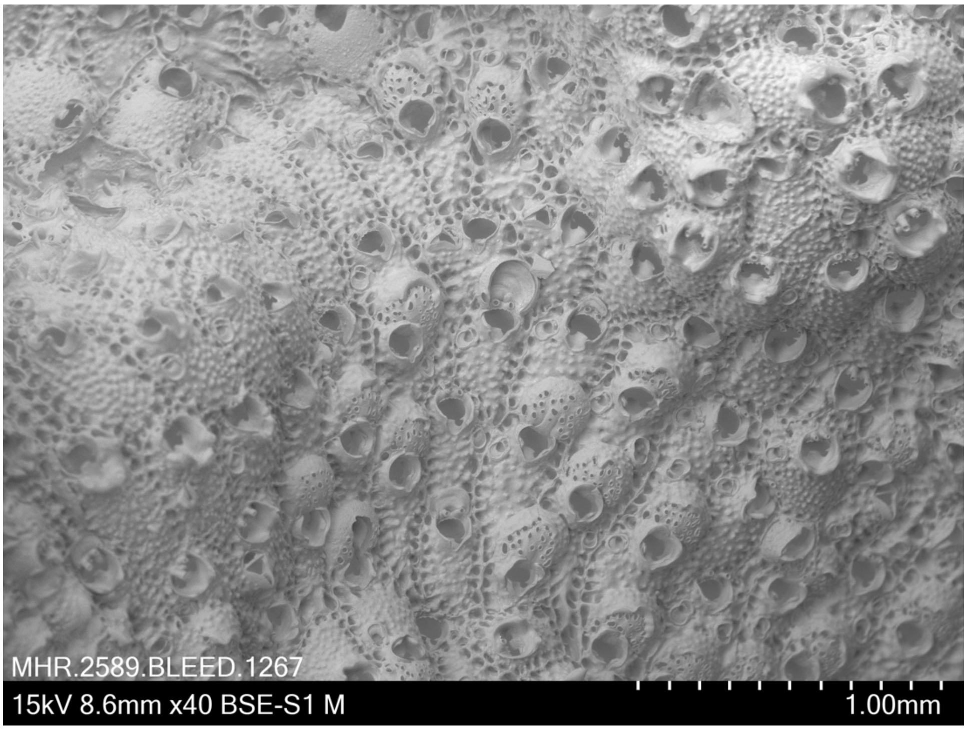


***Parasmittina solenosmilioides*** Hayward & Parker, 1994. **BLEED 1267**
Coll. 16.3.2019 (A. M. Smith, D. P. Gordon) Pres. ethanol
Lat: -35.01, Long: 173.92, Depth: 17 m, SW of Motukawanui Island, New Zealand
Notes: This species is not a native species in New Zealand.

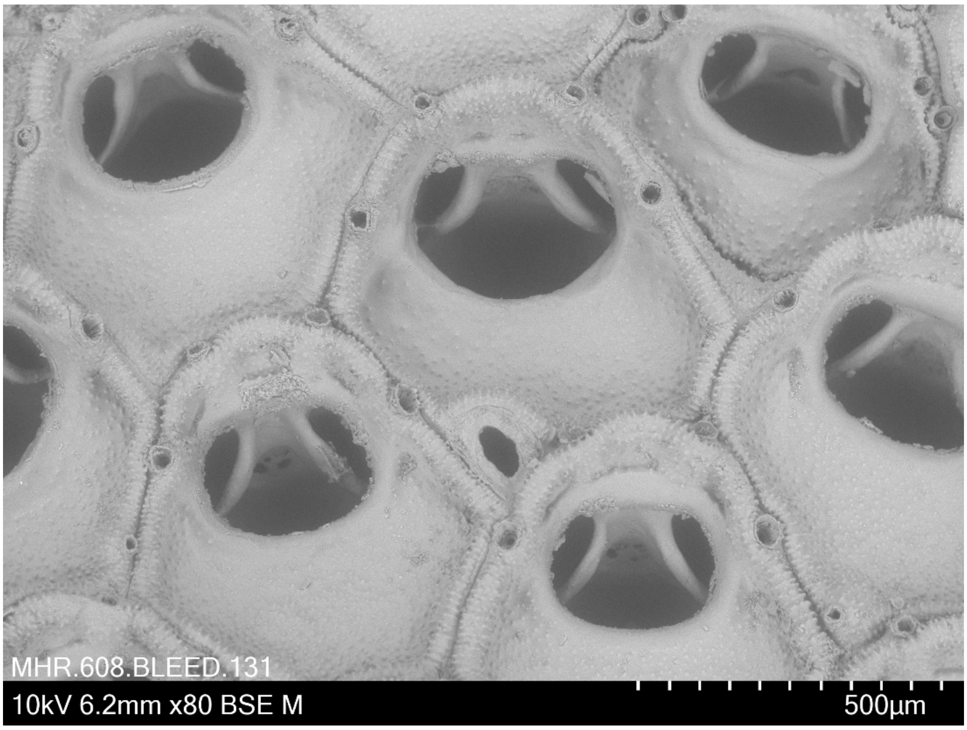


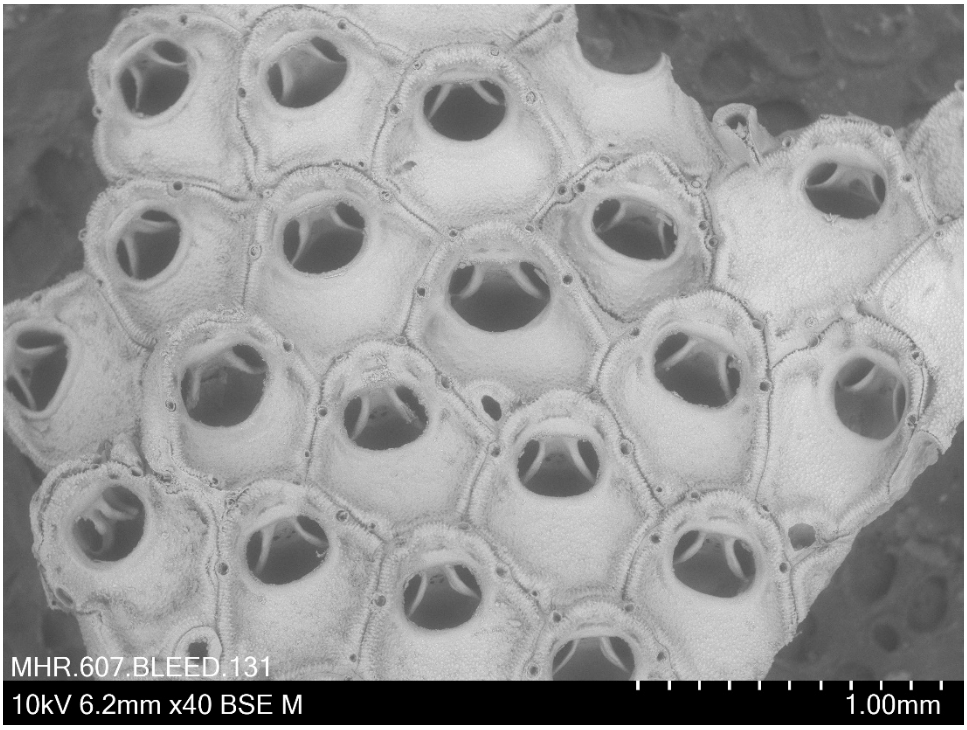


***Patsyella acanthodes* Gordon, 1982 BLEED 131**
Coll. 12.4.2011 Pres. ethanol
Lat: --46.28333, Long: 166.133333, Depth: ?, New Zealand
SOP Station ID: TRIP3298/36 NIWA 69631

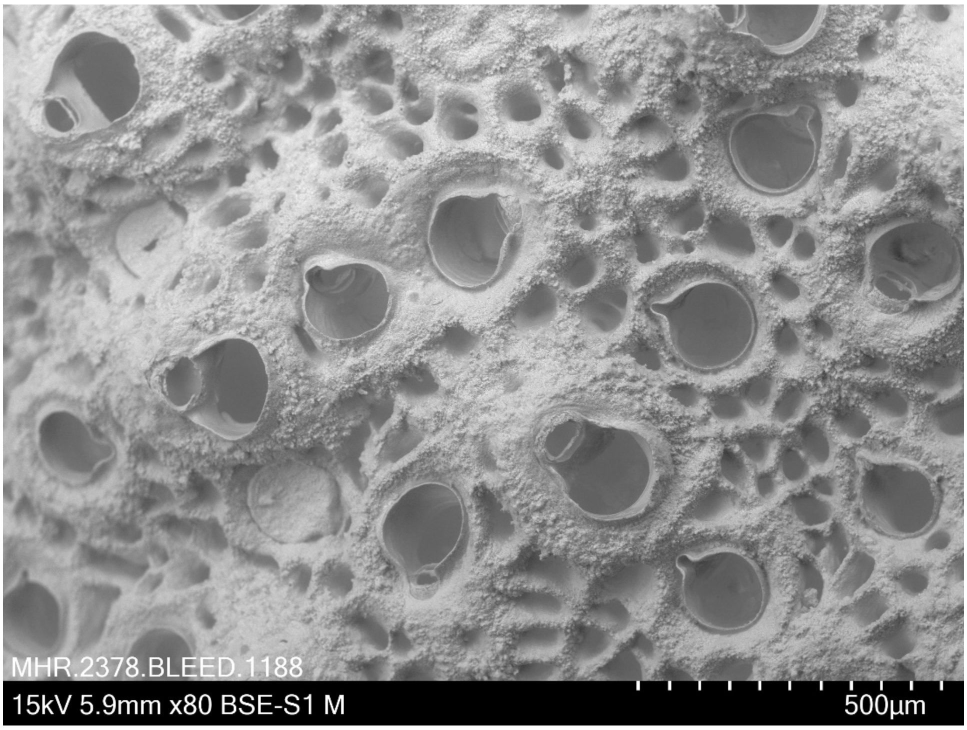


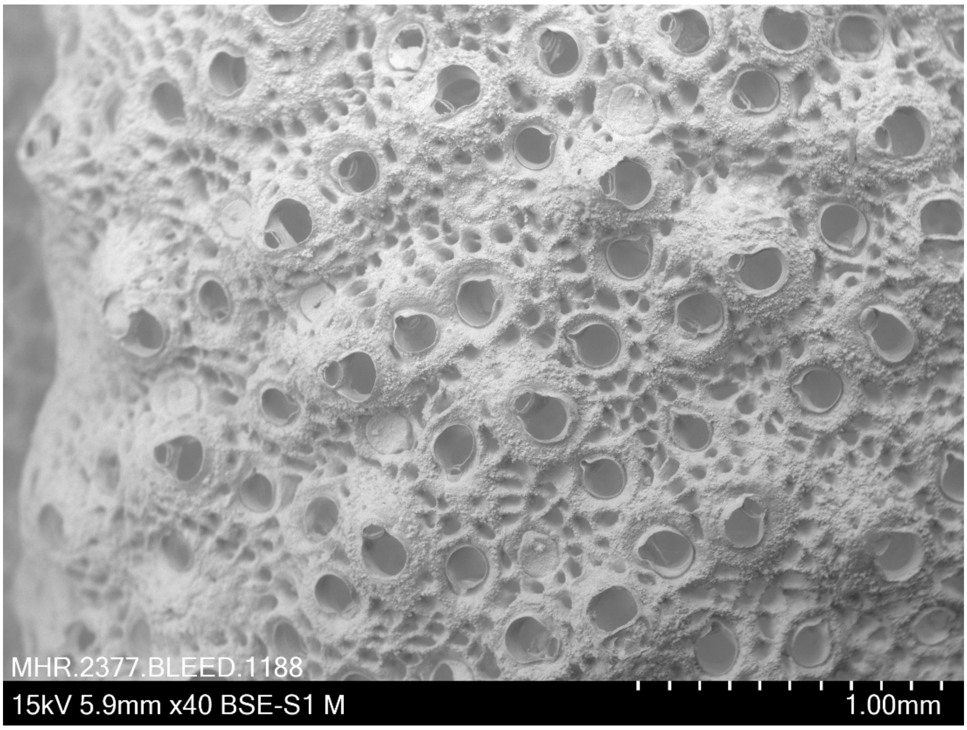


***Porella compressa* (Sowerby, 1805) BLEED 1188**
Pres. dried
Identified by I. Vigeland in 1970 (unregistered specimen from NHM Oslo)
Location: Tjøtta, Norland, Norway. No other metadata available.

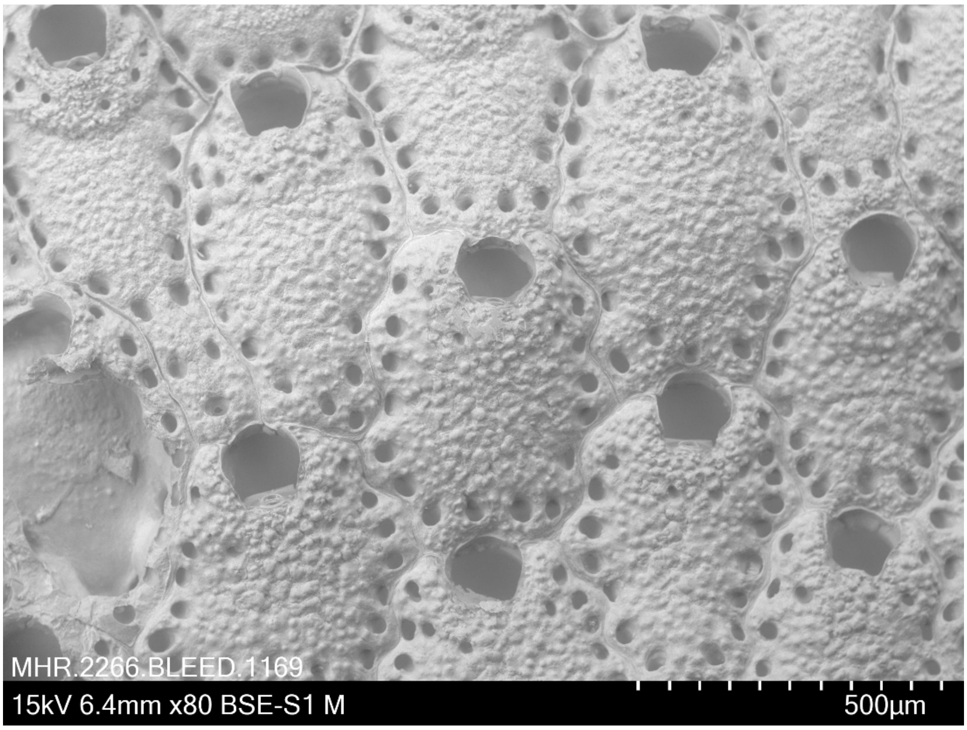


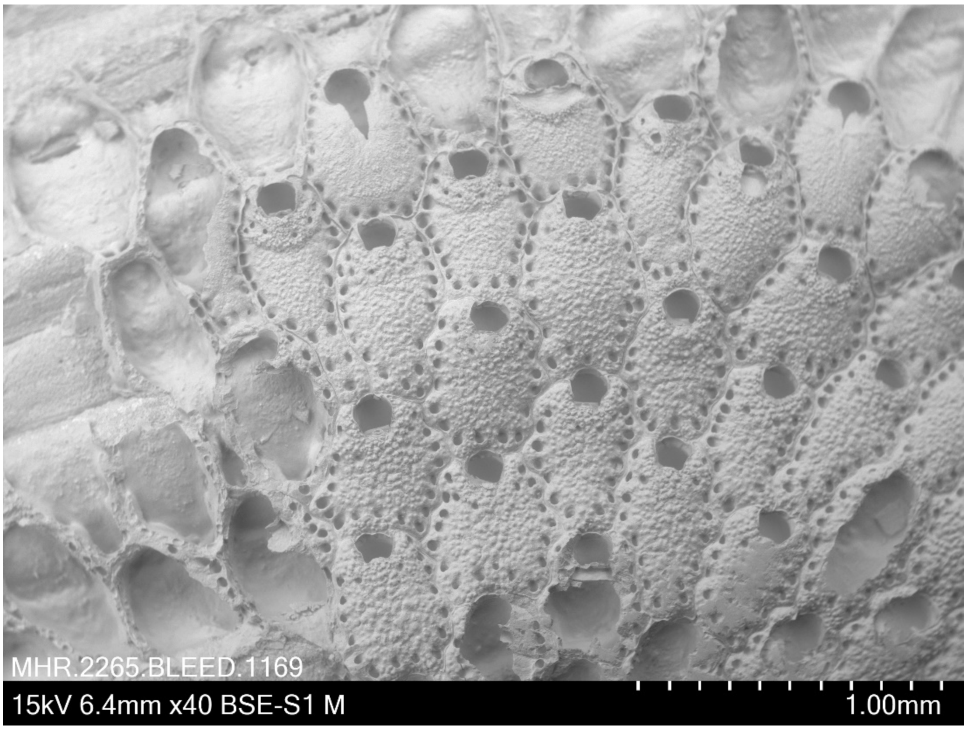


***Porella concinna* (Busk, 1854) BLEED 1169**
Coll. 28 May 2018 (L.H. Liow) Pres. dried
Lat: 58.247, Long: 11.404, 15-45 m, off Kristineberg, Sweden

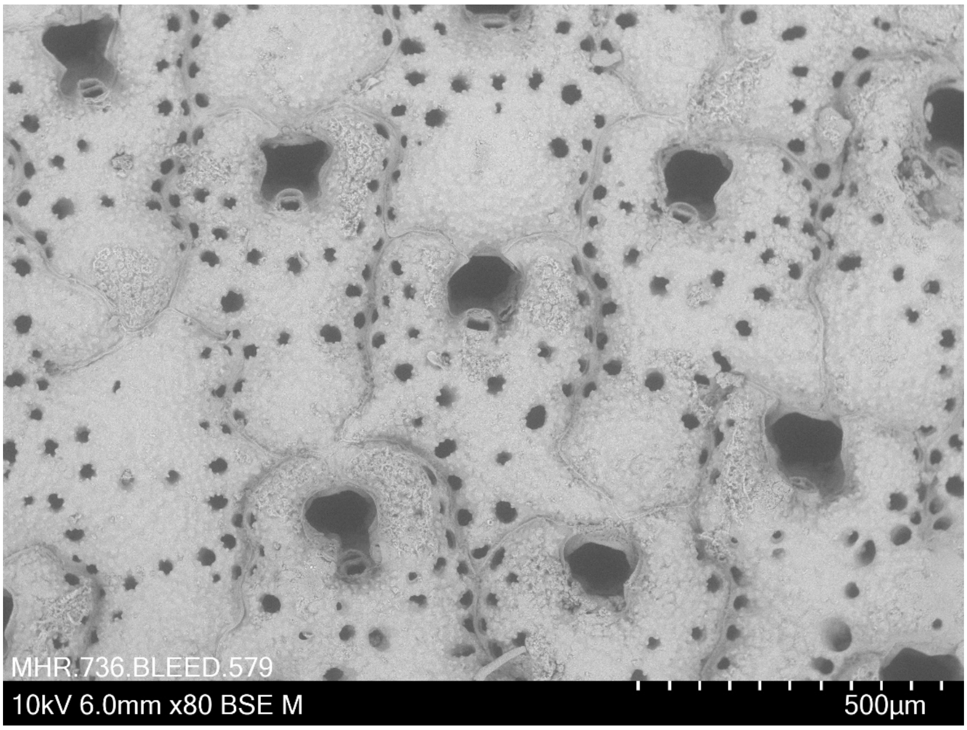


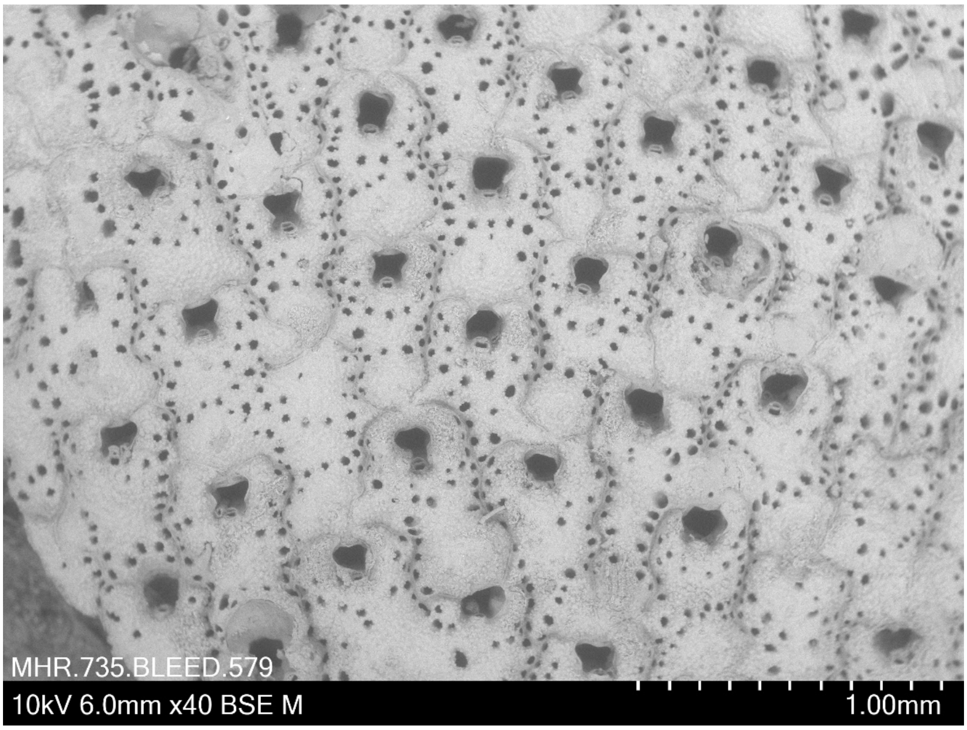


***Porella concinna* (Busk, 1854) BLEED 579**
Coll. 29 May 2018 (M. Obst, R.J.S. Orr, L.H.Liow) Pres. ethanol
Lat: 58.219, Long: 11.410, 20-40 m, off Kristineberg, Sweden

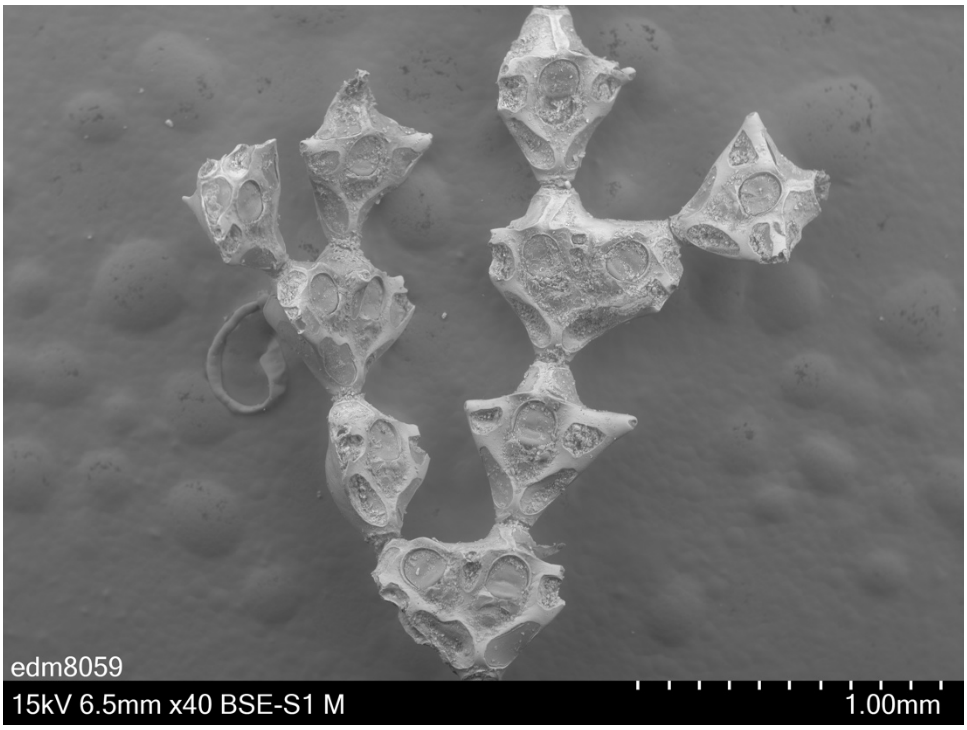


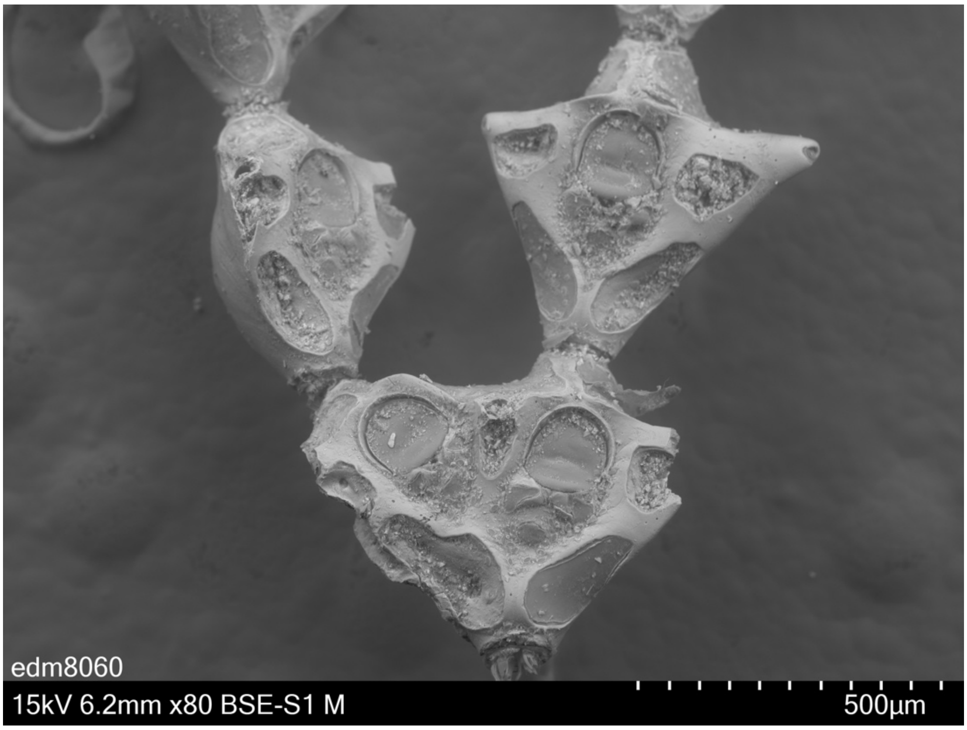


***Pterocella scutella*** **Gordon, 1982 BLEED 104**
Coll. 20.5.2015 (D.P. Gordon) Pres. ethanol
Lat: -40.900, Long: 176.217, Depth: intertidal, Castlepoint, Wairarapa, New Zealand

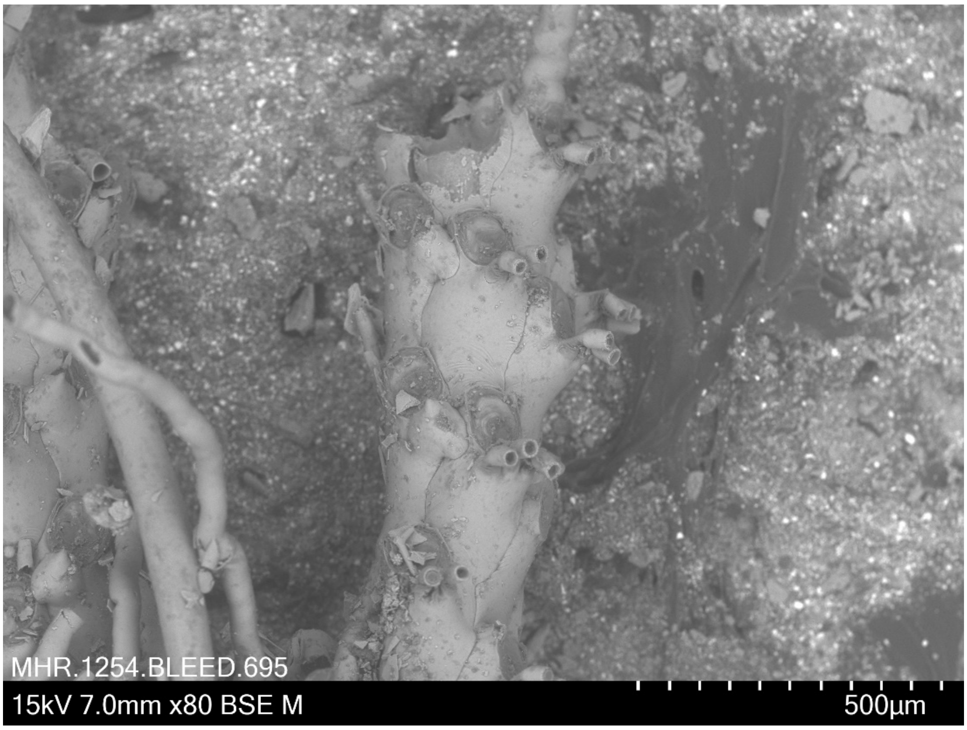


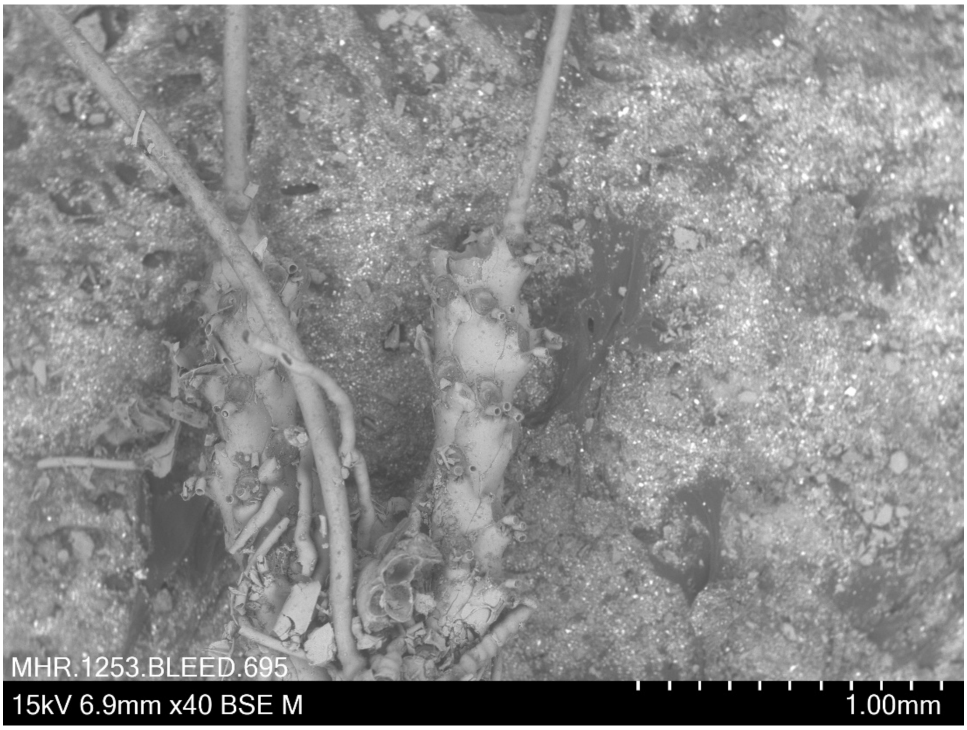


***Rhabdozoum wilsoni*** **Hincks, 1882 BLEED 695**
Coll. 8.4.2018 (A. M. Smith, D.P. Gordon, H. Mello) Pres. ethanol
Lat: - 47.158, Long: 168.011, Depth: 72 m, off White Rock, Stewart Island, New Zealand

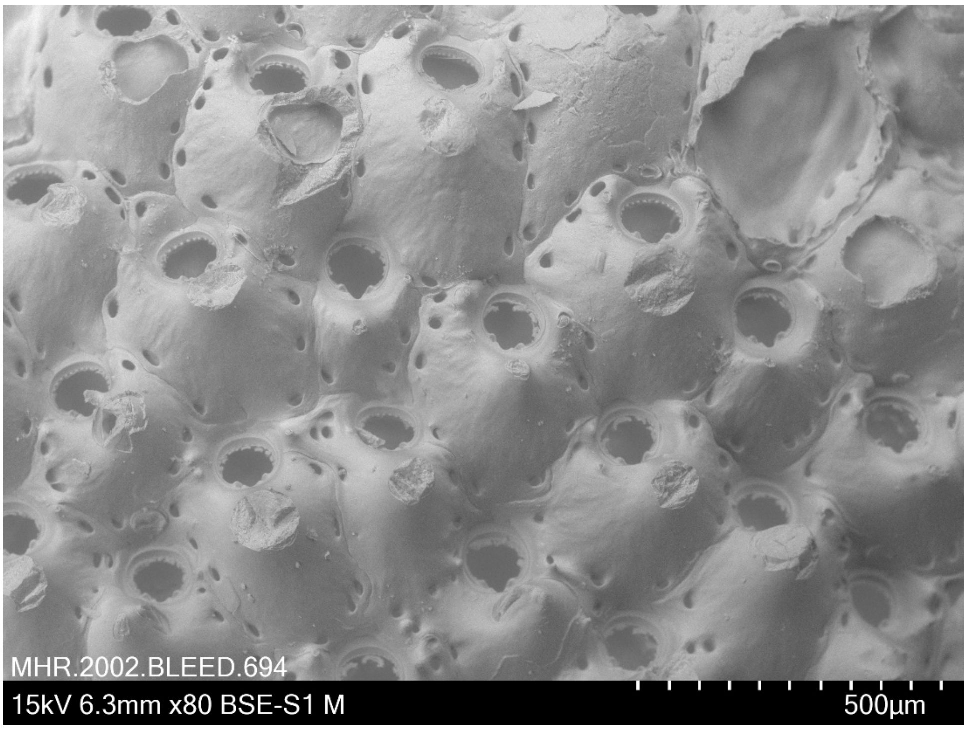


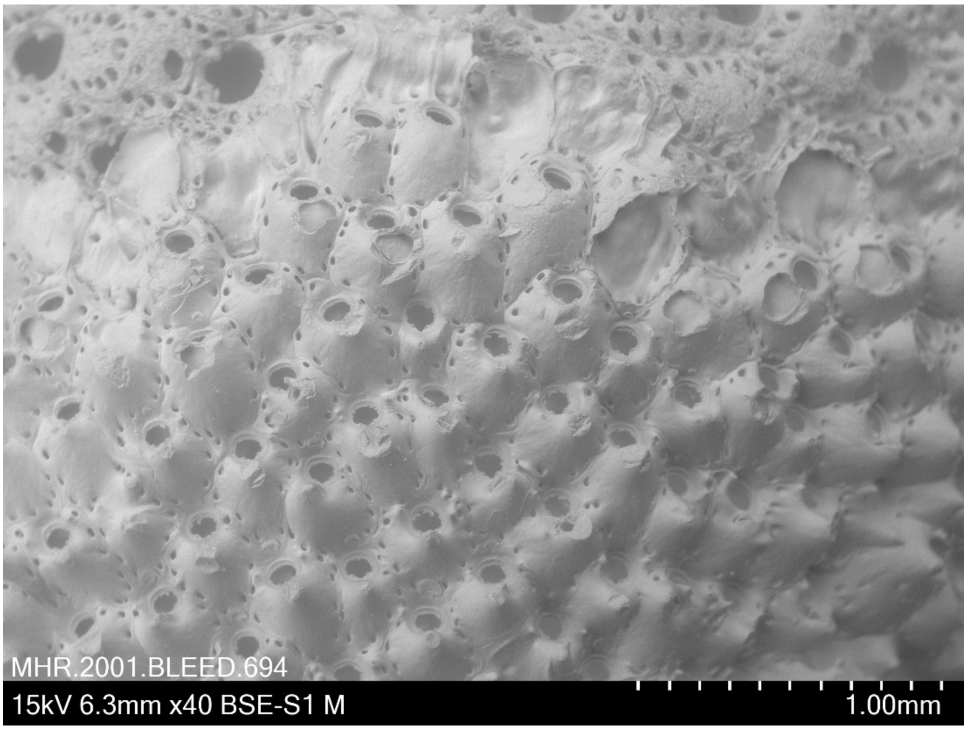


***Rhynchozoon angulatum* (Leveinsen, 1909) BLEED 694**
Coll. 8.4.2018 (A. M. Smith, D.P. Gordon, H. Mello) Pres. ethanol
Lat: - 46.662, Long: 168.386, Depth: 22 m, W of Dog Island, New Zealand
Taxonomic notes: The development of the avicularia of this specimen is weak relative for a *R. angulatum* and also lacks aviculiferous 'mucronate process' above the sinus, such as is found in *R. zealandicum* (and many other *Rhynchozoon* species).

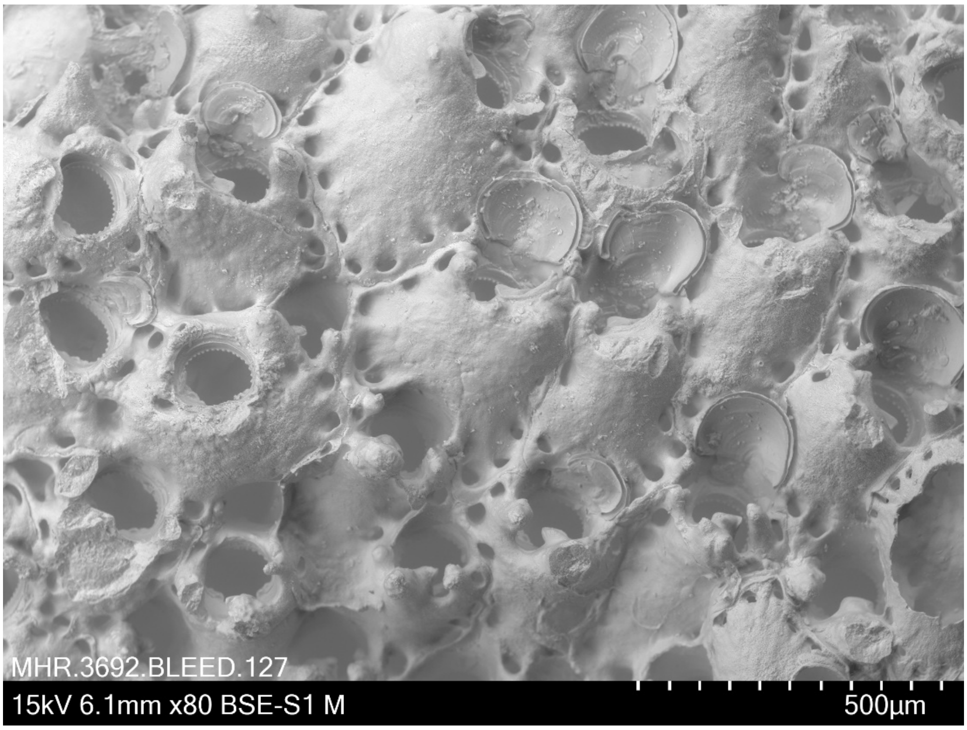


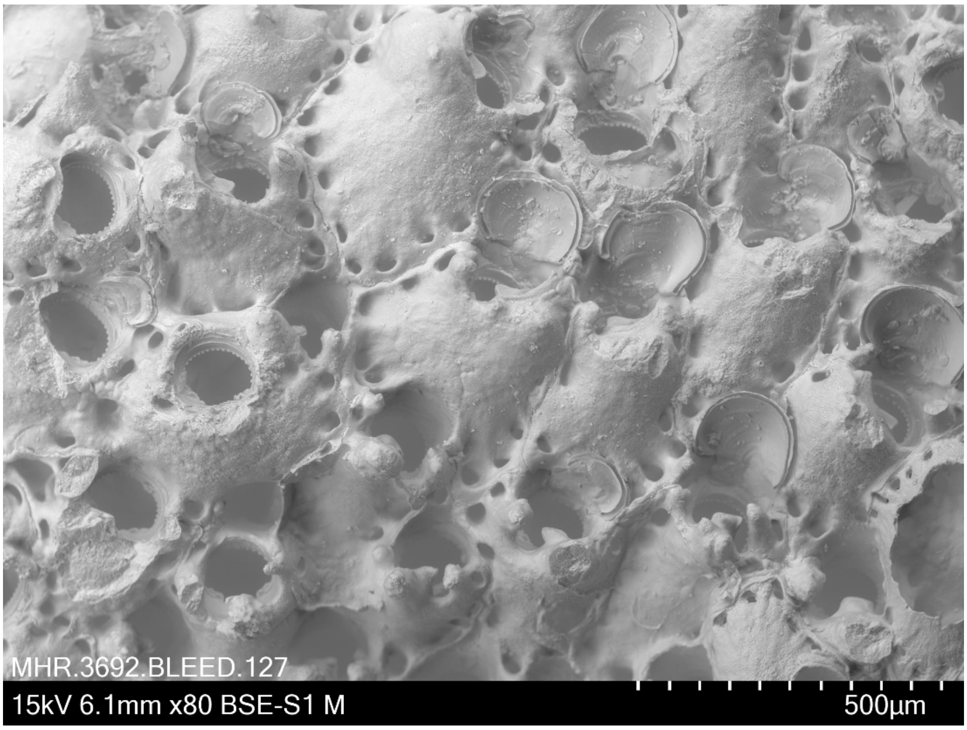


***Rhynchozoon zealandicum* Gordon, 2009 BLEED 127**
Coll. February 2015 (D. P. Gordon) Pres. ethanol
Lat: -41.283, Long: 174.767, Depth 0-1m, New Zealand
Z16029 NIWA 99667

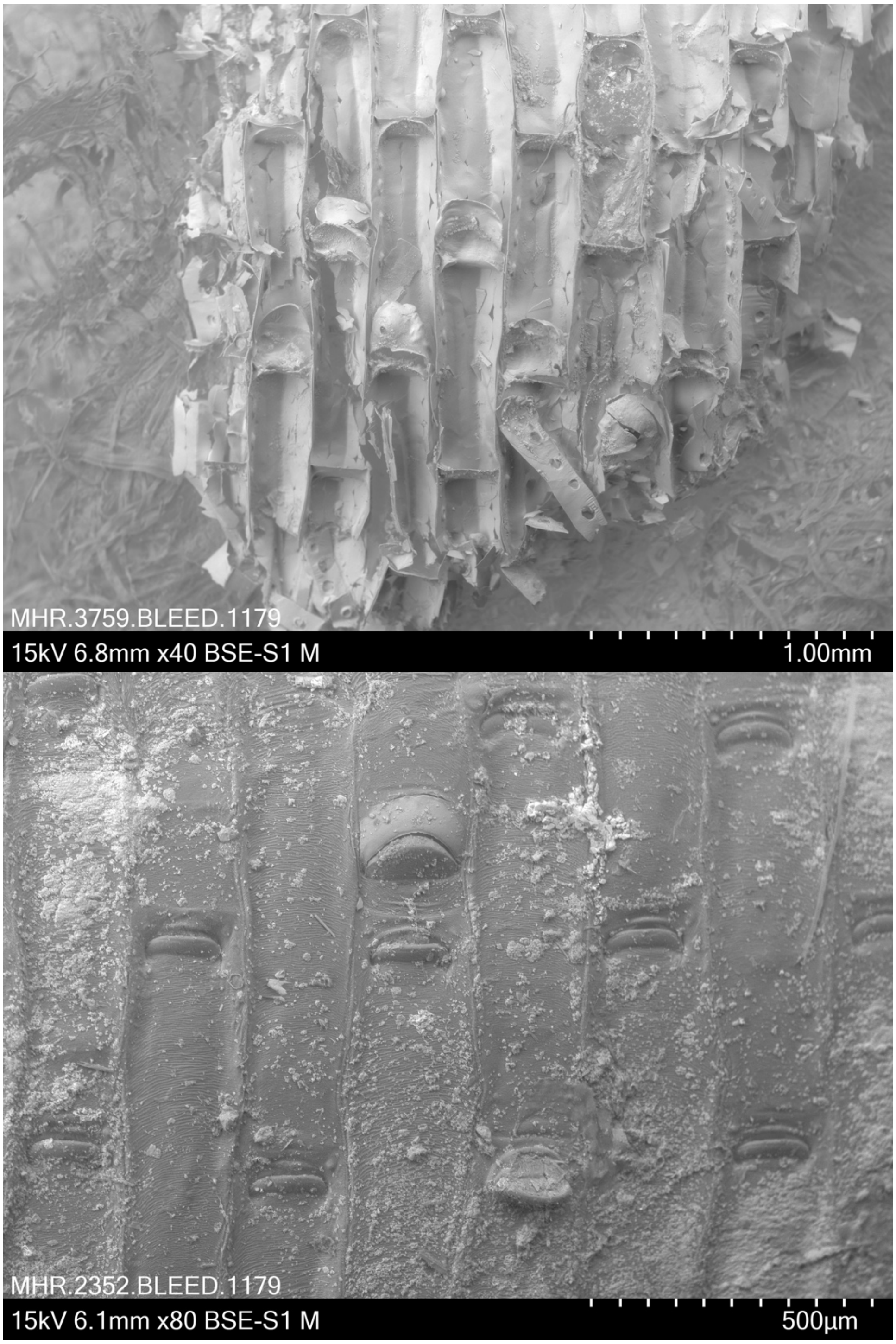


***Securiflustra securifrons*** **(Pallas, 1766) BLEED 1179**
Pres. dried
Identified by I. Vigeland in 1970 (unregistered specimen from NHM Oslo)
Location: Finnmark, Norway. No other metadata available.

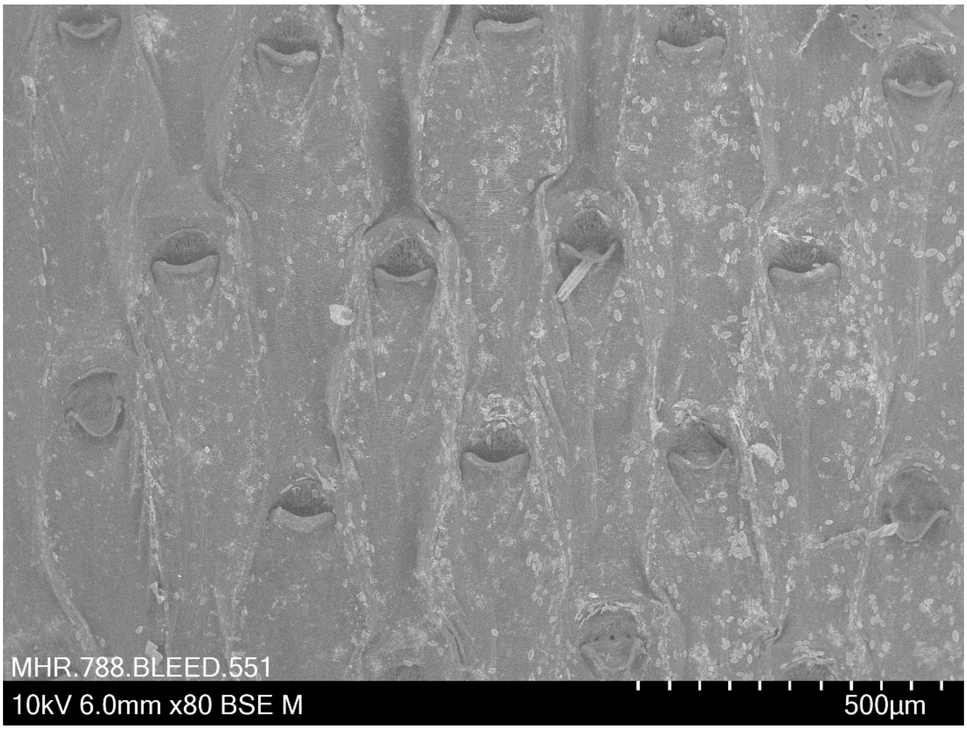


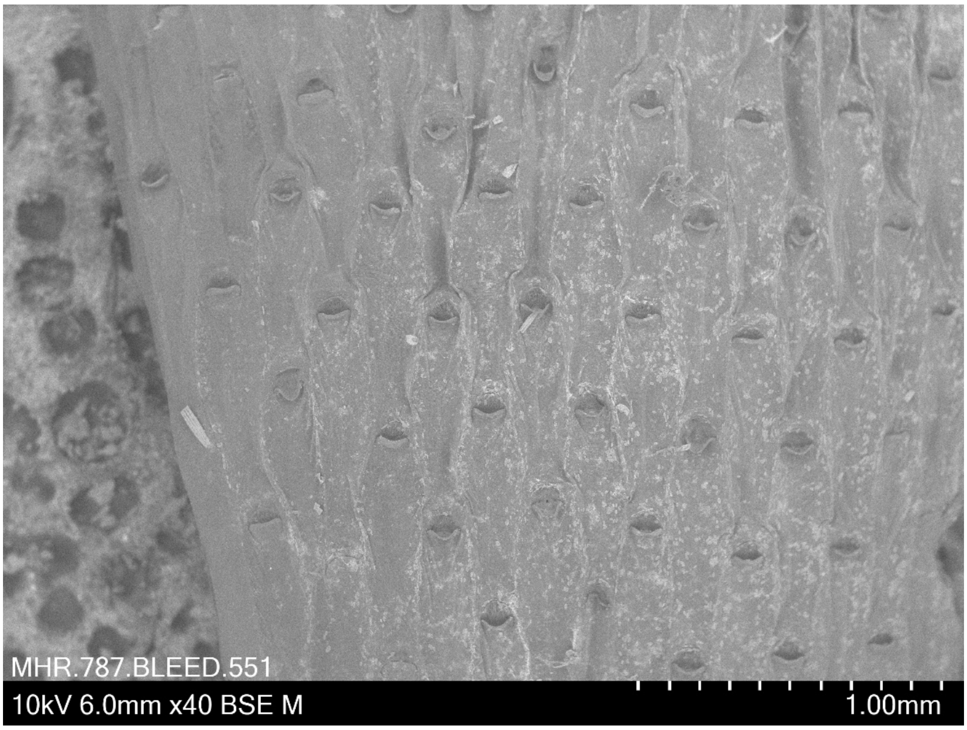


***Securiflustra securifrons* (Pallas, 1766) BLEED 551**
Coll. 28 May 2018 (M. Obst, R.J.S. Orr, L.H.Liow) Pres. ethanol
Lat: 58.247, Long: 11.4041, 15-45 m, off Kristineberg, Sweden

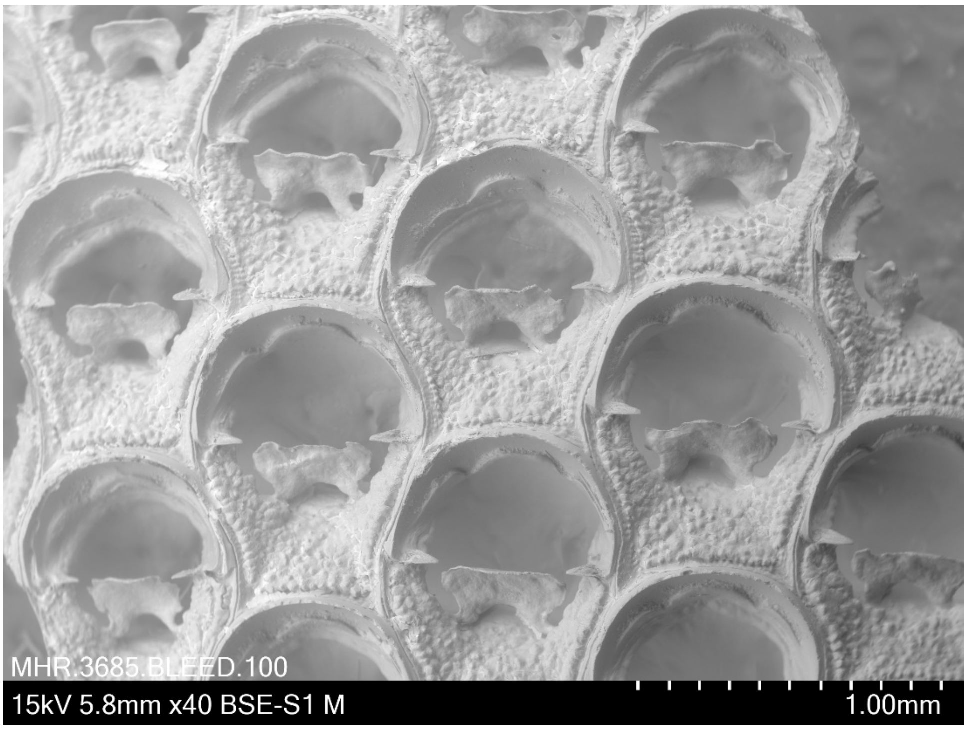


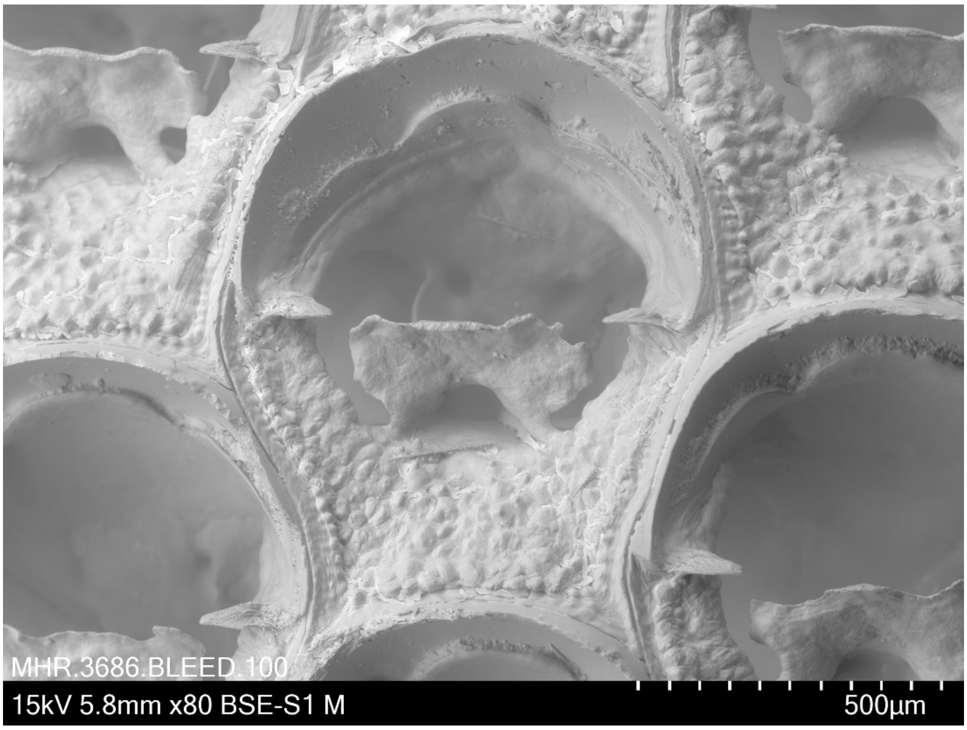


***Steginoporella perplexa*** **Livingstone, 1929 BLEED 100**
Coll. 28.3.2011 Pres. ethanol
Lat: -33.988 Long: 171.751, Depth170-174m, New Zealand
Oceans Survey 2020 Station ID: TAN1105/43 NIWA 73295

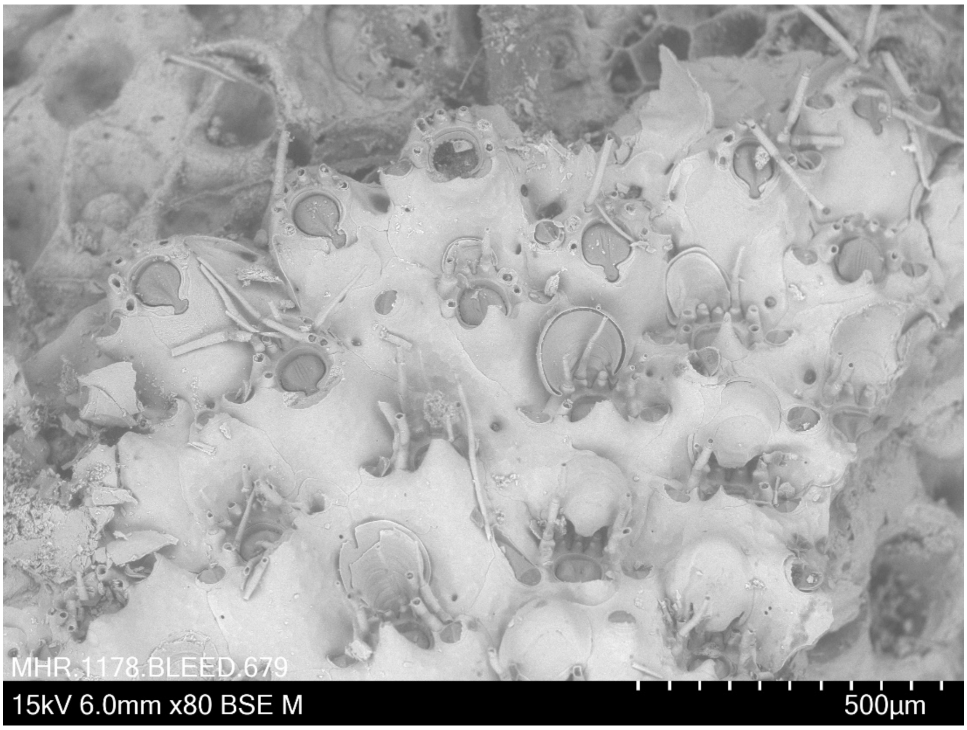


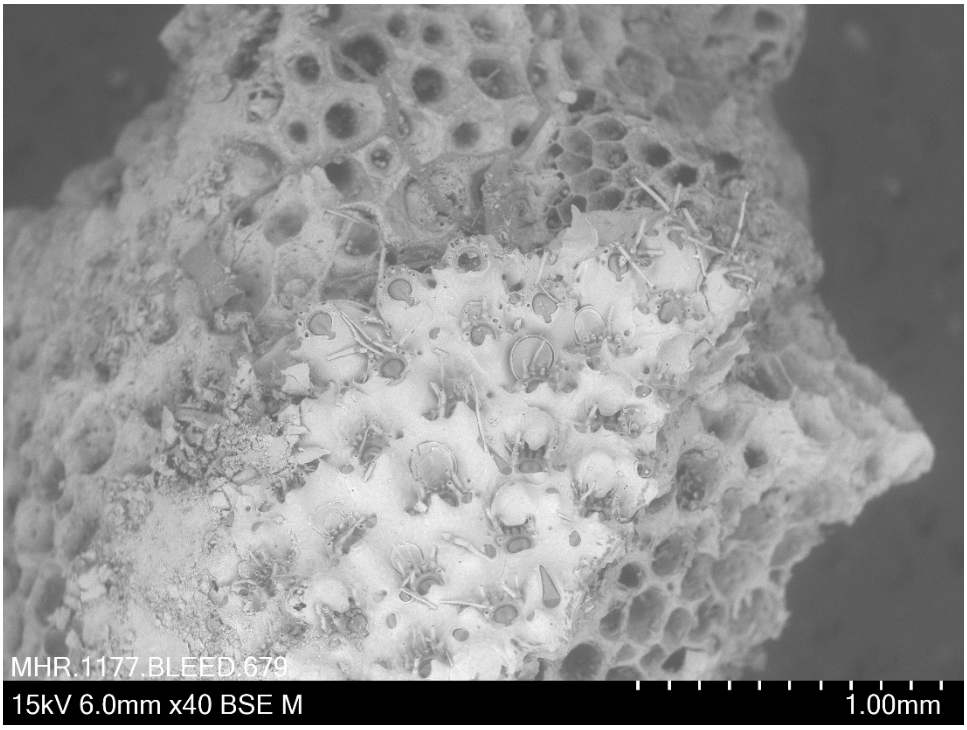


***Stephanollona scintillans* (Hincks, 1885) BLEED 679**
Coll. 8.4.2018 (A. M. Smith, D.P. Gordon, H. Mello) Pres. ethanol
Lat: -47.136, Long: 168.189, Depth: 77m, New Owen Island, New Zealand

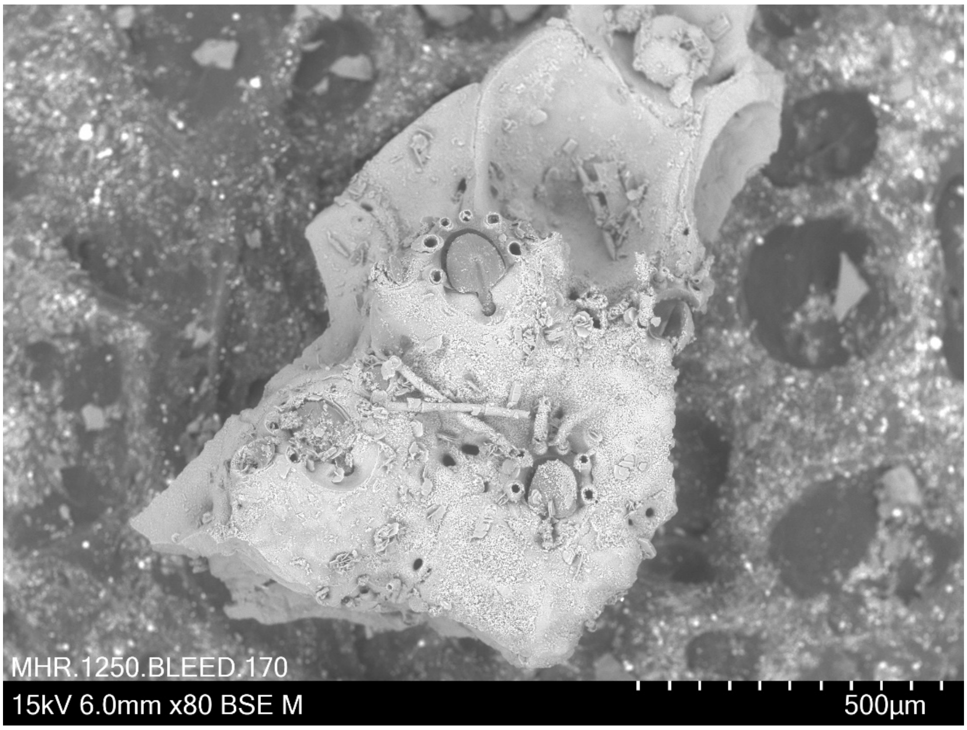


***Stephanollona* sp. BLEED 170**
Coll. 8 July 2009 Pres. ethanol
Lat: 35.004, Long: 174.056 Depth: 105 m, New Zealand
Oceans Survey 2020 Station ID: TAN0906/65 NIWA 55263

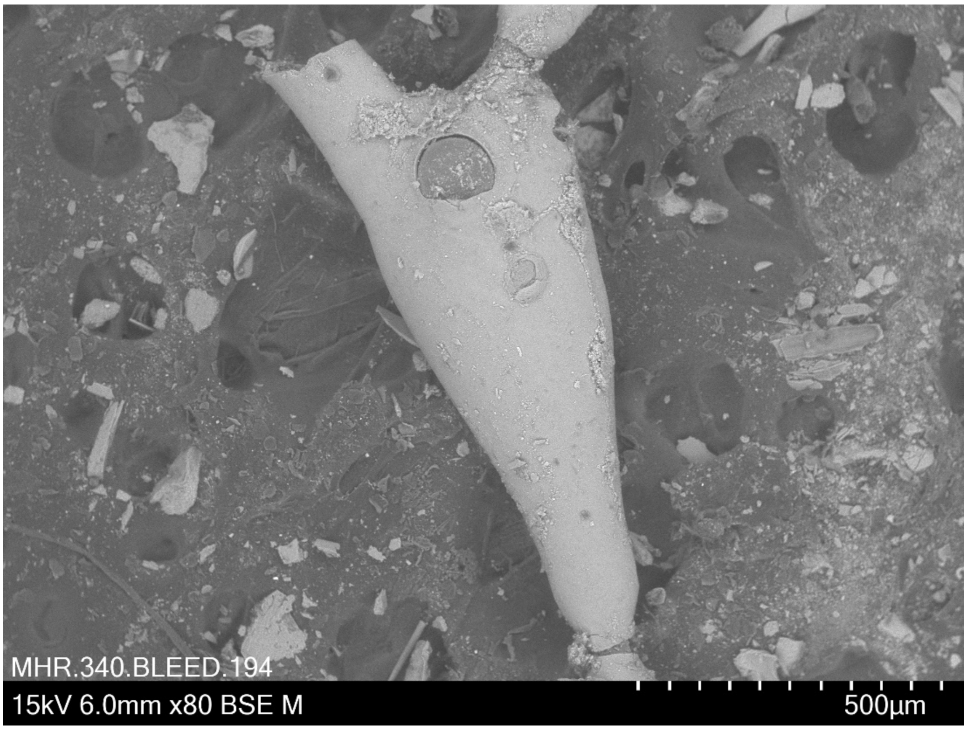


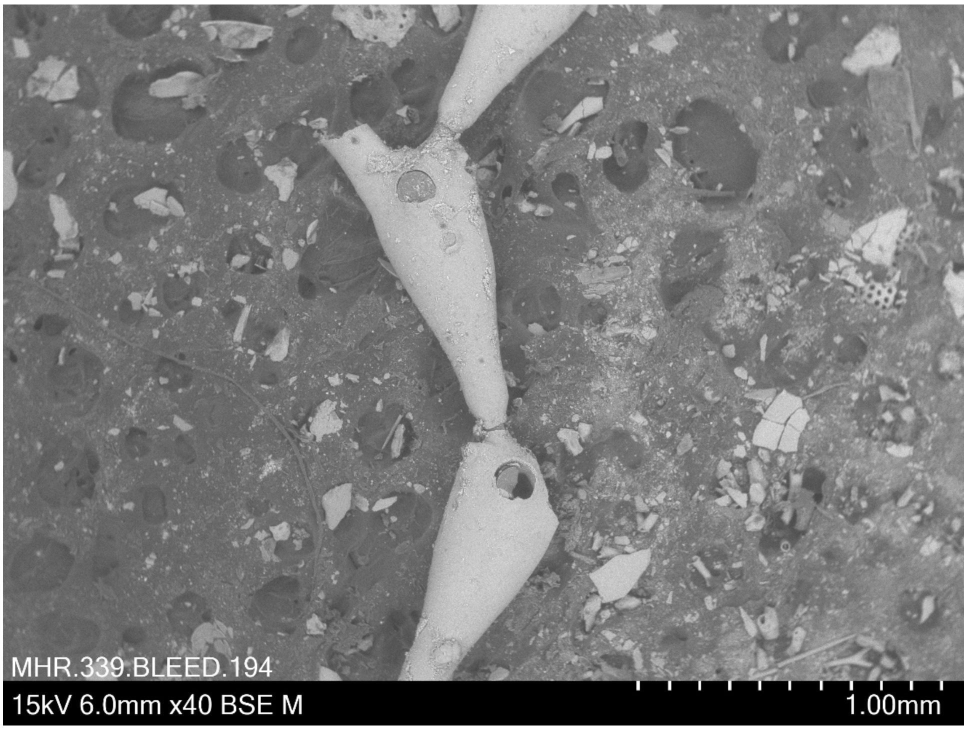


***Terminocella* n. sp. BLEED 194**
Coll. 2.6.2010 Pres. ethanol
Lat: -35.360, Long: 178.509, Depth: 1270m, New Zealand
Karma 2 TAN1007/56 NIWA 64553

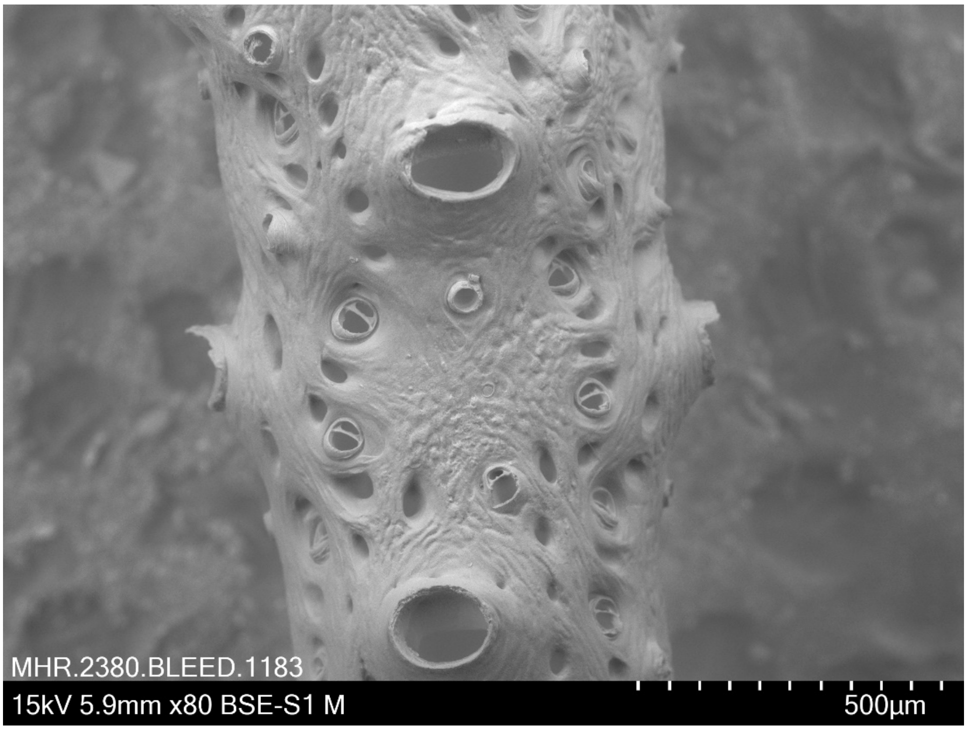


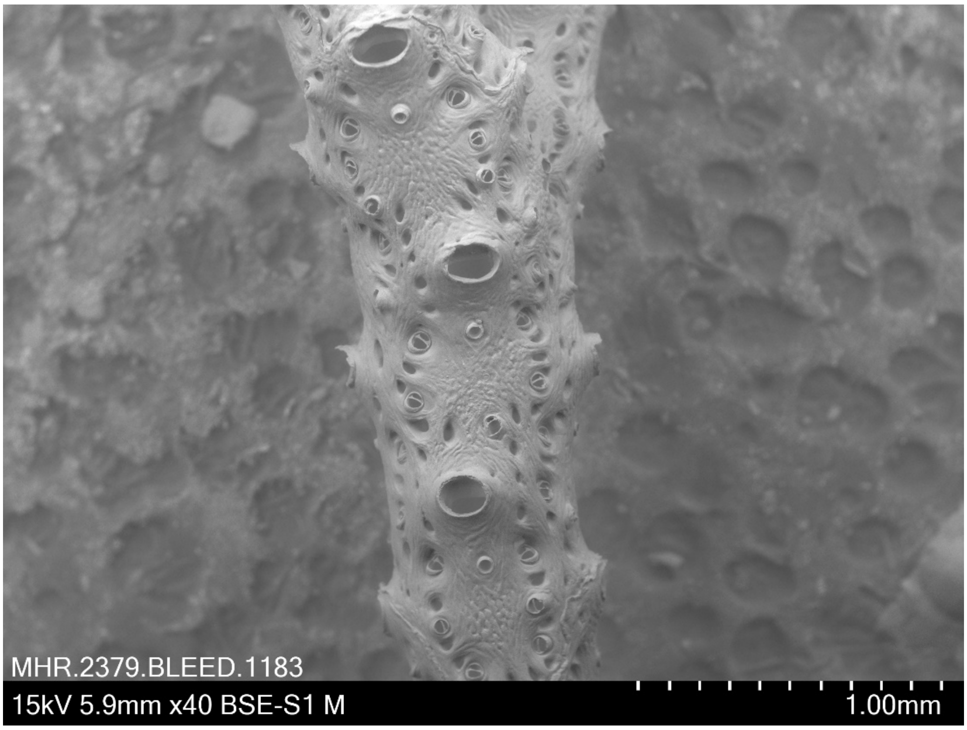


***Tessaradoma boreale* (Busk, 1860) BLEED 1183**
Pres. dried
Identified by I. Vigeland in 1970 (unregistered specimen from NHM Oslo)
Location: Bouge-strømmen mangen, 30 fathoms. No other metadata available.

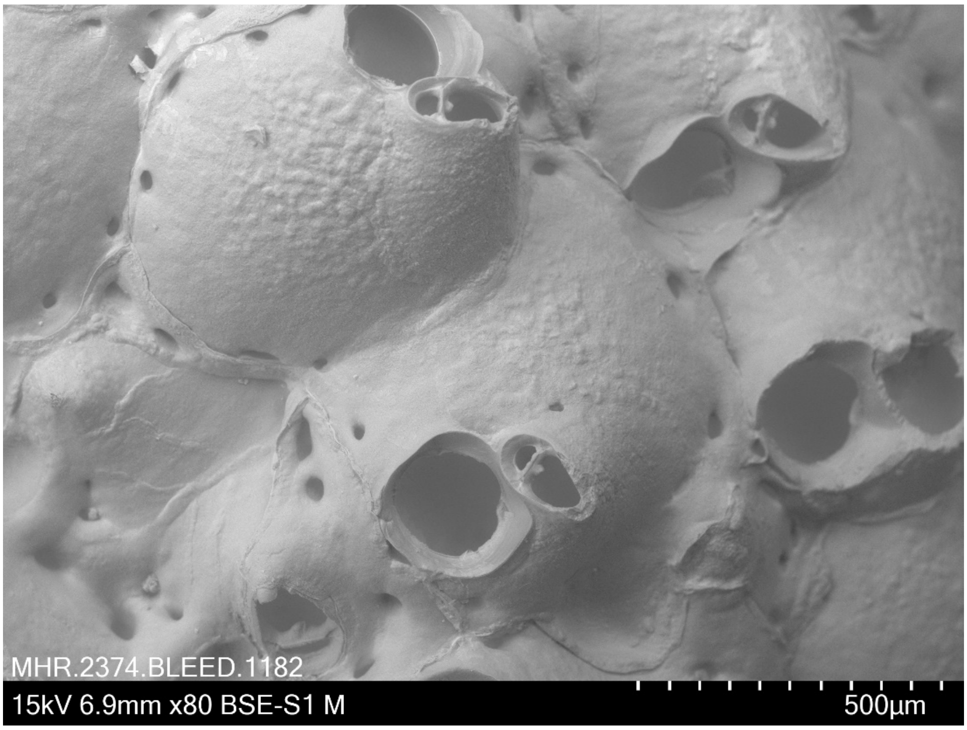


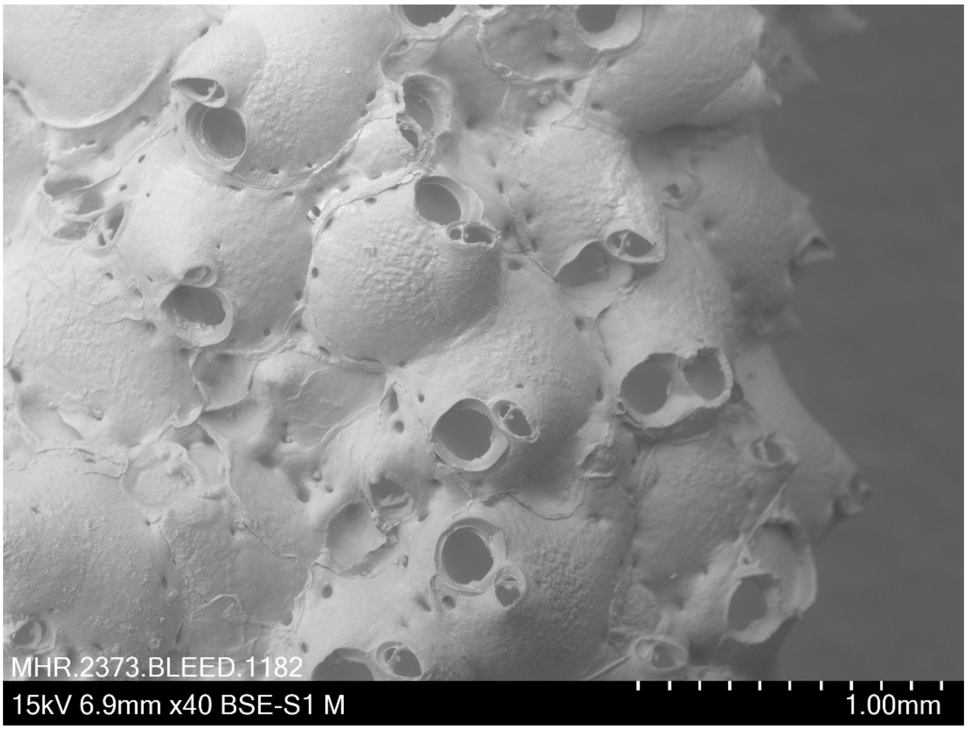


***Turbicellepora smitti* (Kluge, 1962) BLEED 1182**
Pres. dried
Identified by I. Vigeland in 1970 (unregistered specimen from NHM Oslo)
Location: Finnmark, Norway. No other metadata available.
